# Genomic signals of local adaptation and hybridization in Asian white birch

**DOI:** 10.1101/2022.07.01.498522

**Authors:** Gabriele Nocchi, Jing Wang, Long Yang, Junyi Ding, Ying Gao, Richard J. A. Buggs, Nian Wang

## Abstract

Disentangling the numerous processes that affect patterns of genome-wide diversity in widespread tree species has important implications for taxonomy, conservation, and forestry. Here, we investigate the population genomic structure of Asian white birch (*Betula platyphylla*) in China and seek to explain it in terms of hybridization, demography and adaptation. We generate whole genome sequence data from 83 individuals across the species range in China. Combining this with an existing dataset for 79 European and Russian white birches, we show a clear distinction between *B. pendula* and *B. platyphylla*, which have sometimes been lumped taxonomically. Genomic diversity of *B. platyphylla* in north-western China and Central Russia is affected greatly by hybridization with *B. pendula*. Excluding these hybridized populations, *B. platyphylla* in China has a linear distribution from north-eastern to south-western China, along the edge of the inland mountainous region. Within this distribution, three genetic clusters are found, which we model as long diverged with subsequent episodes of gene flow. Patterns of co-variation between allele frequencies and environmental variables in *B. platyphylla* suggest the role of natural selection in the distribution of diversity at 7,609 SNPs of which 3,767 were significantly differentiated among the genetic clusters. The putative adaptive SNPs are distributed throughout the genome and span 1,633 genic regions. Of these genic regions, 87 were previously identified as candidates for selective sweeps in Eurasian *B. pendula*. We use the 7,609 environmentally associated SNPs to estimate the risk of non-adaptedness for each sequenced *B. platyphylla* individual under a scenario of future climate change, highlighting areas where populations may be under future threat from rising temperatures.

## Introduction

Analysis of the population genomic diversity of species can help us to understand their past and predict their future. Past processes affecting present allele frequencies include hybridization, range contraction and expansion, population size changes, gene flow among populations, and natural selection. Understanding the effects of natural selection is particularly important for future predictions as the extent to which alleles are locally adaptive has important implications for estimating population fitness under climate change (Sork et al., 2013; Savolainen et al., 2007; Aitken et al. 2008). Covariance between allele frequencies and environmental variables may be signatures of local adaptation but only if the effects of other past processes are accounted for (Sork et al., 2013; Wang & Bradburd, 2014). Hence, we need to gain the fullest possible understanding of all past processes affecting allele frequencies in populations before we can predict patterns of diversity that may be beneficial in future climates.

Recently, the genetic basis of local adaptation has been investigated for certain non-model species using high-throughput sequencing, high-resolution environmental data, and novel analytical methods that seek to account for the effects of neutral processes (Eckert et al., 2010; Ellegren, 2014). Environmental association analysis (EAA) correlates abiotic data with genomic data, such as SNP allele frequencies (Manel and Holderegger 2013; Sork et al. 2013). Such studies pinpoint SNPs showing signals of elevated differentiation among environments and showing significant correlations with one or more environmental variables (Rellstab et al., 2015), which are putative candidates involved in local adaptation (Hoban et al., 2016). Using EAA, we can also predict the possible degree of maladaptation of local populations to future climate change (Borrell et al., 2019; Rellstab et al., 2021; Rellstab et al., 2016).

The future of forest trees under climate change is critical for carbon sequestration, ecosystem conservation and timber production. Forest trees with wide geographic distributions over spatially heterogeneous environments often show local adaptation with adaptive traits polygenically determined (Savolainen et al., 2007; Alberto et al., 2013). Due to their long generation times, forest trees may show adaptational lag to rapid climate change (Browne et al., 2019; Keenan, 2015). The genus *Betula* (birch) is a case in point, containing several tree species widespread in the Northern Hemisphere, comprising key components of temperate forests (Ashburner and McAllister, 2013; Wang et al., 2016; Wang et al., 2021). Understanding the genomic basis of adaptation in this genus can help to predict their potential to survive under rapid climate change, helping to manage species conservation (Borrell et al., 2019).

In this study, we selected Asian white birch (*Betula platyphylla*), which belongs to the subgenus *Betula*, to understand its past demographical histories, current local adaptation, and future potential to respond to climate change. Asian white birch is of great ecological and commercial value. It commonly occurs in eastern Asia including the Russian Far East, China, and Japan. In China, it is found in northwestern (NW), central (CE), northeastern (NE) and southwestern (SW) regions ranging over 26 latitudinal degrees (Chen & Lou, 2019; Ashburner and McAllister, 2013). It is a pioneer species, colonizing open habitats in grasslands, on roadsides and on the verge of forest (Ashburner and McAllister, 2013). In the south it can exist at altitudes of over 3,700 meters whereas in NW China it can grow by riversides in areas with much higher annual evaporation than precipitation.

Some taxonomists place *B. platyphylla* into the same species as silver birch, *B. pendula* (Ashburner & McAllister, 2013), which is widespread in Europe and Russia. Others see them as sister species (Chen et al., 2021; Wang et al., 2021; Salojärvi et al., 2017; Tsuda et al., 2017), with estimates of divergence times differing between 36,000 years (Tsuda et al., 2017) and 2.6 million years (Chen et al., 2021). Well assembled and annotated reference genomes now exist for both *B. pendula* (Salojärvi et al., 2017) and *B. platyphylla* (Chen et al., 2021) but the latter was not available when we commenced the present study. A large population genomic dataset is publicly available for *B. pendula*, analysis of which identified two genetic clusters with some inter-mixed populations and showed that variants of genes involved in photoperiodic responses were highly associated with local environmental adaptation (Salojärvi et al., 2017). The genetic structure of *B. platyphylla* has been previously investigated using ten microsatellite markers, showing five genetic clusters within its range, with substantial gene flow among them (Chen & Lou, 2019).

In this present study, we generate whole genome sequence data for *B. platyphylla* across its range in China to investigate its population genetic structure. We ask if *B. platyphylla* is differentiated from *B. pendula*, and if so, the extent to which hybridization among them affects *B. platyphylla* populations. We investigate the genetic structure within nonhybridized *B. platyphylla*, modelling how this has been affected by population divergence, population size change, and gene flow. We also model the environmental niche of *B. platyphylla* and how this may affect its range under future climates. We use EAA to identify SNP markers in *B. platyphylla* associated with environmental clines after controlling for population structure. We calculate the risk of non-adaptedness (RONA) (Rellstab et al., 2016) for each individual under a scenario of climate of 2080-2100. Together, this study generates new hypotheses regarding local adaptation in Asian white birch, which could be further tested in common garden trials or trans-genetic analyses in future studies.

## Materials and Methods

### Sampling

Samples of Asian white birch (*B. platyphylla*) were collected from 74 naturally occurring populations throughout its distribution in China between 2016 and 2019. Sample identifiers and GPS locations are presented in Table S1. Dried cambial tissues were used for DNA extraction following a modified CTAB protocol (Wang et al., 2013). One or two samples per population were selected for resequencing, resulting in a total of 83 samples. Extracted DNA was assessed with 1% agarose gel and sent to BerryGenomics (Beijing, China) for library preparation and whole genome re-sequencing. Sequencing was performed on the Illumina NovaSeq6000 platform with 150 bp paired end read sequencing with approximate 11Gbp data obtained for each sample (average depth of ∼25x). We also downloaded genomic read data from Salojarvi et al. (2017) for 78 *B. pendula* individuals from most of this species’ European and Asian geographic range and one *B. platyphylla* individual from Russia.

### Trimming and SNP filtering

The raw data was trimmed using Trimmomatic (Bolger et al., 2014) in paired-end mode with a required quality of 30. Reads shorter than 90 bp were discarded. Clean reads of the sequenced samples were aligned to a genome assembly of *B. pendula* (Salojärvi et al., 2017) using BWA-MEM v.0.7.17-r1188 algorithm in BWA (v0.7.17) with default parameters (Li & Durbin, 2009). Reads with non-specific matches were discarded. Alignments were converted from sequence alignment map (SAM) format to sorted, indexed binary alignment map (BAM) files (SAMtools v1.8) (Li et al., 2009). The MarkDuplicates tool from the Genome Analysis Tool Kit (GATK) (v 4.1.4.) was used to mark duplicates (DePristo et al., 2011; McKenna et al., 2010). We filtered for biallelic SNPs genotyped in all individuals with minor allele frequency greater than 0.05, quality by depth greater than 2, fisher strand test less than 60, root mean square mapping quality greater than 40, mapping quality rank sum test greater than -12, read position rank sum test greater than -8 and strand odds ratio less than 3. We pruned the filtered SNP set by linkage disequilibrium (r^2^ > 0.4) using the “indep-pairphase” function of Plink (Chang et al., 2015) in windows of 50 and step of 5. We term this set of SNPs the China dataset.

Using the same method to the above we also assembled a second SNP dataset that included genomic read data from Salojarvi et al. (2017) for 78 *B. pendula* individuals from most of this species’ European and Asian geographic range, and one *B. platyphylla* individual deliberately included from Russia. We term this set of SNPs the Eurasian dataset.

### Population structure

We used *fastSTRUCTURE* (Raj, Stephens, & Pritchard, 2014) on the China dataset and also on the Eurasian dataset to assign individuals to populations. We used a simple prior, 5-fold cross-validation and number of ancestral populations (K) from one to ten. We used the python function *chooseK*, which is the recommended tool for model selection in *fastSTRUCTURE* (Raj, Stephens & Pritchard, 2014), and the cross-validation profile, which identifies the model complexity for which the prediction error is minimized, to assess the models generated. The function *chooseK* suggests two values for K: one that maximizes the log-marginal likelihood lower bound (LLBO) of the dataset (K*C) and captures strong structure, and another one (Køc) which reports the model components that have a cumulative ancestry contribution of at least 99% and it is aimed at finding additional weak underlying structure (Raj, Stephens, & Pritchard, 2014). Individual trees were assigned to populations according to the value of the admixture coefficient (q) computed by *fastSTRUCTURE* at the chosen model complexity: q ≥ 0.9 to belong to a population, while individuals with q < 0.9 were classified as admixed.

On the basis of the *fastSTRUCTURE* analysis of the Eurasian dataset, we excluded from the China dataset any individuals that had over 10% assignment to *B. pendula* and ran *fastSTRUCTURE* only on Chinese *B. platyphylla* based on 1,387,994 SNPs (after filtering for missing genotypes). Furthermore, we used the program sparse non-negative matrix factorization (*snmf*), part of LEA package (Caye et al., 2019), to perform a population differentiation test and identify F_st_ outlier SNPs between the clusters identified for *B. platyphylla* in China. *snmf* computes fixation also in the presence of admixed individuals in the sample (Martins et al., 2016) and performs a test similar to other F_st_-outlier approaches (Lotterhos & Whitlock, 2014). Multiple testing was controlled by converting the p-values computed by *snmf* into q-values with the R package “qvalue” (Storey et al., 2015). Q-values represent the expected false discovery rate (FDR) associated with a given p-value (Storey and Tibshirani 2003). Finally, we selected candidate outlier SNPs with FDR < 1%.

Principal component analysis (PCA) was performed on the Eurasian dataset and the Chinese *B. platyphylla* dataset using *Plink v2*.*0* (Chang et al., 2015).

### Demographic histories

We inferred the demographic histories of the Chinese *B. platyphylla* dataset (excluding NW China) for populations specified with K=3 and K=2 in the STRUCTURE analyses above (the relative merits of these divisions reported in the Results and Discussion sections below) using a coalescent simulation-based method implemented in fastsimcoal2 v 2.6.0.3 (Excoffier et al. 2013). The pairwise two-dimensional joint SFSs (2D-SFSs) among the populations were constructed from the posterior probabilities of sample allele frequencies using ANGSD v 0.934 (Korneliussen et al., 2014). A total of 28 models were evaluated for K=3 assuming divergence with or without gene flow and a constant or changed population size after divergence (Figure S1; Table S2). Fourteen models specified that NE diverged first, followed by the split of the CE and SW populations and 14 models specified the CE population as of hybrid origin between the NE and SW populations (Figure S1). In addition, we ran nine models for K=2 (Figure S2). The global maximum likelihood estimates for all demographic parameters under each model were obtained from 60 independent runs, with 100,000 coalescent simulations per likelihood estimates and 40 cycles of the likelihood maximization algorithm. The models were compared based on the maximum value of likelihood over the 60 independent runs using the Akaike’s weight. The model with the maximum Akaike’s weight value was chosen as the optimal one. Confidence intervals were generated by performing parametric bootstrapping with 100 bootstrap replicates, and with 30 independent runs in each bootstrap. We used a mutation rate of 9.5 × 10^−9^ per site per year (Salojärvi et al. 2017) and a generation time of 15 years to convert the coalescent scaled time into a speculative time scale in years. The best-fitting model was validated via comparing the observed SFS to the predicted SFS based on the model (Figure S3).

In addition, we inferred the historical change in effective population sizes using Multiple Sequentially Markovian Coalescent approach (MSMC v2.0.0) (Schiffels & Durbin, 2014) at K=3 for each of the three populations, respectively. For each population, we randomly chosen seven individuals and applied simple trio-phasing on any four individuals (eight haplotypes) using scripts provided in the msmc-tools github repository (https://github.com/stschiff/msmc-tools). This resulted in 35 different configurations for the MSMC analysis. We ran MSMC on each of the 35 configurations and estimated medians and standard deviations of effective population sizes changes across time.

### Introgression test

We used the ABBA-BABA tests to detect signals of hybridization (Green et al. 2010; Durand et al. 2011). We based these tests on all individuals from the SW, NW and CE populations, ten individuals from the NE population and ten *B. pendula* individuals from Salojarvi et al. (2017). F1 hybrids detected by fastSTRUCTURE with a Q-score of approximately 0.5 were not included. We detected asymmetries in the frequency of shared and derived variants (Green et al. 2010; Durand et al. 2011) using different combinations of three populations, using *Betula occidentalis* from Salojärvi et al. (2017) as an outgroup for all tests. The D and Z statistic were calculated for each combination using function “Dtrios” in Dsuite 0.4-r38 (Malinsky et al., 2021). A block jack-knifing was performed to obtain an associated p value for the test of whether D statistic deviates significantly from zero. An average Z score larger than 3.09 (equal to a p value < 0.001 of D statistic) was regarded as a significant signal of introgression.

### Population diversity and differentiation

We estimated linkage disequilibrium decay along the *B. platyphylla* genome with the tool *PopLDdecay* (Zhang et al., 2018) separately for the different populations identified for *B. platyphylla* in China by *fastSTRUCTURE* at K = 3 (Figure 3). For this calculation we used the SNP set prior to linkage disequilibrium pruning but filtered to retain SNPs with minor allele frequency > 0.05. After excluding individuals admixed between the three populations (admixture coefficient < 0.9 in *fastSTRUCTURE* at K = 3), Weir and Cockerham F_st_ (Weir & Cockerham, 1984) was calculated between the three *B. platyphylla* populations on a per SNP site basis with the *vcftools* function *weir-fst-pop* (Danecek et al., 2011). We calculated nucleotide diversity π (Nei & Li, 1979) within each population using the *vcftools* (Danecek et al., 2011) function *window*-*pi* in non-overlapping windows of 5 kb. For the π calculation we used the SNPs set prior to minor allele frequency filtering and linkage disequilibrium pruning (28,058,885 SNPs in total).

### Environmental niche modelling

We used environmental niche modelling (ENM) to characterize the present habitat and predict the future distribution of non-hybrid *B. platyphylla* in China and to identify the climatic variables influencing this species distribution. ENM analysis was performed by using the samples’ geographic locations in this study and observation records of this species from 1970 to 2010 in China, sourced from the Global Biodiversity Information Facility (http://www.gbif.org). To mitigate the effects of spatial autocorrelation, we removed records within 0.2 degrees (approximately 22-25 km) of one another, which resulted in a total of 138 presence locations: 66 records were *B. platyphylla* from the 83 individuals sampled (excluding admixed *B. pendula* -*B. platyphylla* individuals) and another 72 records were from GBIF (excluding NW China as all the trees that we had sampled there were hybrids – see Results). We downloaded 19 bioclimatic variables related to temperature and precipitation from the WorldClim database (www.worldclim.org) at 1km resolution (Hijmanset al., 2005) for the period 1970-2000 representing “current climate”, as this is the estimated time period of establishment of the sampled trees and also corresponds to the time that most GBIF observations included in the model were recorded. We downloaded elevation data at the same resolution and used it to compute slope, aspect and four additional terrain related characteristics (topographic position index, terrain ruggedness index, terrain roughness and water flow direction) using the *raster* R package (Hijmans & Etten, 2012). In order to avoid overfitting, we selected environmental variables with Pearson’s correlation coefficient < 0.7, preferring annual rather than monthly or quarterly values, which resulted in 11 variables retained for the ENM (Table S4, Figure S4-5). We assembled eight further datasets with the same 11 variables under four Shared Socioeconomic Pathways (SSPs) defined by the Intergovernmental Panel on Climate Change sixth Assessment (Masson-Delmotte et al., 2021) at each of two future time points (2041–2060 and 2081–2100). The ENMs were generated using Maxent (Phillips et al., 2006) with 50 subsampled replicate 5000 iterations runs with 20% of observations left-out for cross-validation. Variables’ importance was assessed with jack-knife tests and multiple models were generated and evaluated using a test of omission rate and area under the receiver operating characteristic curve (AUC). An environment suitability threshold was defined by “maximum training sensitivity plus specificity,” which optimizes the trade-off between commission and omission errors (Liu et al., 2016).

### Genome environment association analysis: identification of adaptive SNPs

We used *LFMM2* (Caye et al., 2019) to test for associations between environmental variables and SNP allele frequencies in 71 Chinese non-hybrid *B. platyphylla* individuals. *LFMM2* performs multivariate linear regressions to evaluate the association between a response matrix, corresponding to SNP frequencies of individuals, and a matrix of environmental variables. It combines the environmental fixed effects with latent effects, which are unobserved confounding effects due to population structure (Frichot et al., 2013; Caye et al., 2019). *LFMM2* has a two steps approach: first it estimates the latent factors and then tests for the genotype-environment associations. Each SNP locus is tested separately using the latent scores as covariates to control for demographic history and population structure. The null hypothesis is that the environmental variables have no effects on the SNP frequency, which is tested using a student distribution with n−K−1 degrees of freedom (Caye et al., 2019). *LFMM2* requires to specify the number (K) of latent factors to include in the model which was chosen on the basis of *fastSTRUCTURE* cross-validation schemes and a Tracy-Widom test on the PCA eigenvalues. We chose the more conservative K = 3 (see Results section) to run *LFMM2*, including all the 11 uncorrelated environmental variables used in the ENM (Table S4). Multiple testing was controlled by converting the p-values computed by *LFMM2* into q-values with the R package “qvalue” (Storey et al., 2015). Finally, we selected candidate adaptive SNPs with FDR < 1%.

### Characterization of candidate adaptive SNPs

We performed two further principal component analyses of the Chinese *B. platyphylla* samples with *Plink* (Chang et al., 2015): one including only the putative adaptive SNPs (detected by *LFMM2* (K = 3) with FDR < 1%) and another only including a similar number of putatively neutral SNPs. The neutral set of 7,500 SNPs was generated by selecting SNP loci at random from those that reported a q-value above 0.4 across all environmental variables in the *LFMM2* analysis. Weir and Cockerham F_st_ between the three *B. platyphylla* populations was re-computed based on the adaptive and neutral SNP sets with the *vcftools* function *weir-fst-pop* (Danecek et al., 2011) after excluding admixed individuals (admixture coefficient < 0.9 in *fastSTRUCTURE* at K = 3).

In order to functionally annotate our putative adaptive SNPs, we performed a gene ontology (GO) annotation of the whole *B. pendula* reference gene set provided in Salojärvi et al. (2017), retrieving GO terms associated with the coding regions (Ashburner et al., 2000) using *Omics Box* (Conesa & Götz, 2008). We used *Omics Box* default functional annotation workflow which incorporates BLAST, run on the Viridiplantae non-redundant database, and Interproscan (Zdobnov & Apweiler, 2001). We then identified the genic regions overlapping our candidate adaptive SNPs with bedtools v2.28.0 (Quinlan & Hall, 2010) and performed a functional enrichment analysis of this adaptive gene set against the fully annotated reference gene set using Fisher’s exact test in *Omics Box* (Conesa & Götz, 2008). We applied multiple testing corrections (FDR < 5%) and reduced the resulting significantly enriched GOs to the most specific term in the hierarchy for all three ontology levels: biological process, molecular function, and cellular component (Ashburner et al., 2000).

### Adaption to future climate

We carried out risk of non-adaptedness (RONA) analysis using a slight modification of the method of Rellstab et al. (2016), implemented in the software *pyRona* (Pina-Martins et al., 2018). RONA represents an estimate of the average change in allele frequencies at climate associated SNP loci required in a population to cope with future climatic conditions and is calculated separately for each environmental variable (Rellstab et al., 2016). RONA uses simple linear regressions of the selected current climate environmental variable and the alternative allele frequencies of the candidate loci identified in genome-environment association analyses (Rellstab et al., 2016). The regression coefficients of the environmentally associated loci are then used to predict the allele frequencies of the adaptive loci at a future climate value for the chosen environmental variable (Rellstab et al., 2016). The main improvement in the implementation by Pina-Martins et al. (2018) is that the RONA for an environmental variable is given by the weighted mean RONA of all relevant SNP loci for that environmental variable, weighted by the r^2^ value of each locus regression, while the original method (Rellstab et al., 2016) uses unweighted means. We computed RONA on an individual sample basis for the seven uncorrelated environmental variables that are expected to change in the future, therefore excluding elevation and its derived measures. We used as future climate the profile ssp370 for 2080-2100 (Figure 4C). For each environmental variable, RONA was calculated by using the candidate adaptive SNP loci identified with *LFMM2* (FDR < 1%). For each individual tree we then calculated weighted mean RONA, giving each environmental variable RONA a weight equivalent to the contribution reported by that variable in the ENM. Furthermore, we reported the maximum RONA for each individual, out of the seven variables for which it was calculated.

## Results

### SNP discovery

After filtering, the number of reads retained for each sample in the China dataset that we generated ranged between 66,016,635 and 134,129,672 and between 91.5% and 96.7% of these reads per individual were mapped to the reference genome (Table S1). SNPs filtering of the 83 individuals resulted in 1,497,547 unlinked (r^2^ < 0.4) SNPs. After filtering, the number of reads retained for each sample in the Salojarvi et al. (2017) dataset ranged between 16,506,088 and 237,845,424 and between 82.9% and 97.2% of these reads per individual were mapped to the reference genome. The combined Eurasian dataset resulted in 278,717 SNPs after filtering for missing genotypes. When the China dataset was restricted to pure *B. platyphylla* (71 individuals in SW, CE, and NE China), we were left with 1,387,994 SNPs.

### Population structure, diversity, and differentiation

For the China dataset *fastSTRUCTURE* analysis with K = 2 separated the samples in the NW population (Xinjiang) from the other populations. At K = 3, the SW and NE populations were further separated and at K = 4 the CE population was revealed (Figure 1). The model that maximized the log-marginal likelihood lower bound (LLBO) of the data and that best explained additional weak underlying structure was K = 3, as suggested by the function *“chooseK”* (Figure S6).

**Figure 1.**
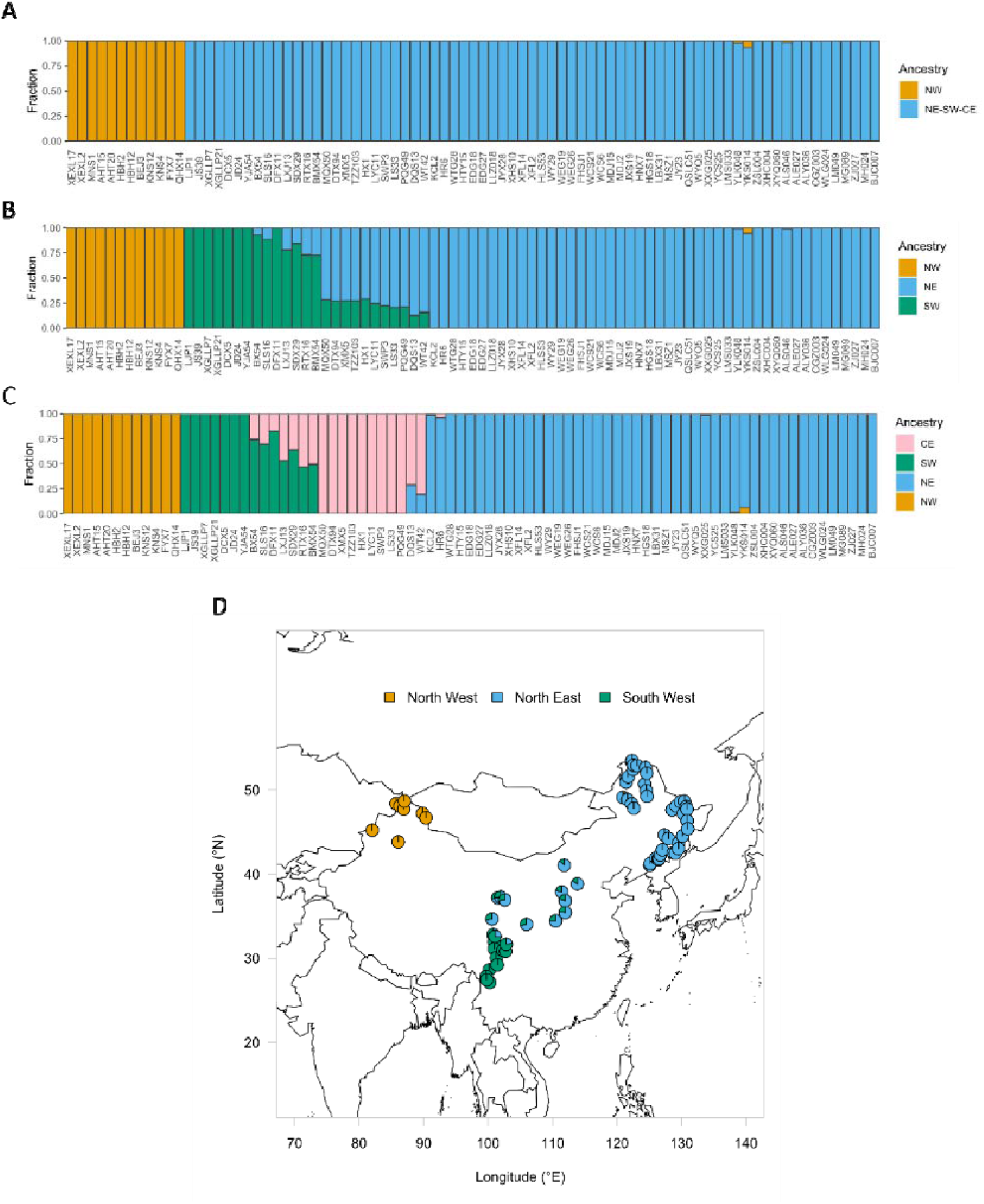
*fastSTRUCTURE* results for the China dataset, ordered by longitude. **A)** Bar plot of *fastSTRUCTURE* results of 83 sampled birch individuals, based on 1,497,547 unlinked SNPs (r^2^ < 0.4) at K = 2, **B)** K=3 and **C)** K =4. **D)** Map of the individuals showing their genetic ancestry composition according to *fastSTRUCTURE* at the ideal model complexity, K = 3.

For the Eurasian dataset at K = 2, one cluster contained the *B. pendula* individuals from Salojarvi et al. (2017) and the other cluster contained 71 of the 83 individuals that we sequenced from China and the Russian *B. platyphylla* individual from Salojarvi et al. (2019) (Figure 2). The 12 individuals we sampled in the NW population were shown as hybrids between *B. pendula* and *B. platyphylla* (Figure 2A). At K = 3, the pattern was similar but a SW *B. platyphylla* clustered separately from a NE *B. platyphylla* cluster (Figure 2B). At K = 4, the only difference was that Irish and two Finnish *B. pendula* samples showed an unknown component (Figure 2C). The *fastSTRUCTURE* model that maximized the log-marginal likelihood lower bound (LLBO) of the data and that best explained additional weak underlying structure was K = 4, as suggested by the function choose K (Figure S7). The LLBO curve and the cross-validation profile showed that the marginal likelihood plateaued at K = 2, and so did the prediction error, which stayed within 1 standard error from K = 2 with added complexity (Figure S7A-B). These separations suggested by *fastSTRUCTURE* were reflected in the PCA (Figure S8).

**Figure 2.**
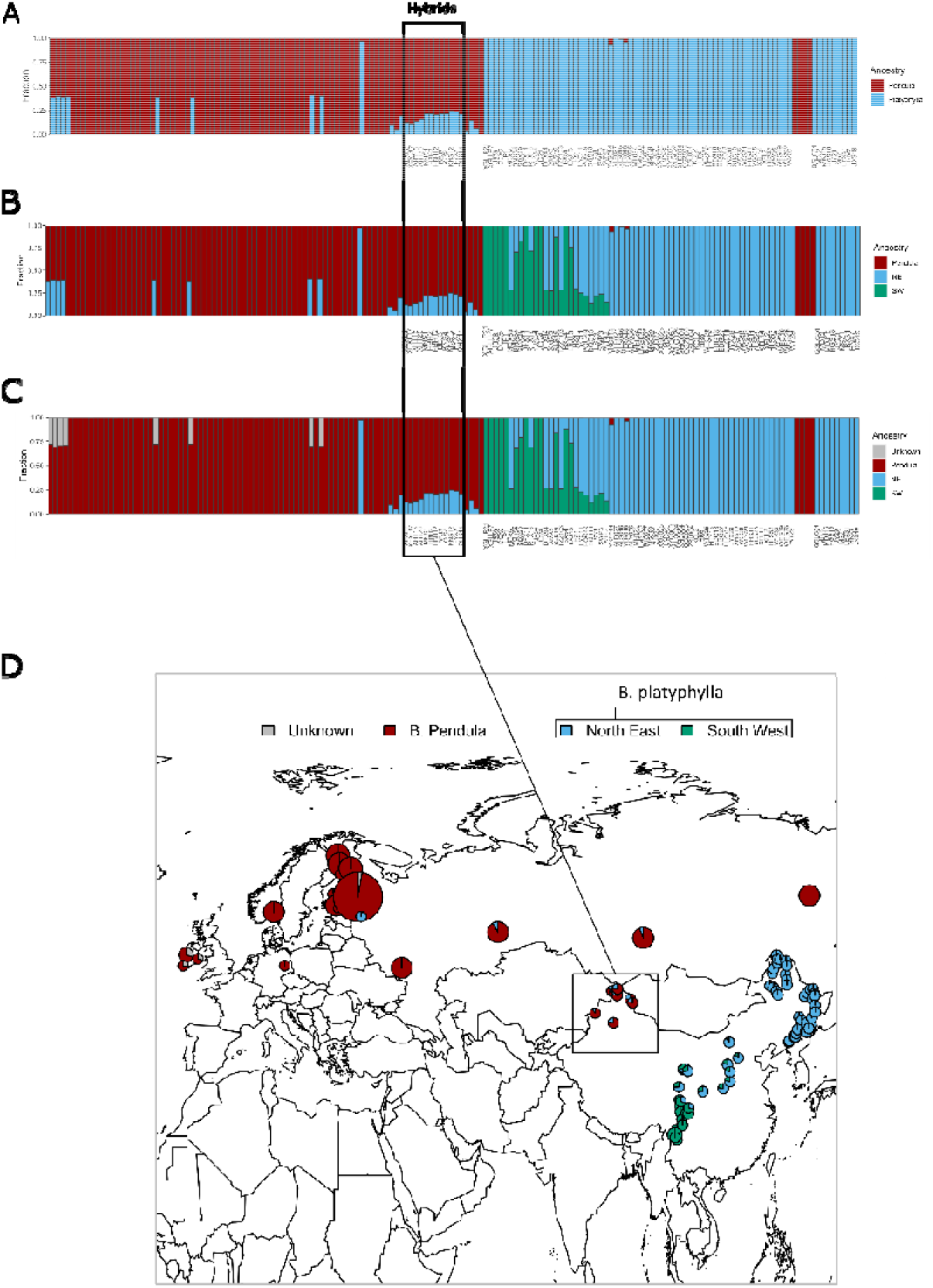
*fastSTRUCTURE* results for the Eurasian dataset, ordered by longitude. The size of the pies is proportional to the number of samples. The labelled bars correspond to individuals sampled in this study, while the unlabelled bars are the samples added from Salojarvi et al. (2017) **A)** Bar plot of *fastSTRUCTURE* results of 162 *B. pendula* and *B. platyphylla* individuals, based on 278,717 unlinked SNPs (r^2^ < 0.4) at K = 2, **B)** K=3 and **C)** K =4. **D)** Map of the individuals showing their genetic ancestry composition according to *fastSTRUCTURE* at K = 4.

In *fastSTRUCTURE* restricted to the 71 non-hybrid *B. platyphylla* individuals in China, the model that maximized the log-marginal likelihood lower bound (LLBO) of the data was K = 2 and that best explains additional weak underlying structure was K = 3, as suggested by the function *“chooseK”* (Figure S9). The *fastSTRUCTURE* cross-validation score profile showed that the prediction error is lowest at K = 3 (Figure S9B). At K = 2 the NE population was separated from the SW population and the individuals in the CE population appeared admixed between these two populations, while at K = 3 the CE population was distinct (Figure 3A and 3B).

**Figure 3.**
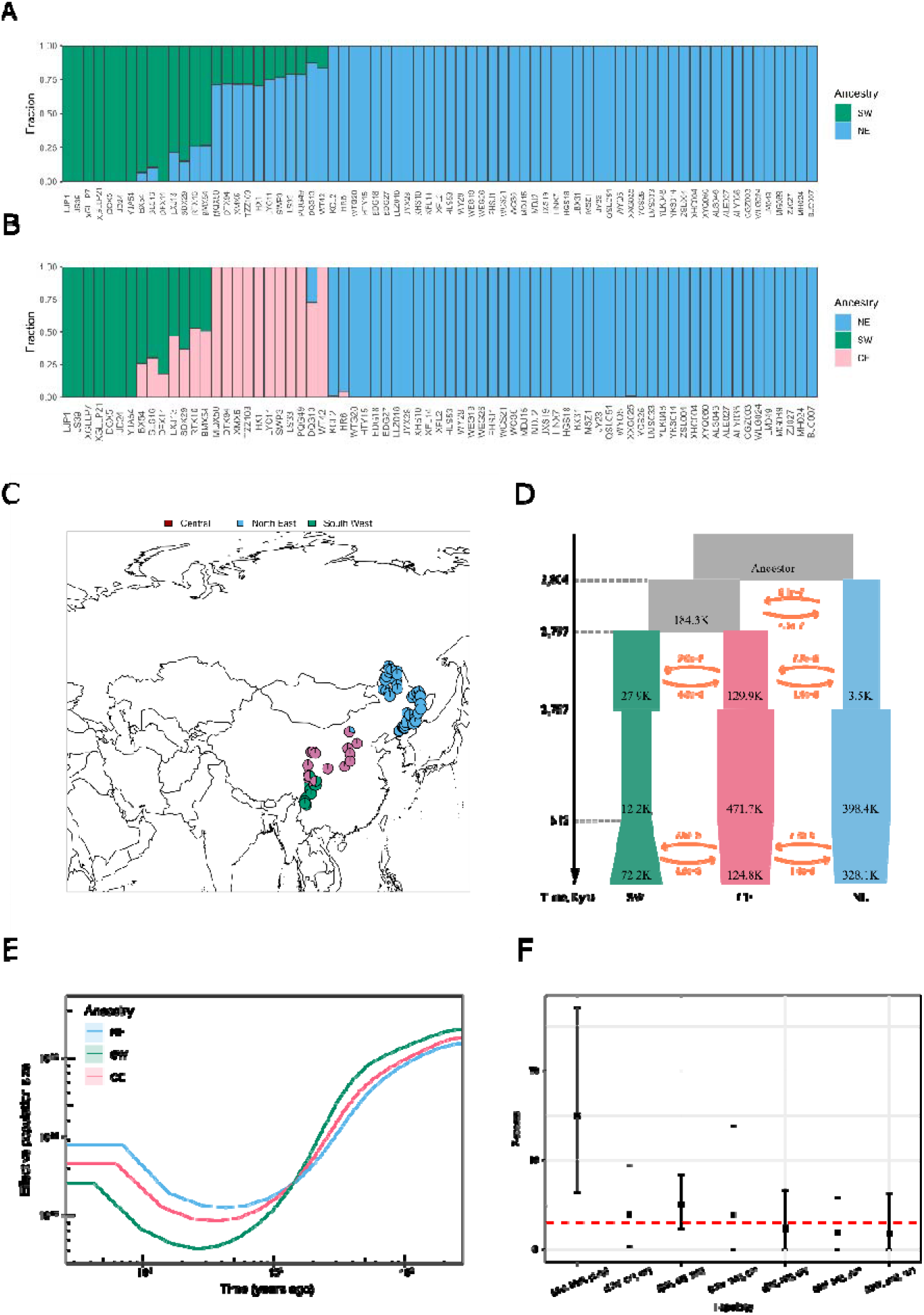
*fastSTRUCTURE, MSMC* and *fastsimcoal* results for Chinese *B. platyphylla*, ordered by longitude. **A)** Bar plot of *fastSTRUCTURE* results of 71 *B. platyphylla* individuals, based on 1,387,994 unlinked SNPs (r^2^ < 0.4) at K = 2, and **B)** K=3. **C)** Map of the sampled individuals showing their genetic ancestry composition according to *fastSTRUCTURE* at K = 3. **D)** A demographic scenario of the NE, CE and SW lineages modelled using fastsimcoal2. Shown in light grey are the ancestral populations. The arrows show the per generation migration rate (m) between the three lineages. **E)** Historical effective population size of the three populations estimated by MSMC. Solid lines and shading represent medians and ±standard deviation calculated across pairs of haplotypes, respectively. **F)** The results of “ABBA-BABA” tests. The red line denotes topologies with significant signals of introgression, which are indicated by a mean Z scores > 3.09. EU and platy represent *B. pendula* and *B. platyphylla*, respectively.

In PCA of the 71 pure *B. platyphylla* individuals, PC1 explained ∼33% of the total variance and separated the NW, CE, and SW populations (Figure S10). PC2 (15% of total variance) also separated the three populations while PC3 (8% of total variance) scattered the individuals of the NE population. The Tracy-Widom test identified three significant principal components (p <.05) and the explained variance levelled-off after PC3 (Figure S10D).

Due to the differentiation between the individuals located in central China and those in the North-East shown by the PCA (Figure S10) and the recommendation by *fastSTRUCTURE* cross-validation scheme (Figure S9B), we assigned individuals to populations according to the admixture coefficients computed with *fastSTRUCTURE* at K = 3. In total, ten individuals were assigned to the central population (CE), seven to the south-western population (SW), 46 to the north-eastern population (NE) and the remaining eight individuals were classified as admixed (Figure 3B-C). Based on this assignment, *snmf* identified 17,218 outlier SNPs with FDR < 1% (Figure S11).

Linkage disequilibrium decays rapidly in *B. platyphylla* populations and reaches background levels at approximately 50 kb (Figure S12). Assessment of pairwise F_st_ between populations showed that the highest levels of differentiation were between the NE and SW populations (mean F_st_ = 0.10) or the CE population and SW populations (mean F_st_ = 0.11) (Figure S13). On the other hand, the CE and NE populations reported significantly lower mean pairwise F_st_ (0.05) (Figure S13).

Nucleotide diversity π in windows of 5kb showed a slight increase in populations from south to north geographically, reporting a mean of 0.0057 (∼0.6%) in the SW population, 0.0075 (∼0.7%) in the CE population and 0.0083 (∼0.8%) in the NE population (Figure S14-15).

### Species population size and separation history

Assuming that there are three populations *of B. platyphylla* in NE, CE and SW China (K=3), and the NE population was the first to diverge, the best-supported fastsimcoal2 model (Figure 3D; Table S2) suggested that the NE population diverged from the common ancestral population of CE and SW at 2.804 Mya (95% CI: 2.892–3.506 Mya) (Figure 3D). The NE population initially had a small population size. The CE and SW populations diverged 2.797 Mya (95% CI: 2.294–2.768 Mya; Figure 3D). The SW population underwent a slight bottleneck at 2.787 Mya (95% CI: 2.129–2.489 Mya) and experienced a slight population expansion at 0.812 Mya (95% CI: 0.477–0.706 Mya) (Figure 3D). The estimated present day effective population sizes (Ne) are 3.3 x 10^5^ (95% CI: 3.0–3.4 × 10^5^) for NE, 1.2 × 10^5^ (95% CI: 1.6–2.2 × 10^5^) for CE and 0.7 × 10^5^ (95% CI: 1.6–2.2 × 10^5^) for SW. The best-supported model detected low levels of ancient gene flow between the populations soon after their divergence, no gene flow between 2.797 Mya and 0.812 Mya, and recent gene flow between the NE and CE populations and between the SW and CE populations (Figure 3D). We also ran fastsimcoal2 models assuming K=2 and the best model suggested an older initial divergence than we found with K=3 and an early bottleneck in the SW population only. Patterns of gene flow and current Ne were similar to those of the K=3 model.

(Figure S2, Table S3).

MSMC results suggested that the three populations experienced severe bottlenecks (2.51-2.89 million years ago) followed by population expansion during the period of 26.1-42.6 thousand years ago (Figure 3E). The NE lineage experienced the least population reduction (Ne, 11235.88 ± 204.75) whereas the SW lineage underwent the most severe population decline (Ne, 6152.70 ± 265.12) (Figure 3E).

### Hybridization between B. platyphyllya and B. pendula

The ABBA-BABA tests showed a high degree of introgression from *B. pendula* into the NW *B. platyphylla* population (mean D-statistic = 0.17, mean Z-score = 15.02), which was also identified as having mixed ancestry in the fastSTRUCTURE analysis (Figure 2; Figure 3F). The ABBA-BABA results also indicatpotential gene flow between the NE, CE and SW populations (Figure 3F).

### Environmental niche modelling

Environmental niche models were built based on eleven uncorrelated environmental variables (Figure S4-5; Table S4) and were based on our *B. platyphylla* samples’ locations and GBIF observations of this species (Figure 4).

**Figure 4.**
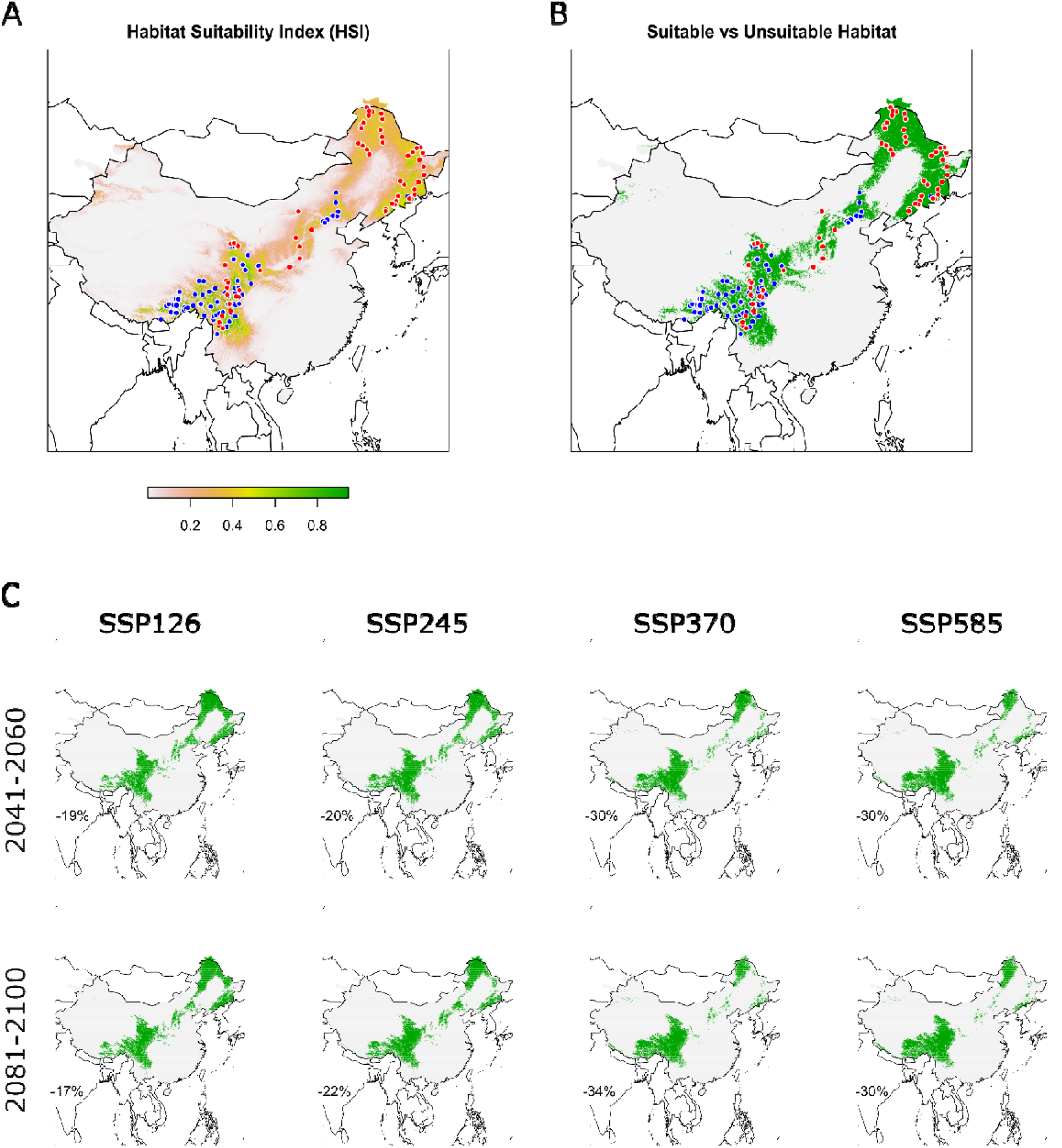
Environmental niche model (ENM) of *B. platyphylla* in China. Red points represent the individuals sampled in this study included in the model, blue points represent observations downloaded from the GBIF (http://www.gbif.org). **A)** Present time ENM showing the habitat suitability index (HSI) throughout China using a coloured scale from 0 (white) to 1 (green). **B)** Binomial representation of the HSI model, using the maximum training sensitivity plus specificity threshold of 0.2. **C)** Binomial HSI projections (> 0.2) of *B. platyphylla* under four different future climate scenarios (columns: ssp126, ssp245, ssp370 and ssp585) at two different time points (rows: 2041-2060 and 2080-2100). The percentage in each plot shows the decrease in suitable environment area compared with the current habitat.

Jack-knife tests suggested that the variables “aspect” and “water-flow direction” did not play a relevant contribution either singly or in combination, so they were excluded (Figure S16-18). The final Maxent model with the nine remaining variables reported high mean test AUC (0.913 ± 0.017) and low mean test omission rate (0.13, p <.001) at a maximum training sensitivity plus specificity logistic threshold of 0.2 (Figure 4A-B; Figure S19-20). Two variables, “annual precipitation” and “mean temperature of wettest quarter”, together contributed to > 50% of the predicting performance of the model, defined as the increase in regularized gain added (or subtracted) to the contribution of the corresponding variable, over each model iteration (Table S5).

Response curves for the final Maxent model are available in Supporting Information (Figures S28-S36). Future projections show an overall contraction of the *B. platyphylla* habitat throughout China, particularly noticeable in central China and in the north-east (Figure 4C). Currently suitable environments located at higher elevations seem to contract less in the future (Figure 4C). The predicted habitat reduces between 19 and 30 % in 2040-2060, and between 17 and 34 % in 2080-2100 (Figure 4C), compared to the present. In the most adverse scenario (ssp370), *B. platyphylla* habitat in China may be reduced by approximately a third of its current extent by 2080-2100 (Figure 4C) and the NE and SW populations will be spatially separated by a large uninhabitable region.

### Genome-environment association analysis: identification and characterization of putatively adaptive SNPs

We choose a q-value threshold of 0.01 (FDR < 1%) to select candidate associations in the LFMM2 analysis on 1,387,994 SNPs (r^2^ < 0.4) restricted to the 71 non-hybrid *B. platyphylla* individuals identified with *fastSTRUCTURE* (i.e. excluding the individuals in NW China).

In total, 7,609 SNPs showed significant associations with one or more environmental variables in *LFMM2* (K = 3) using our criteria (Figure S21), for a total of 11,304 associations. In more detail: 4,643 SNP were associated to one environmental variable, 2,424 SNPs to two variables, 384 SNPs to three variables, 133 SNPs to four variables, 21 SNPs to five variables and four SNPs to six variables. Of the 7,609 putative adaptive SNPs identified with *LFMM2*, 3,767 were shown to be more differentiated among the NE, CE and SW populations than was expected due to drift in our snmf F_st_ outlier test (FDR < 1%).

The number of the putative environmentally associated SNPs detected with LFMM2 differed among the variables tested (Figure S22). “Isothermality” and “mean diurnal range” had 5,679 and 2,794 putative SNP-environment associations identified, respectively. These were followed by “annual precipitation” (1,059 SNPs hits), “precipitation of driest quarter” (735 SNPs hits), “annual mean temperature” (391 SNPs hits), “flow direction” (263 SNPs hits), “mean temperature of wettest quarter” (227 SNPs hits), “TPI” (104 SNPs hits), “slope” (49 SNPs hits) and “precipitation seasonality” (3 SNPs hits). The variable “aspect” did not have any SNP association with an FDR < 1% in *LFMM2*. There was no significant correlation between the number of identified SNP association per variable and the percentage of contribution of each variable to the ENM (Pearson’s test p > 0.05).

The candidate adaptive SNPs were spread across the entire genome and present on all linkage groups in the *B. pendula* reference genome (Figure S22). The chromosome with the largest number of candidates is chromosome 2 (738 SNPs), while the lowest number was recorded for chromosome 6 (344 SNPs). The number of SNPs identified per chromosome does not appear correlated with chromosome size (Pearson’s test p > 0.05). A hotspot was visible in the second half of chromosome 7 (Figure S22).

The PCA based only on the 7,609 adaptive SNPs showed a steep discrepancy in the variance explained by PC1 compared to that of the other PCs (Figure S23). Furthermore, PCA based on only putatively adaptive SNPs did not show a clear separation between the central and north-eastern population in the first three PCs, while it separated the south-western population (Figure S23). When using only 7,500 putatively “neutral” SNPs in PCA, the separation between populations was slightly weaker compared to that observed with the whole data set, and individuals in each population were more scattered along the PCA axes (Figure S24). The pairwise F_st_ patterns of differentiation between populations based only the 7,609 adaptive SNPs and only on neutral loci reflected those observed with the whole SNPs set, however loci under selection showed substantially higher F_st_ compared to those recorded for neutral markers and for the whole SNP set (Figure S25-26).

We identified 1,633 mRNA regions in the B. pendula reference genome (Salojärvi et al., 2017) containing at least one candidate adaptive SNP,. A functional enrichment analysis of these genic regions reported significant hits (FDR < 5%) at all three gene ontology levels: biological process (3 hits), cellular component (1 hits) and molecular function (3 hits) (Figure S27).

### Risk of non-adaptedness to future conditions (RONA)

We calculated RONA for seven environmental variables projected under the future climate profile ssp370, identified as the worst-case scenario for *B. platyphylla* for the years 2080 – 2100 in terms of habitat reduction (Figure 5; Table S6). The most represented environmental variables in terms of number of associations in the RONA computation were “isothermality” (5,679 SNPs, mean r^2^ = 0.459), “mean diurnal range” (2,794 SNPs, mean r^2^ = 0.0157) and “annual precipitation” (1,059 SNPs, mean r^2^ = 0.0976) (Table S6). The variables that reported higher average r of adaptive SNPs were “isothermality” (5,679 SNPs, mean r^2^ = 0.459), “annual mean temperature” (391 SNPs, mean r^2^ = 0.195), and “mean temperature of wettest quarter” (227 SNPs, mean r = 0.185) (Table S6).

**Figure 5.**
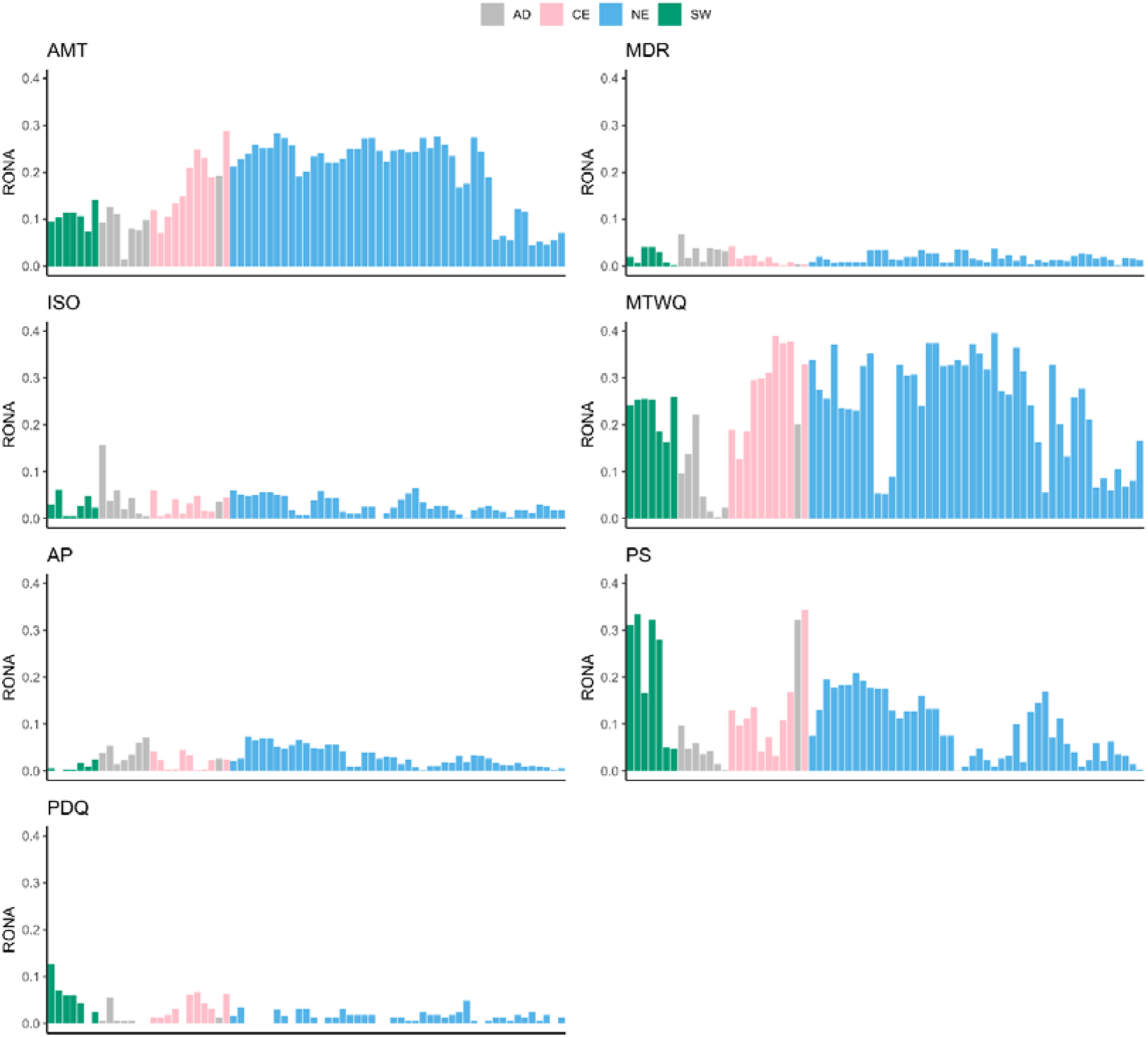
Risk of non-adaptedness (RONA) for the 71 *B. platyphylla* individuals. RONA calculated independently for each of the seven climatic variables projected to change in the future. The future climate profile ssp370 at 2080-2100 was used for RONA calculation. AD: admixed individuals. AMT: annual mean temperature. MDR: mean diurnal range. ISO: isothermality. MTWQ: mean temperature of wettest quarter. AP: annual precipitation. PS: precipitation seasonality. PDQ: precipitation of driest quarter.

There was large variability in the expected allele frequency changes required to match future conditions across environmental variables. “Annual mean temperature” and “mean temperature of wettest quarter” showed notably higher mean individual RONA compared to all the other variables, being 0.1773 and 0.2314 respectively (Table S6).

Weighted mean RONA scores were overall relatively low across individuals, with mean of 0.08 (sd = 0.03) and maximum of 0.1455, reported for an individual in the north-eastern population (HTY15) (Figure 6A; Table S7). The maximum RONA scores showed greater variability across individuals, with mean of 0.25 (sd = 0.09) and maximum of 0.3956, recorded for an individual in the north-eastern population (JY23, “mean temperature of wettest quarter”) (Figure 6B; Table S7).

**Figure 6.**
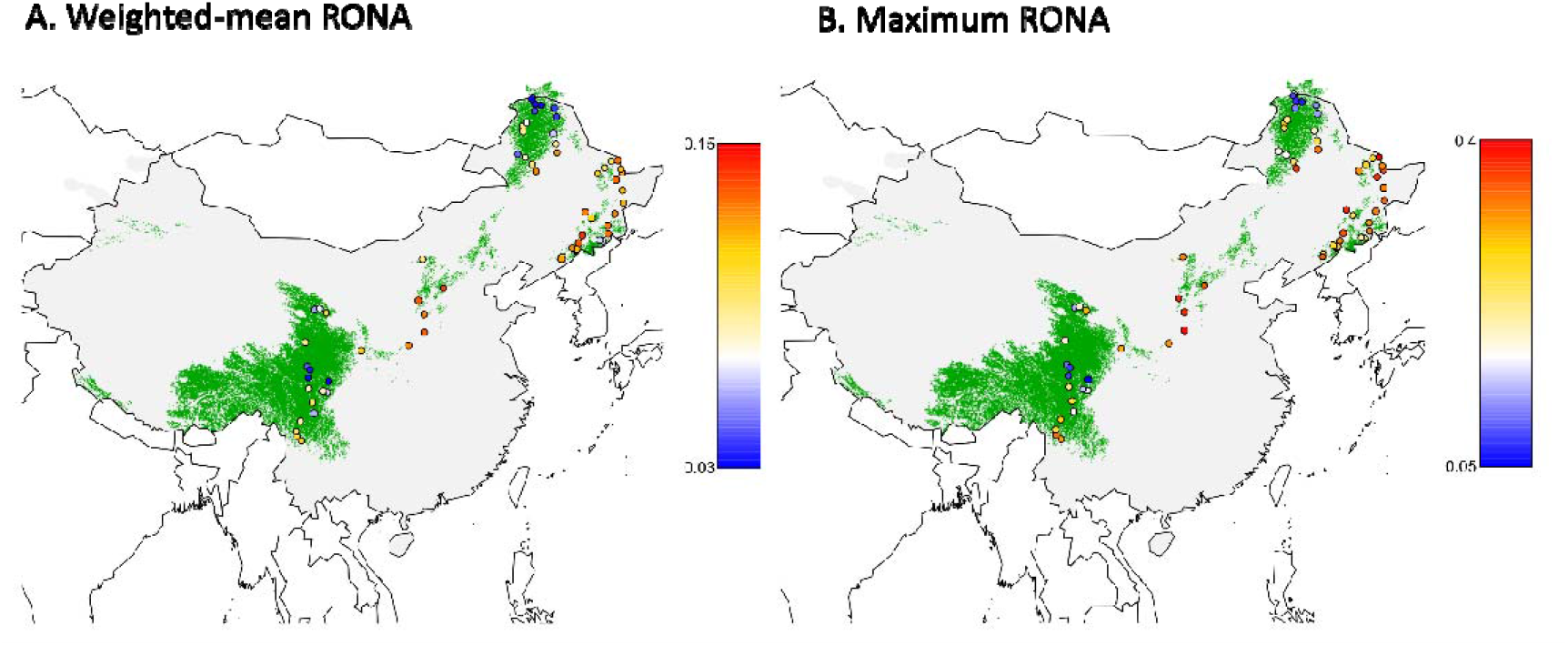
Weighted-mean RONA and MAX RONA. **A)** Map showing weighted mean RONA for the 71 *B. platyphylla* individuals, out of the seven environmental variables included in the analysis. Each variable RONA was given a weight equivalent to its percent contribution in the ENM. **B)** Map showing maximum RONA per individual, out of the seven environmental variables tested. Green shading represents suitable environment in 2080-2100 under profile ssp370 according to our ENM.

There were some differences between the three Chinese *B. platyphylla* populations in regard to the environmental variables that will require greater allele frequency changes (in ssp370 at 2080 – 2100) (Table S8). Although all populations showed similar RONA patterns across the environmental variables tested, with the largest mean RONA (averaged over the individuals of each population) reported for temperature derived variables (“annual mean temperature” and “mean temperature of wettest quarter”) rather than precipitation-derived variables (Table S8), the south-western population, had much higher mean RONA for “precipitation seasonality” (0.21) compared to the other two populations that had relatively low mean RONA for this specific environmental variable (Table S8).

## Discussion

### Pan Eurasian population genetic structure of white birch

Our study clearly differentiated between *B. pendula* and *B. platyphylla* across Eurasia, showing strong separation between the two in PCA (Figure S8) and *fastSTRUCTURE* (Figure 2) analyses. We found evidence for extensive hybridization between these two species in NW China, and to a lesser extent in central Russia. Populations in NW China that were initially identified as *B. platyphylla* turned out to be mainly *B. pendula* with introgression from *B. platyphylla*. A previous microsatellite study of *B. pendula* and *B. platyphylla* across Eurasia (Tsuda et al., 2017) showed admixture between the two species in Siberia, but in Asia this study only included populations from Russia and Japan. In a microsatellite study of white birches in China, Chen & Lou (2019) found populations in northwest China to be highly differentiated from other populations, and to be of high genetic diversity. However, as no *B. pendula* were included in their study, they could not pick up a signature of hybridization in these populations. Instead, they attributed their high differentiation and diversity to a northern glacial refugium for white birch in the Altay Mountains. If we had not co-analysed our data with that of Salojärvi et al. (2017), we might also have concluded that the populations in NW China were a divergent lineage, rather than introgressed. In this study, we removed individuals from the NW population from our downstream analyses on *B. platyphylla*.

### Genetic structure of white birch within China and signals of local adaptation

With the NW individuals excluded, *B. platyphylla* in China has a linear distribution from north-eastern to south-western China, along the edge of the inland mountainous region. Occurrence data and environmental niche models suggest that there is currently a single contiguous region currently habitable by *B. platyphylla*. Within our *B. platyphylla* samples, we found three genetic clusters with some admixed individuals between them. The three clusters seem to mirror similar phylogeographic breaks in many other plant species (Zhout et al., 2021; Ye et al., 2017; Fan et al., 2016; Bai et al., 2016) and our interpretation of B. platyphylla population structure is similar to that of Chen & Lou (2019). Our fastsimcoal results suggest that the three *B. platyphylla* populations exchanged genetic material soon after their divergence and recently, but endured a long period of isolation in between these contacts (Figure 3D). The absolute date estimated from fastsimcoal suggests that this period of isolation between the *B. platyphylla* populations was from 2.8 Mya to 0.8 Mya. This is hard to fit with estimates of the time of speciation between *B. platyphylla* and *B. pendula* which different studies place between 36,000 years (Tsuda et al., 2017) and 2.6 million years (Chen et al., 2021) ago. We note, however, that our demographic simulation results are provisional as it is hard to distinguish among the many factors that affect patterns genetic variation (Johri et al., 2022), and our assumptions about mutation rate and generation time may be incorrect. We cannot therefore exclude the possibility that our simulations are picking up signals of *B. platyphylla* population isolation due to separate refugia in recent glaciations.

Consistent with a previous phylogeographic study of Asian white birch, showing that microsatellite diversity of Asian white birch increased with latitude (Chen & Lou, 2019), our results confirmed that genomic diversity in the NE is higher than that in CE which is higher than that in SW (Figure S14-15). This favours the scenario of separate glacial refugia at different latitudes, rather than a Europe-like postglacial expansion from southern refugia (Birks & Willis, 2010; Hewitt, 1999). This pattern may also reflect less severe geographical barriers to pollen movement in northern China compared to the south-western Qinghai-Tibet Plateau. It might have been expected that higher diversity in the NE population is due to admixture from *B. pendula*, but we find no evidence for this.

Ecological niche models indicated that mean temperature of wettest quarter and annual mean precipitation are the most important predictors of *B. platyphylla* distribution and both north-eastern China and the south-western Tibetan plateau currently provide the most suitable environments for this species in China. Future climate projections show an overall decline of the species range and environmental suitability throughout of its current distribution in China, with suitable habitats persisting mostly in areas of higher elevation (Figure 4) and the central China habitable area vulnerable to being lost.

Our PCA based only on adaptive SNPs showed that the NE and CE populations formed an overlapping cluster whereas in PCA based on neutral SNPs the two populations were separated (Figure S23-24). This is possibly due to the relatively lower diversifying selection acting on the two populations and the existence of gene flow which can counteract the impact of diversifying selection (Guichoux et al., 2013; Nosil et al., 2009). F_st_ analyses suggested that the level of differentiation is much higher at loci under selection rather than neutral loci, as expected, and the SW populations appears to be highly differentiated from the others at adaptive loci (Figure S25).

Functional enrichment analysis of mRNA intersecting the identified adaptive SNPs suggested that these genes were mostly enriched for growth and environmental stress response (Figure S27), although further studies and functional experiments are needed to confirm these findings. Among the significantly enriched categories, the most interesting in view of climate adaptation were: “regulation of response to stimulus”, “DNA helicase activity” which are enzymes involved in DNA repair and have a crucial role as caretakers of the plant genome against environmental damages (Sami et al., 2021) and “inorganic cation transmembrane transporter activity”, as potassium, the most abundant inorganic cation in plant cells, it is essential for plant growth and development and it has been shown to have a major role in resistance to drought, salinity and fungal infections (Sharma et al., 2013). The 1,633 identified adaptive genes also included several light-response and growth-related genes, according to our OmicsBox annotation, as well as members of all the three GO categories significantly enriched in the putative genes under selection detected by Solojarvi et al. (2017): transmembrane receptor protein tyrosine kinase, peptidyl-histidine phosphorylation, and axis specification. Furthermore, 87 of the 1,633 genic regions detected in this study were identified as putative selective sweeps in *B. pendula* (Salojarvi et al., 2017). Even though it would be interesting to explore all the identified genes singularly, this goes beyond the extent of this work and further experimental validation would still be necessary to confirm their role in growth and environmental stress response.

In line with similar studies in other species (Jordan et al., 2017; Rellstab et al., 2016; Borrell et al., 2019; Pina-Martins et al., 2018), local adaptation appears to be a highly polygenic in *B. platyphylla*, therefore it is likely that maintaining the standing variation and adaptive diversity may be a better solution to aid future climate adaptation, rather than focusing on raising the frequencies of a specific set of adaptive SNPs, as it was proposed previously (Jordan et al. 2017). Two variables, “mean diurnal range” and “isothermality” reported a substantially larger number of significant SNPs association compared to the other climatic variables (Figure S22), which may have arisen due to the spatial correlation of these environmental variables and genotypes across the sampling range therefore these results should be interpreted with caution and further validation is necessary, however we cannot exclude and highly polygenic adaptation to these variables with many SNPs with low effects involved. We did not identify any significant correlation between the number of SNPs associated with each climate variable, and their percentage contribution in our ENM, unlike some other studies (Borrell et al., 2019; Rellstab et al., 2016). However, we note that it is not a logical necessity as the variables with higher discriminatory power in the ENM could be limiting species ranges either because they lack adaptation (Borrell et al., 2019), or contrarily they could be subjected to strong adaptation but at a global level throughout the species range, rather than locally. Therefore, the lack of variation in these variables across the sampling range prevents the detection of these adaptive signals with methods designed for local selection, such as LFMM2.

To investigate how the samples of *B. platyphylla* will respond to rapid future climate change, we calculated RONA for the 71 *B. platyphylla* individuals according to the implementation by Pina-Martins et al. (2018). Overall, RONA was relatively low across the environmental variables tested (Figure 5), with the exception of “annual mean temperature” and “mean temperature of wettest quarter”, very likely reflecting the larger relative projected change of temperature derived variables compared with that of environmental variables related to precipitation throughout China. The expected allelic frequencies changes are similar to those reported in other studies (Jordan et al., 2017; Pina-Martins et al., 2019; Rellstab et al., 2016). Weighted mean RONA incorporated the ENM with EAA and generally remains low and it is always below 0.2 (Figure 6A). We also reported the maximum RONA per individual out of the seven variables tested (Figure 6B) because even if a population, in this case an individual, has “low” average RONA for a given future projection, it might still be its highest RONA that will determine how much it will need to shift its allelic frequencies in order respond to future selective pressure (Pina-Martins et al., 2018). Maximum RONA across individuals showed significantly higher values, as expected. Interestingly, the maximum RONA plot also showed particularly high values for four individuals in the south-western population located at the very southern edge of this population distribution, reported for precipitation seasonality which is expected to have a much larger relative change at these locations (Figure 6B). Overall, both weighted mean and maximum RONA match well with our ENM for ssp370 2080-2100, with lower RONA values reported within the inferred projected suitable environment (Figure 6).

The RONA method is of course a huge simplification, and its limitations are accurately described in the original publication (Rellstab et al., 2016), therefore our results should be viewed with caution. It should also be noted that RONA was originally designed to be used with population allelic frequencies rather than individuals, but given the nature of our sampling, using individuals was to our knowledge the best approach. Grouping the individuals in populations for the RONA calculation would have been problematic as it would have required averaging climatic conditions across different locations. This would have led to inaccuracy due to the broad geographic range of our study and the high heterogeneity of some of the environmental variables tested.

It is not straightforward to determine when RONA is high enough to result in a lag between allele frequencies and adaptation. A previous study on Fagus sylvatica have observed allele frequencies changes of 0.1 – 0.2 per decade (Jump et al., 2006), but this estimate was based on amplified fragment length polymorphism (AFLP) molecular markers rather than SNPs. A more recent study on Swiss stone pine, Pinus cembra, based on SNPs markers, investigated allele frequencies shifts over time by looking at the differences in SNPs frequencies between adults and juvenile cohorts at seven locations in Switzerland, and validated the observed shifts with forward-in-time simulations (Dauphin et al., 2020). This analysis reported an average rate of allele frequency shifts of 1.26 × 10^−2^ per generation (P. cembra generation time = 40 years) for neutral SNPs, and a slightly lower estimate for adaptive SNPs. Based on this estimate we can hypothesize that SNPs frequencies changes < 0.05 may be achieved by populations naturally within a couple of generations, whereas larger changes, such as those predicted for temperature derived variables (Figure 5), are unlikely to be achieved naturally by long-living forest trees with long-generation times, even when they exhibit high levels of standing genetic variation. This reinforces the importance of considering conservation strategies such as assisted gene flow or assisted migration (Borrell et al., 2019).

### Concluding remarks

This study provides insights into both the evolution and ecology of Asian white Birch, *B. platyphtylla*, in China. We showed that *B. platyphylla* and *B. pendula* are better considered distinct species, as their differentiation is clear at the genomic level. We identified a hybrid zone between *B. platyphylla*/*B. pendula* in north-western China and three distinct lineages of *B. platyphylla* in eastern and southern China distributed along a latitudinal gradient from the north-east to the south-west, and. The three *B. platyphylla* lineages appear to have had a long period of isolation from one another, followed by more recent gene flow. Our species distribution model shows that *B. platyphylla* prefers mountainous environments with cool summer and moderate levels of precipitations. Future prediction shows significant reduction on the suitable habitat of this species in China under every future scenario tested, up to a third reduction compared to the current extent. We identified signals of local adaptation throughout the genome of this species, suggesting that climate adaptation is a highly polygenic mechanism. We estimated the degree of maladaptation to inform conservation strategies. Despite the limitation of our method, we showed that rapidly rising temperature, particularly in the summer months, poses a risk to this species and our environmental niche model suggests that current habitats located in central China and in the north-eastern region adjacent to the Changbai mountains will likely become unsuitable by 2080-2100, while current habitats located at higher elevations, such as the south-western Tibetan plateau and the region surrounding the Greater Khingan mountains range in the north-east will remain suitable. It is possible that populations outside the predicted suitable environment may be able to adapt, however climate, particularly temperature, might change to an extent that adaptation will not be possible and assisted migration might be the only option possible to rescue some populations. The estimated required changes in allelic frequencies at adaptive loci do not appear as concerning within the predicted suitable environment, however with the current dataset, based on individuals’ allelic frequencies, it is difficult to estimate with confidence an allelic frequency change threshold after which populations will fail to adapt on time naturally and will require assisted gene flow from donor pre-adapted populations.

## Acknowledgements

This work was funded by the National Natural Science Foundation of China (31770230 and 31600295) to Nian Wang. Gabriele Nocchi is on a PhD programme run jointly between Queen Mary University of London and the Royal Botanic Gardens, Kew funded by the UK Government Department for Environment, Food and Rural Affairs (Defra), in association with Action Oak.

We thank Stefan Götz, David Seide and the BioBam Bioinformatics team for kindly letting us use their software OmicsBox for the functional annotation and functional enrichment analysis of the identified adaptive genes. We thank Richard Nichols (Queen Mary University of London), Yiye Liang and Baosheng Wang (South China Botanical Garden, Chinese Academy of Sciences) for their expert advice on our analyses.

This research utilised Queen Mary’s Apocrita HPC facility, supported by QMUL Research-IT: http://doi.org/10.5281/zenodo.438045.

## Data accessibility

Sequencing read data are archived in the NCBI through the BioProject accession PRJNA854976.

### Benefit sharing statement

This research was led by scientists from the country providing genetic samples. As described above, all data have been shared with the broader public via appropriate biological databases.

## Author contributions

RB and NW conceived the project. NW and JD performed the sampling and extracted DNA. LY, JD, and YG performed mapping and SNPs calling. GN and JW analysed the data and wrote the manuscript. RB oversaw the project and helped write the manuscript.

## Notes

### Competing Interest Statement

The authors have declared no competing interest.

### Summary of Updates

ABBA-BABA test was added to clarify hybridization; Fastsimcoal results updated to include more scenarios; Figure 2 revised; Supplemental files updated.

