## Supplemental tables and figures for "Genomic signals of local adaptation and hybridization in Asian white birch"

**Figure S1.** The twenty-eight models of three populations evaluated using fastsimcoal2. Model 01-Model 14 show CE lineages was divergent from the common ancestral population of CE and SW. Model 15-Model 28 show CE lineages originated as a hybrid between the SW and CE lineages. Models 01, 04, 07, 10, 15, 18, 21 and 24 show divergence with no gene flow; Models 02, 05, 08, 11, 16, 19, 22 and 25 show divergence with continuous gene flow among the NE, CE and SW lineages; Models 03, 06 ,09, 13, 17, 20, 23 and 27 represent divergence with continuous gene flow between NE and CE and between SW and CE; Model 12 and 26 represent divergence with ancient gene flow and gene flow after secondary contact among the three lineages (no gene flow during the period of bottleneck); Model 14 and 28 show divergence with ancient gene flow and gene flow after secondary contact between NE and CE and between SW and CE (no gene flow during the period of bottleneck).

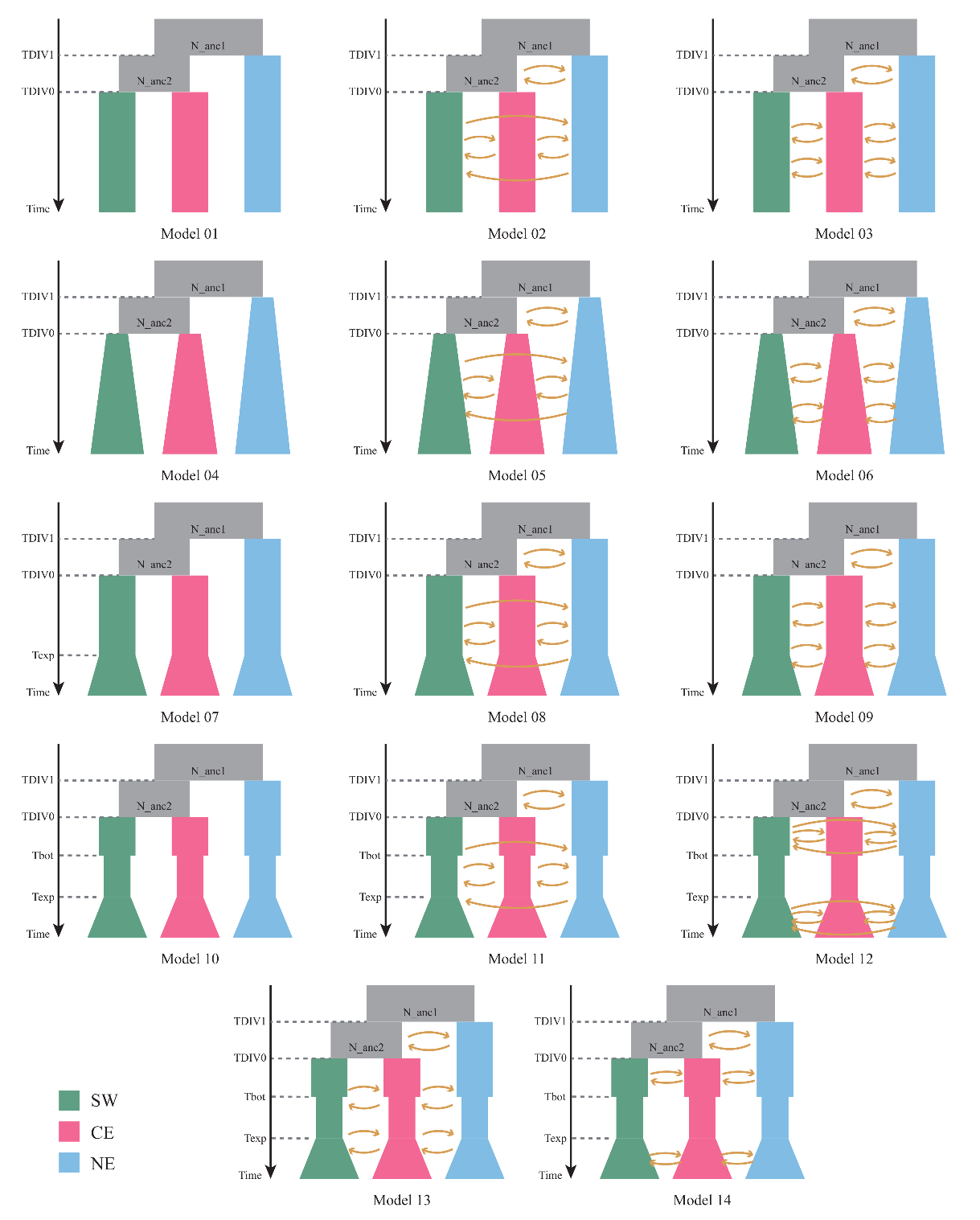

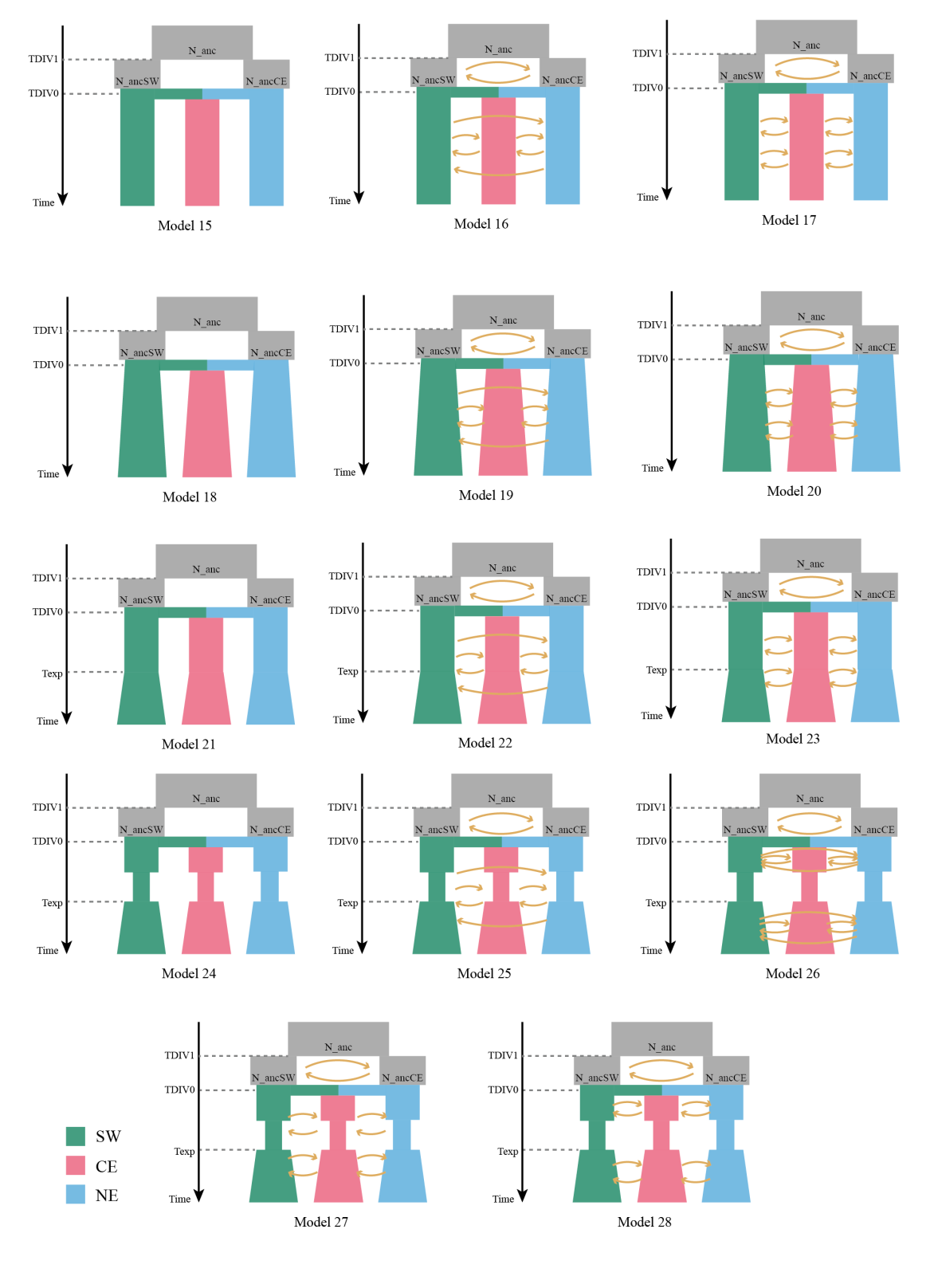

**Figure S2.** The nine models of two populations evaluated using fastsimcoal2. Models 2pop 01, 03, 05 and 07 show divergence with no gene flow; Models 2pop 02, 04, 06 and 08 show divergence with continuous gene flow among the NE and SW lineages; Model 2pop 09 shows divergence with ancient gene flow and gene flow after secondary contact between NE and SW lineages (no gene flow during the period of bottleneck).

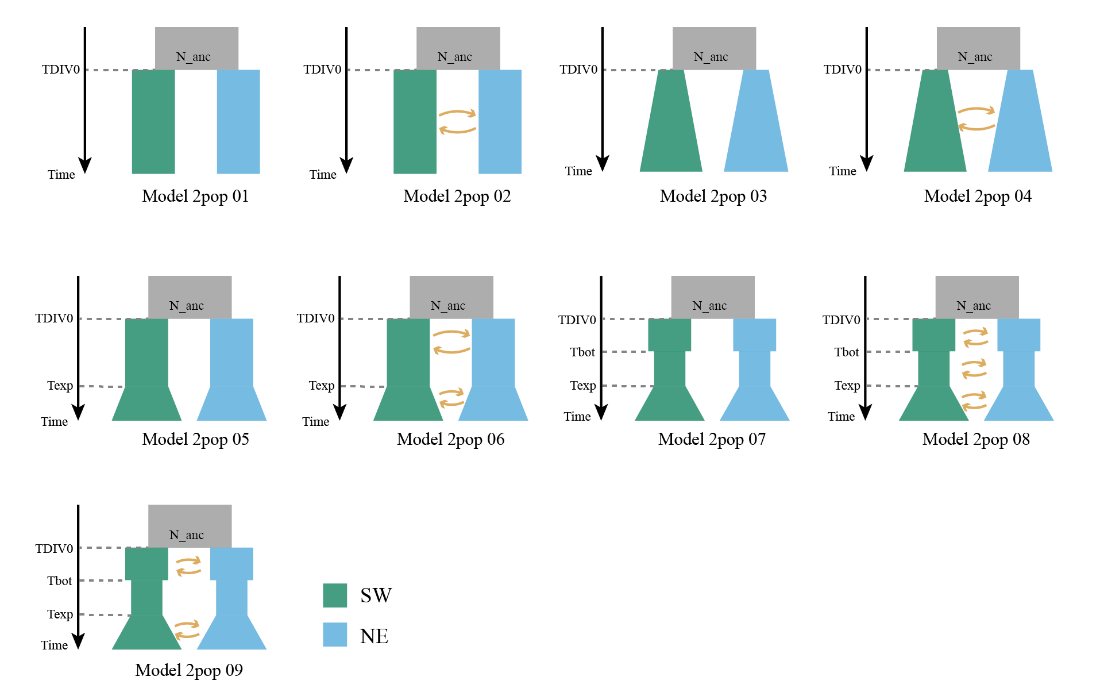

**Figure S3.** Goodness-of-fit of the demographic models inferred by fastsimcoal2. A) Results for the three populations demographic model and B) results for two populations model. Colours in observed (data) joint site frequency spectrum (SFS) and predicted (model) joint SFS reflect the number of SNPs. Residuals between data and model are plotted in a colourmap and a histogram. In the colour map, red or blue residuals indicate that the model predicts excessive or insufficient alleles in a given cell, respectively.

A)

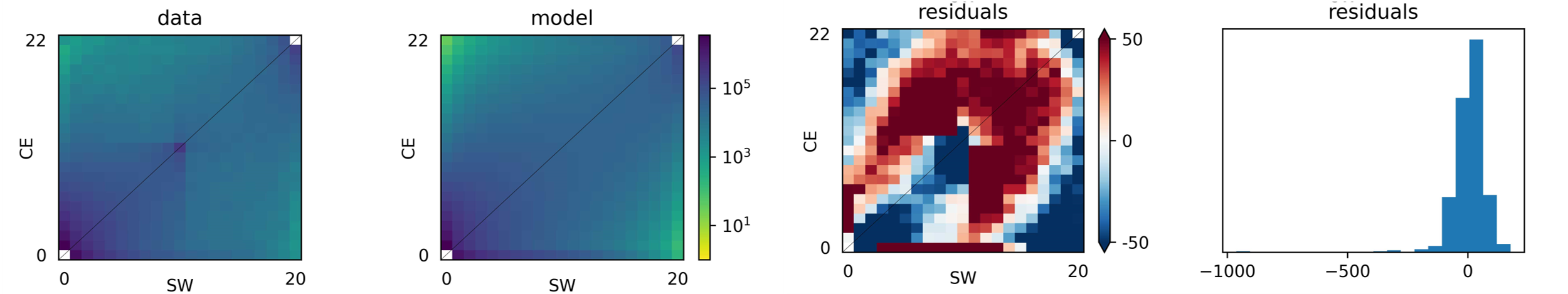

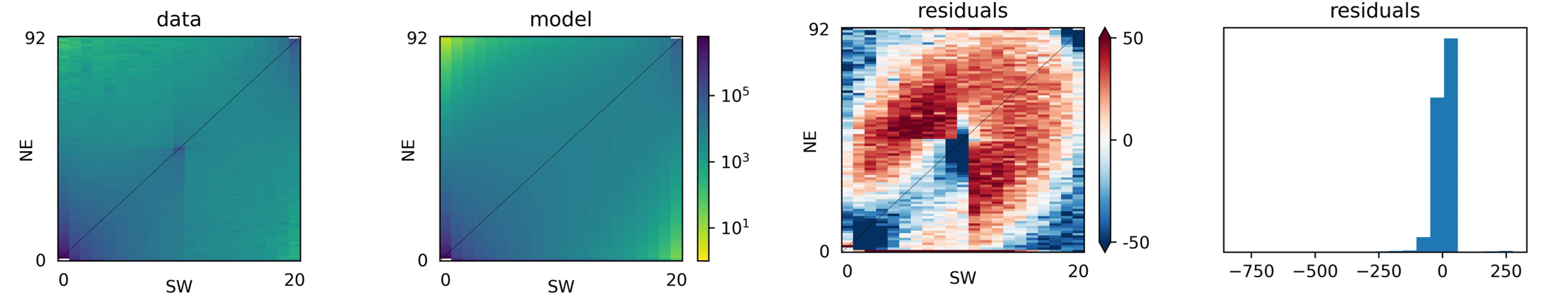

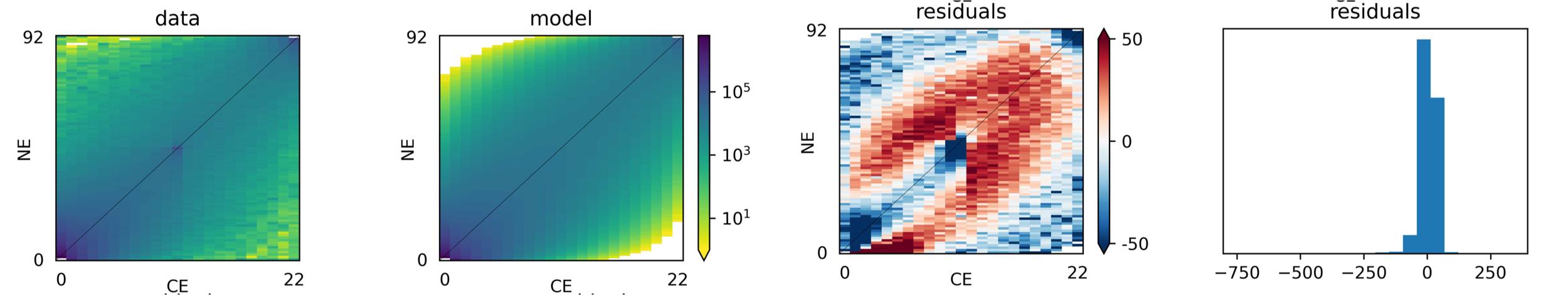

B)

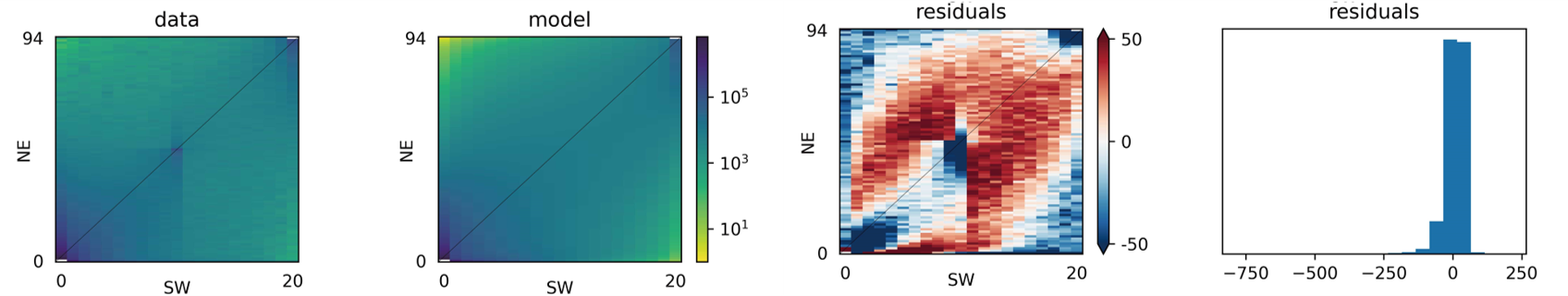

**Figure S4.** The 11 uncorrelated environmental variables selected for ENM and EAA.

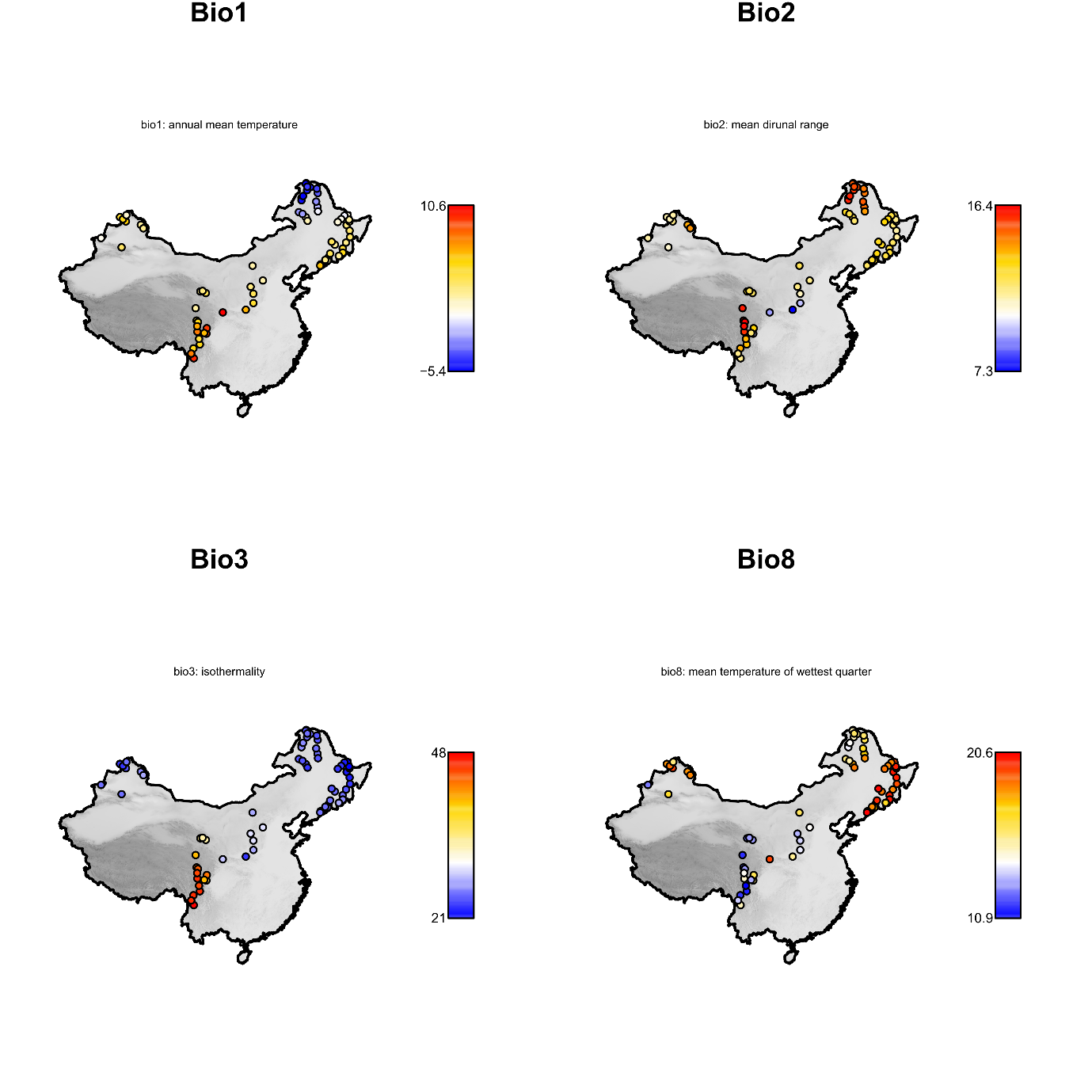

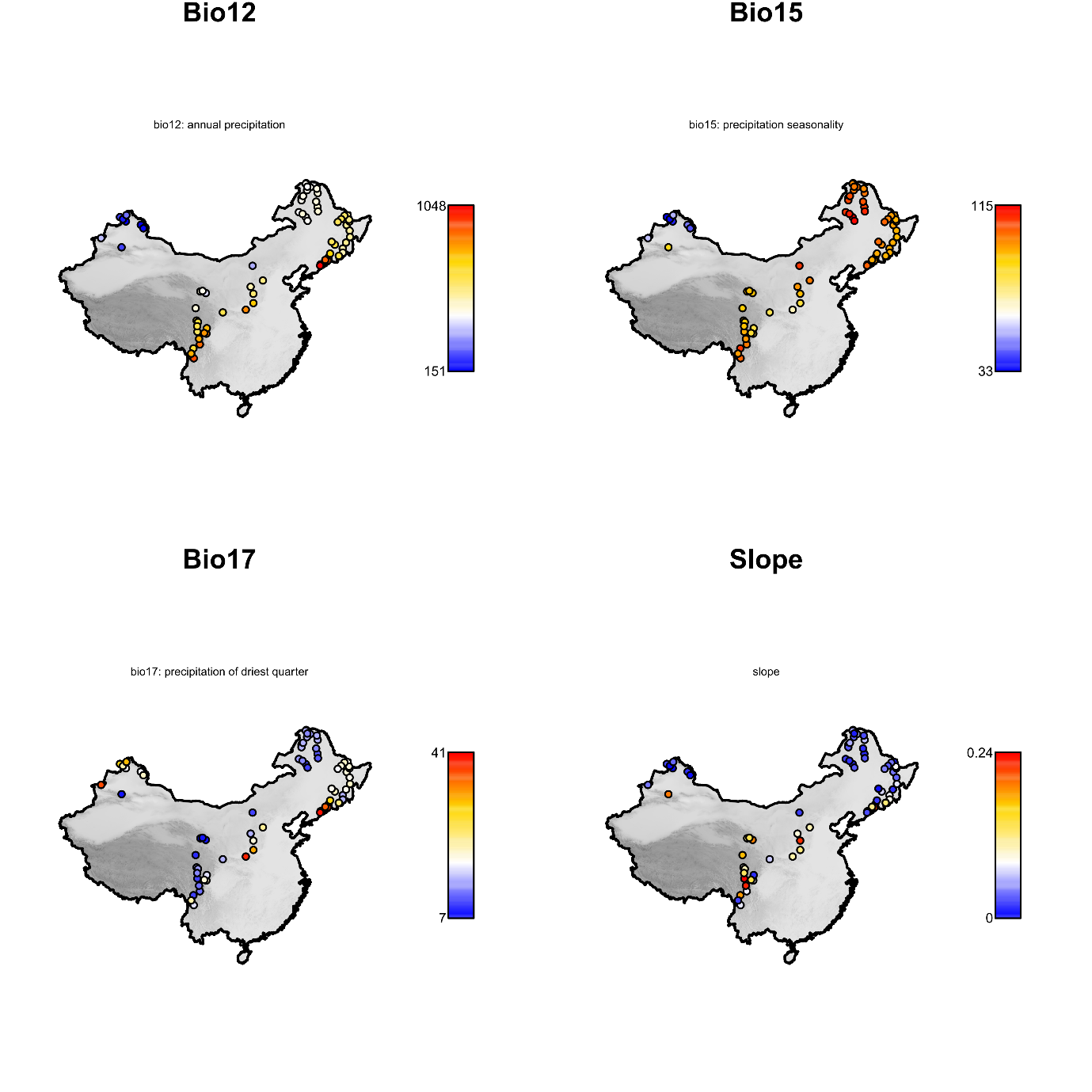

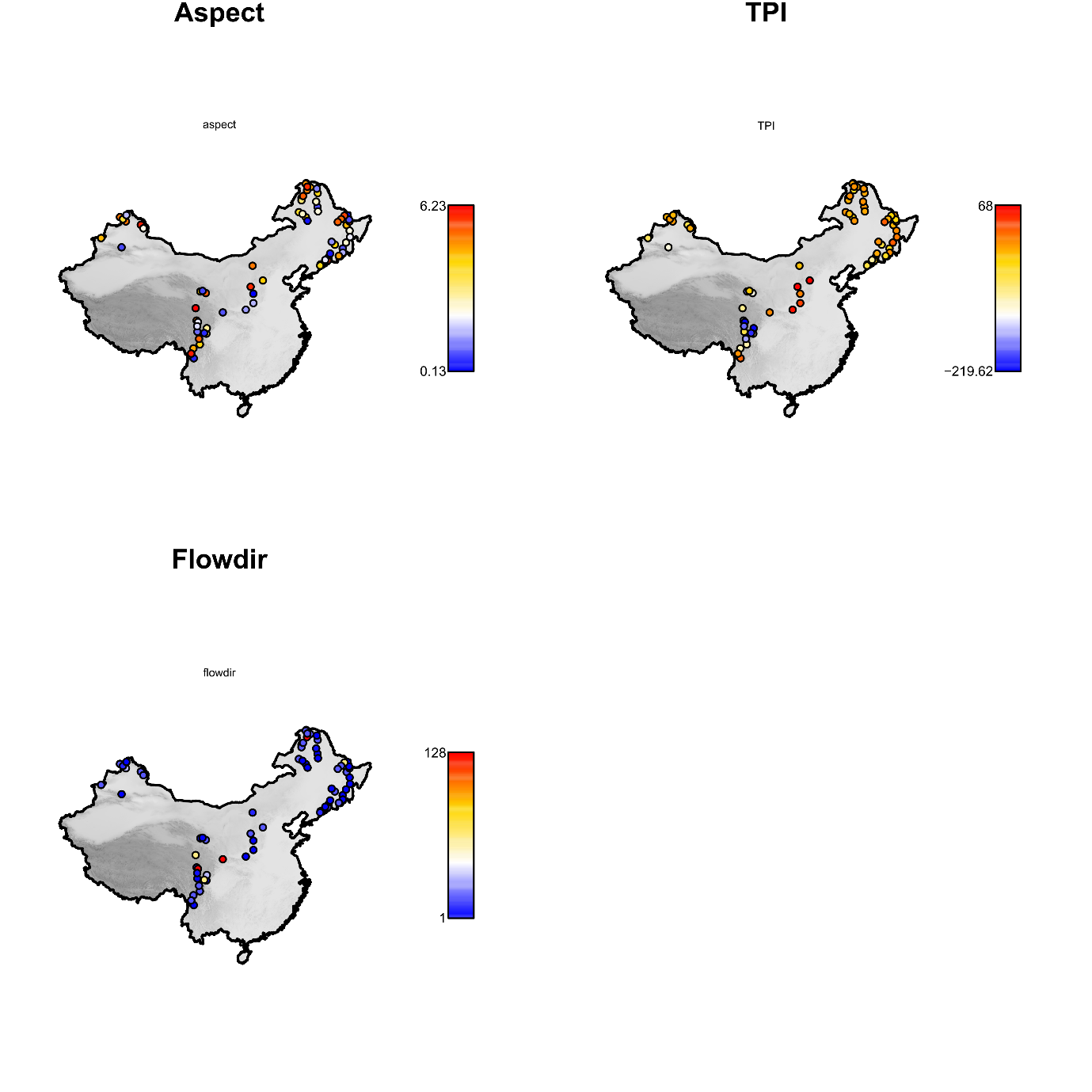

**Figure S5.** Correlation between the original 26 climatic variables.
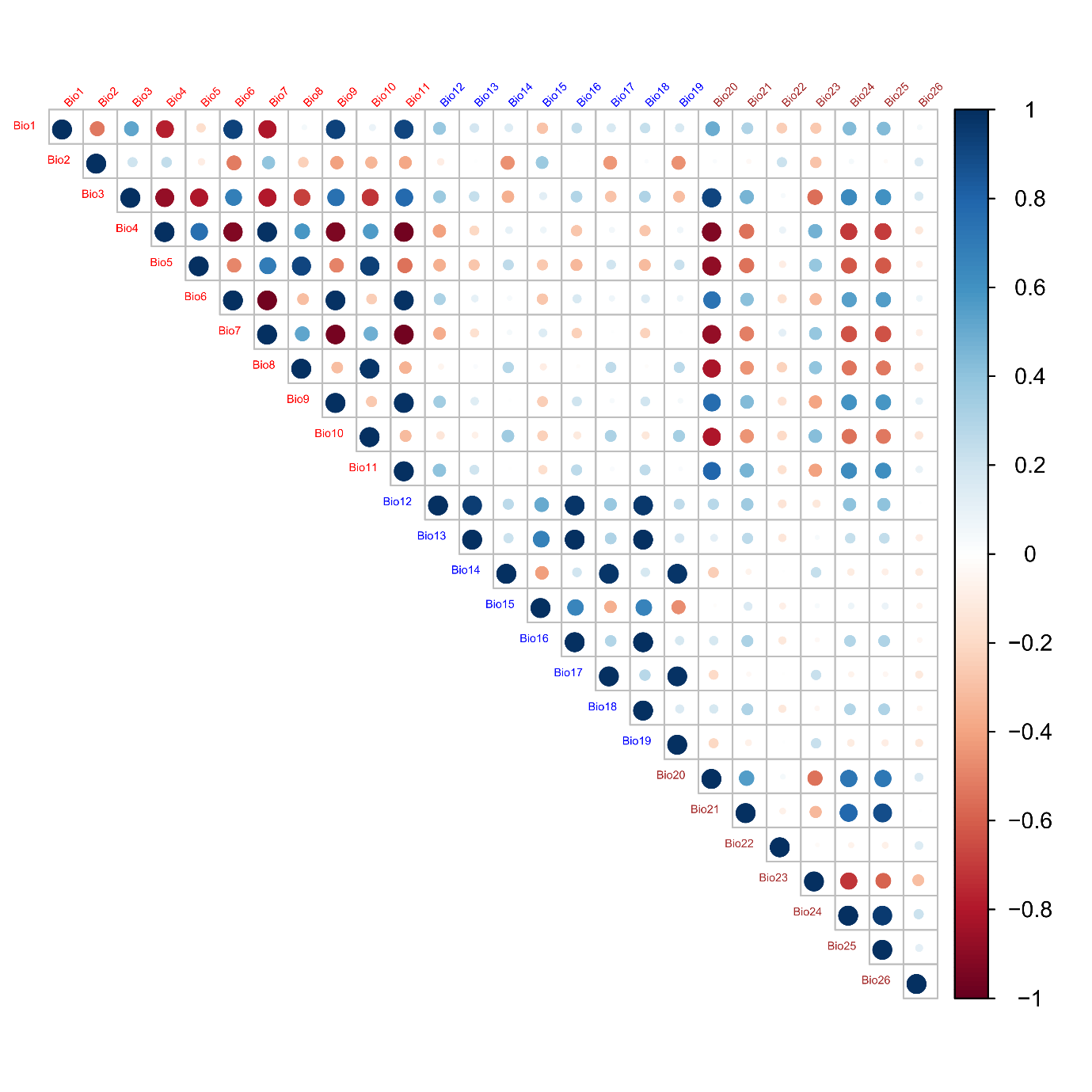

**Figure S6.** Results of the fastSTRUCTURE analysis including 83 birch individuals. A) Log-marginal likelihood lower bound (LLBO) of the data for Ks from 1 to 10. B) Cross-validation error calculated using 5-fold cross-validation for Ks from 1 to 10, with standard error bars.

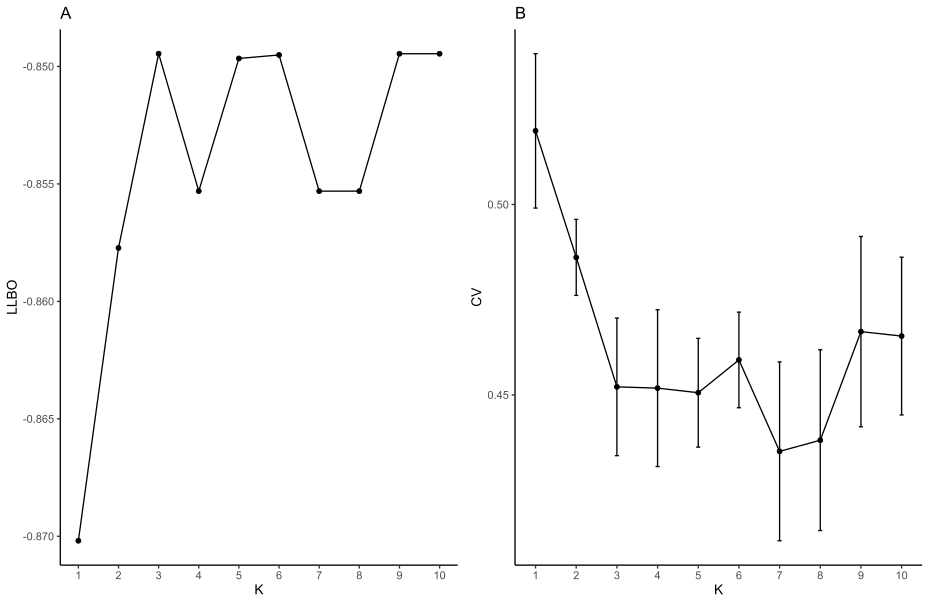

**Figure S7.** Results of the fastSTRUCTURE analysis including 162 *B. pendula* and *B. platyphylla* individuals. A) Log-marginal likelihood lower bound (LLBO) of the data for Ks from 1 to 10. B) Cross-validation error calculated using 5-fold cross-validation for Ks from 1 to 10, with standard error bars.

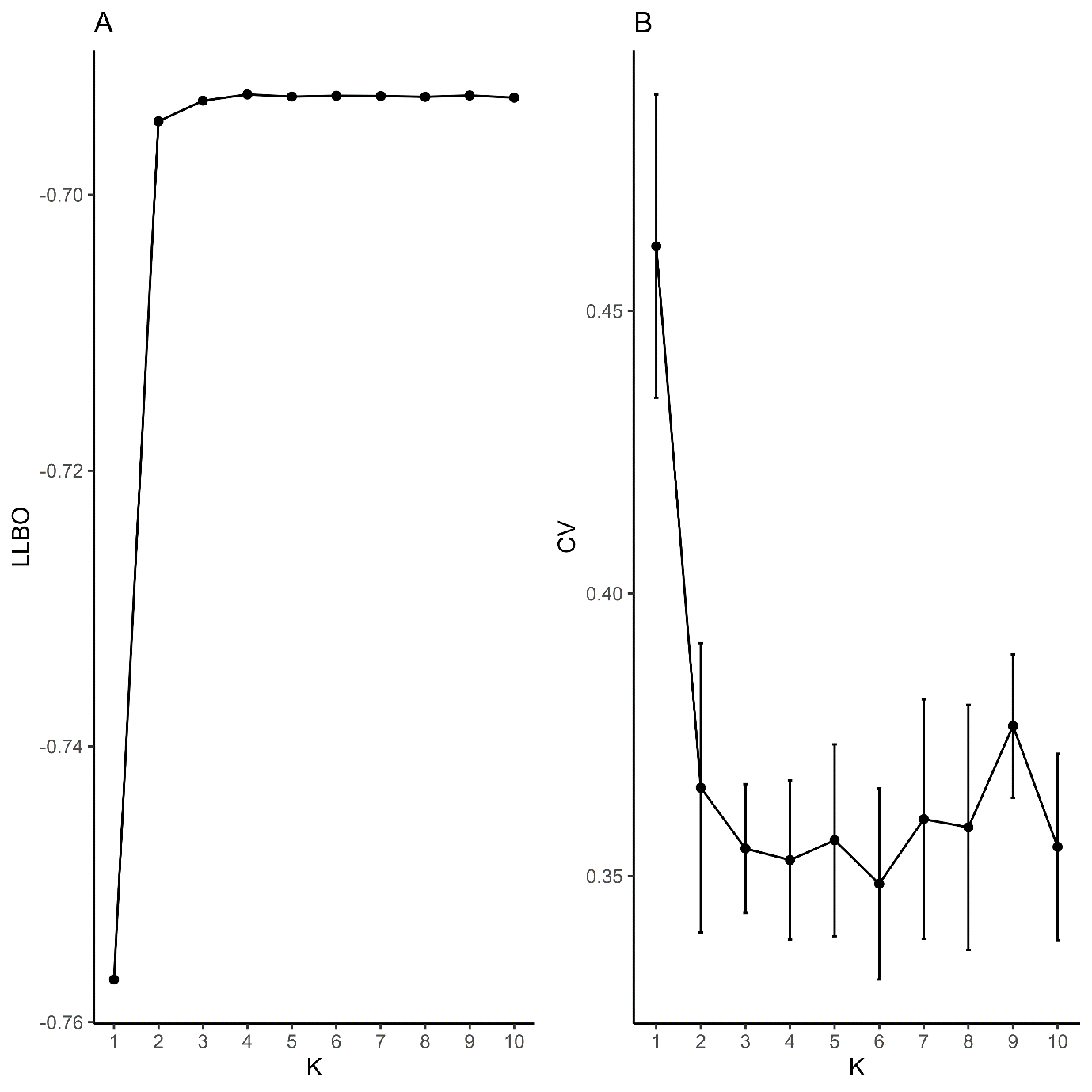

**Figure S8.** PCA of 162 *B. pendula* and *B. platyphylla* individuals based on 278,717 unlinked SNPs (r2 < 0.4). Colours represent populations assignments. A) PC1 against PC2. B) PC1 against PC3. C) PC2 against PC3. D) Eigenvalues of the computed principal components.

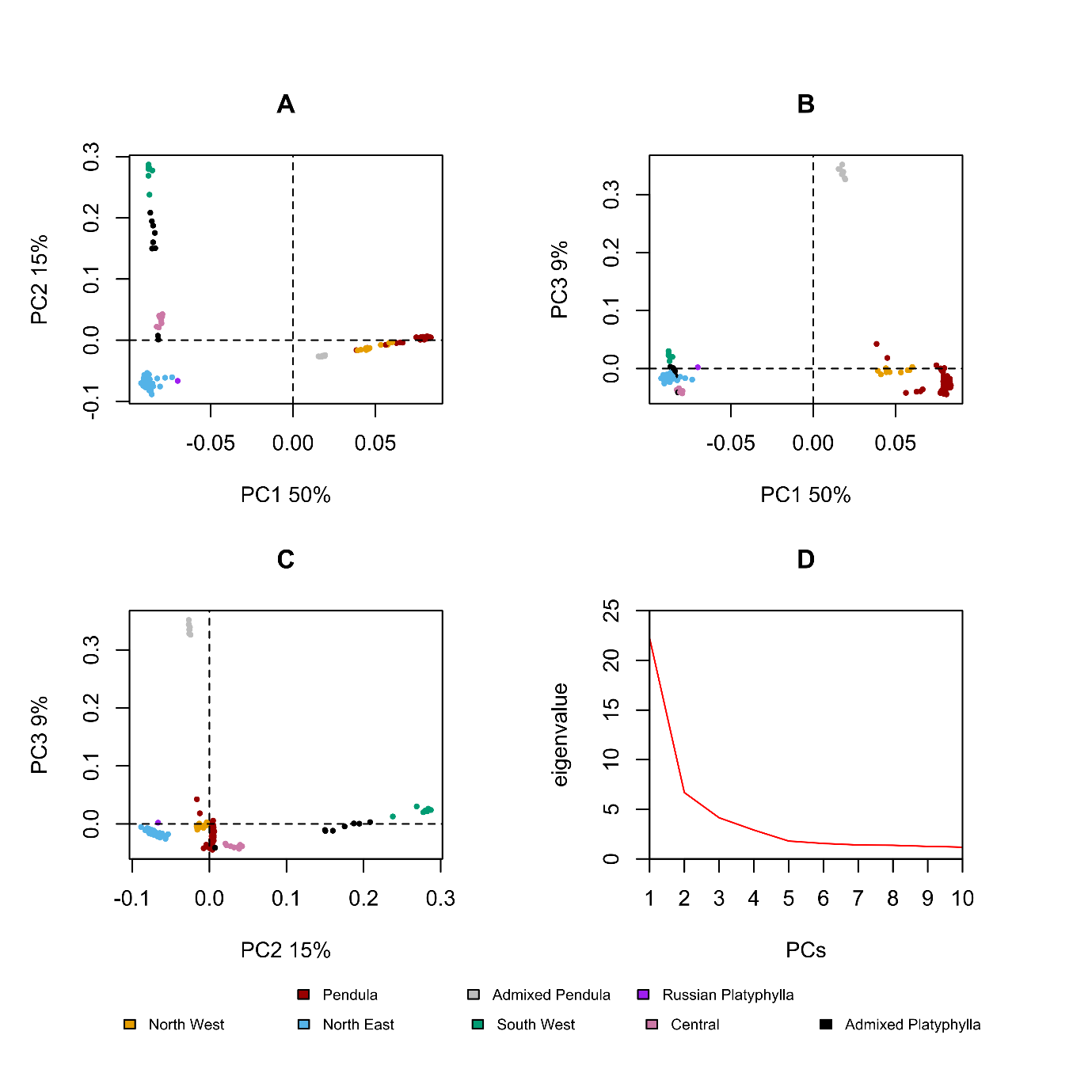

**Figure S9.** Results of the fastSTRUCTURE analysis including 71 *B. platyphylla* individuals. A) Log-marginal likelihood lower bound (LLBO) of the data for Ks from 1 to 10. B) Cross-validation error calculated using 5-fold cross-validation for Ks from 1 to 10, with standard error bars.

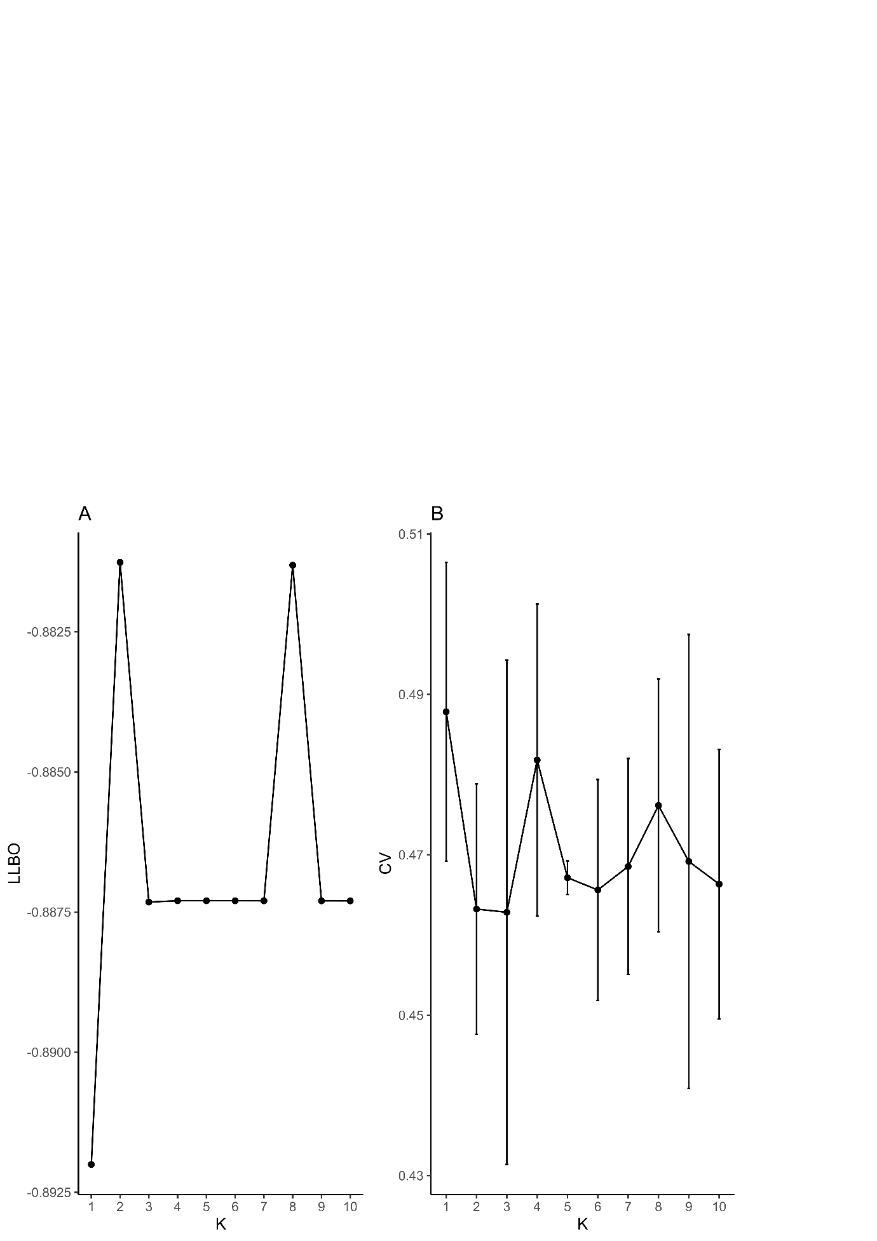

**Figure S10.** PCA of 71 *B. platyphylla* individuals based on 1,387,994 unlinked SNPs (r^2^ < 0.4). Colours represent fastSTRUCTURE populations assignments at K = 3. A) PC1 against PC2. B) PC1 against PC3. C) PC2 against PC3. D) Eigenvalues of the computed principal components.

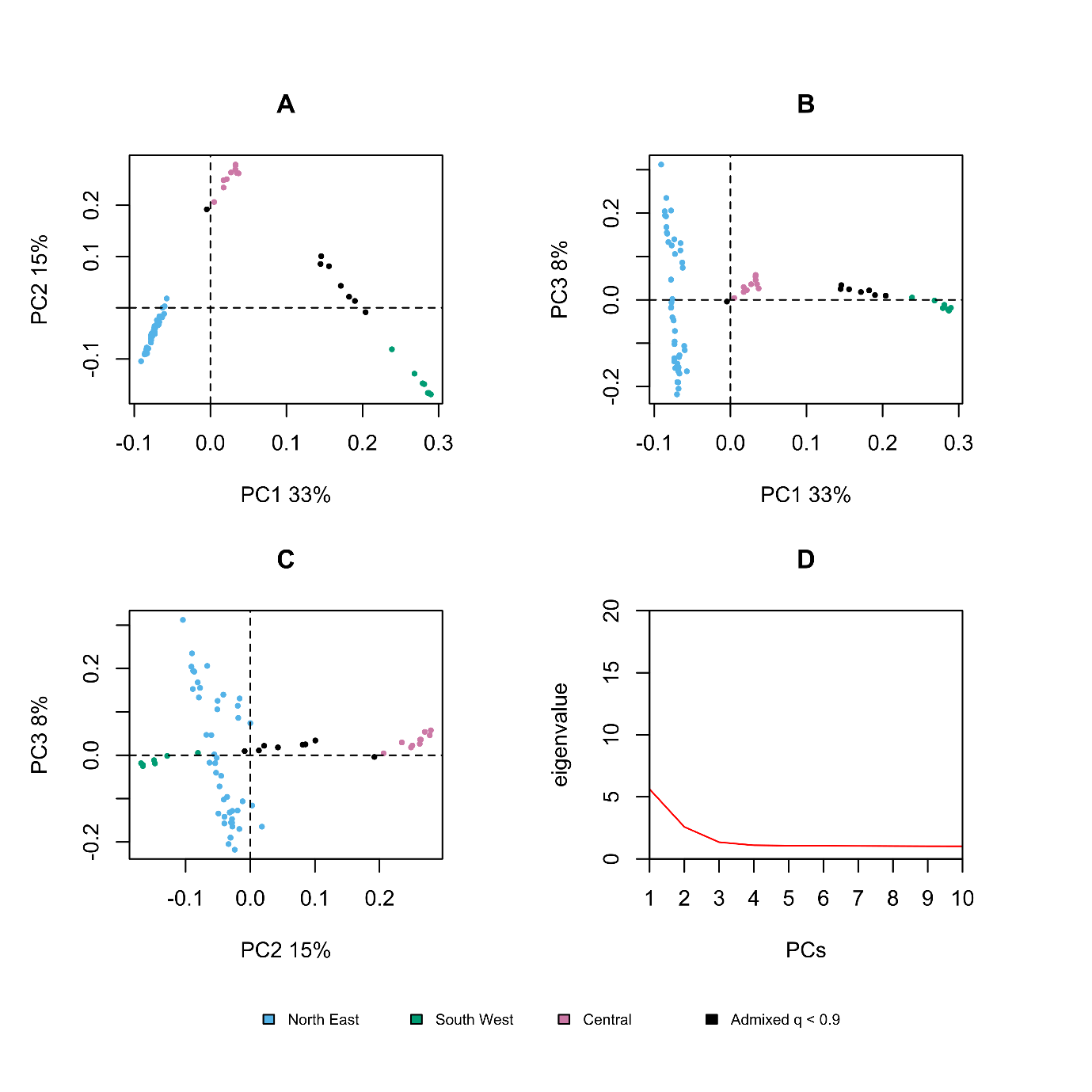

**Figure S11.** *snmf* Fst outlier test. A) Histogram of outlier test p-values. B) -log10 of p-value for each SNP.

**A)**

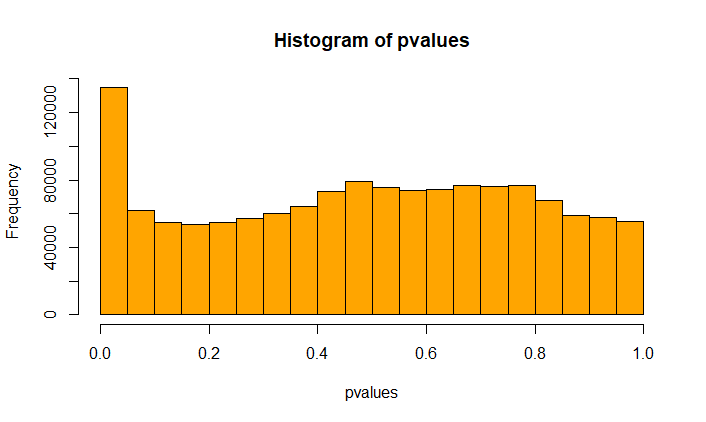

**B)**

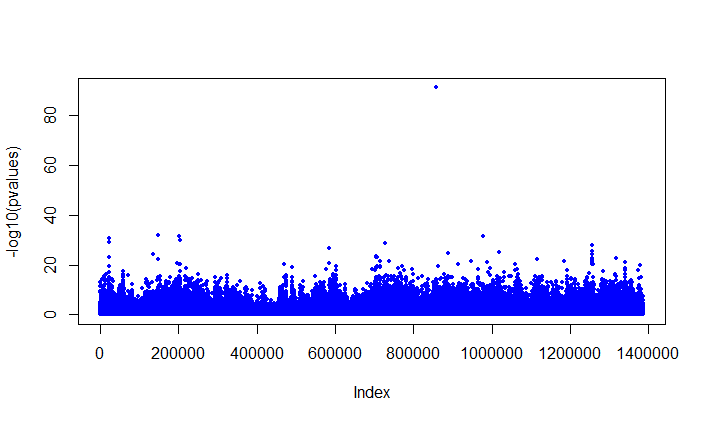

**Figure S12.** Linkage disequilibrium (LD) decay in *B. platyphylla* populations, calculated with the tool PopLDDecay (Zhang, Dong, Xu, He, & Yang, 2018).
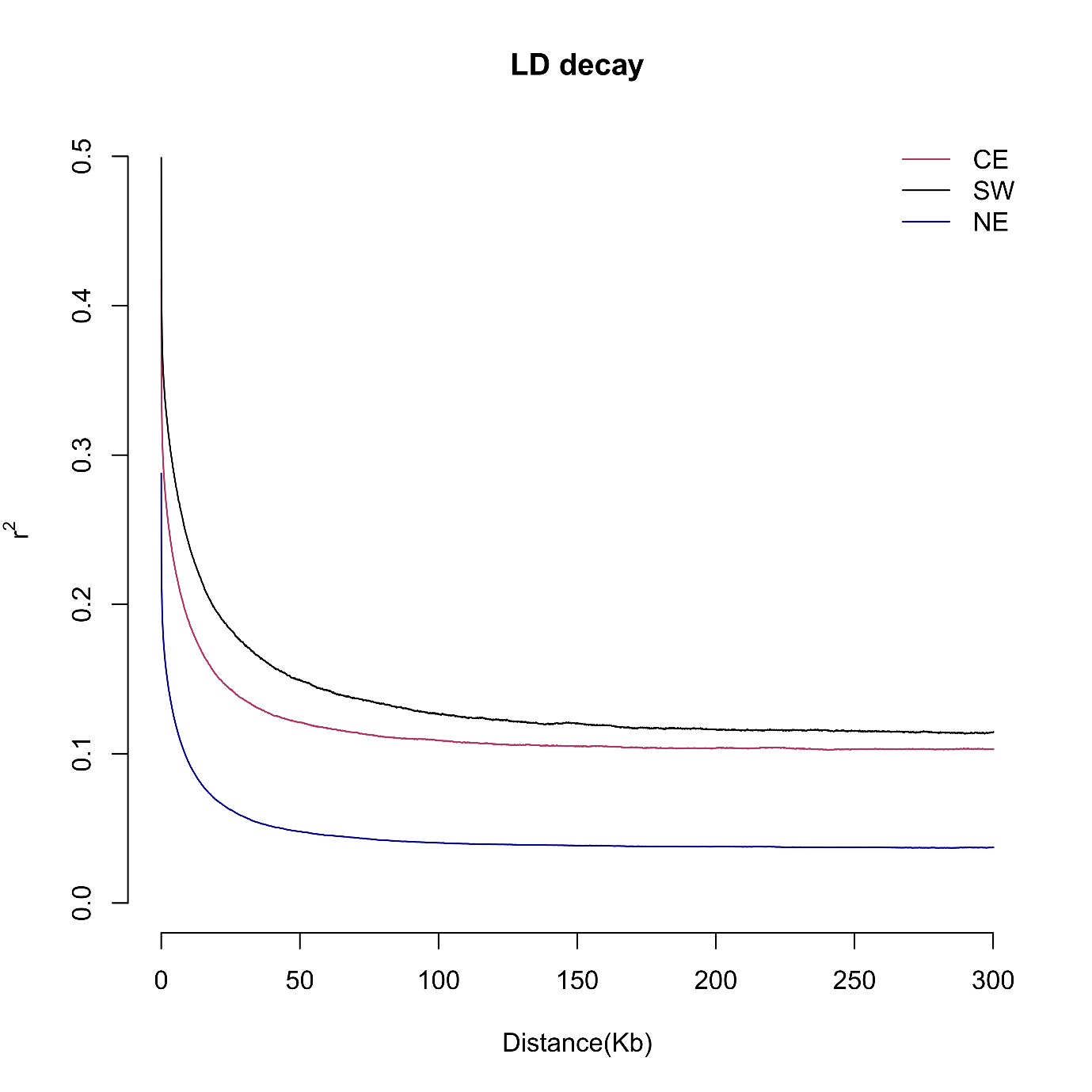

**Figure S13.** Genome-wide Fst (by SNP site) distribution between *B. platyphylla* populations, based on 1,387,994 SNPs and excluding admixed individuals. Populations’ assignment based on *fastSTRUCTURE* at K = 3, excluding admixed individuals (q < 0.9).

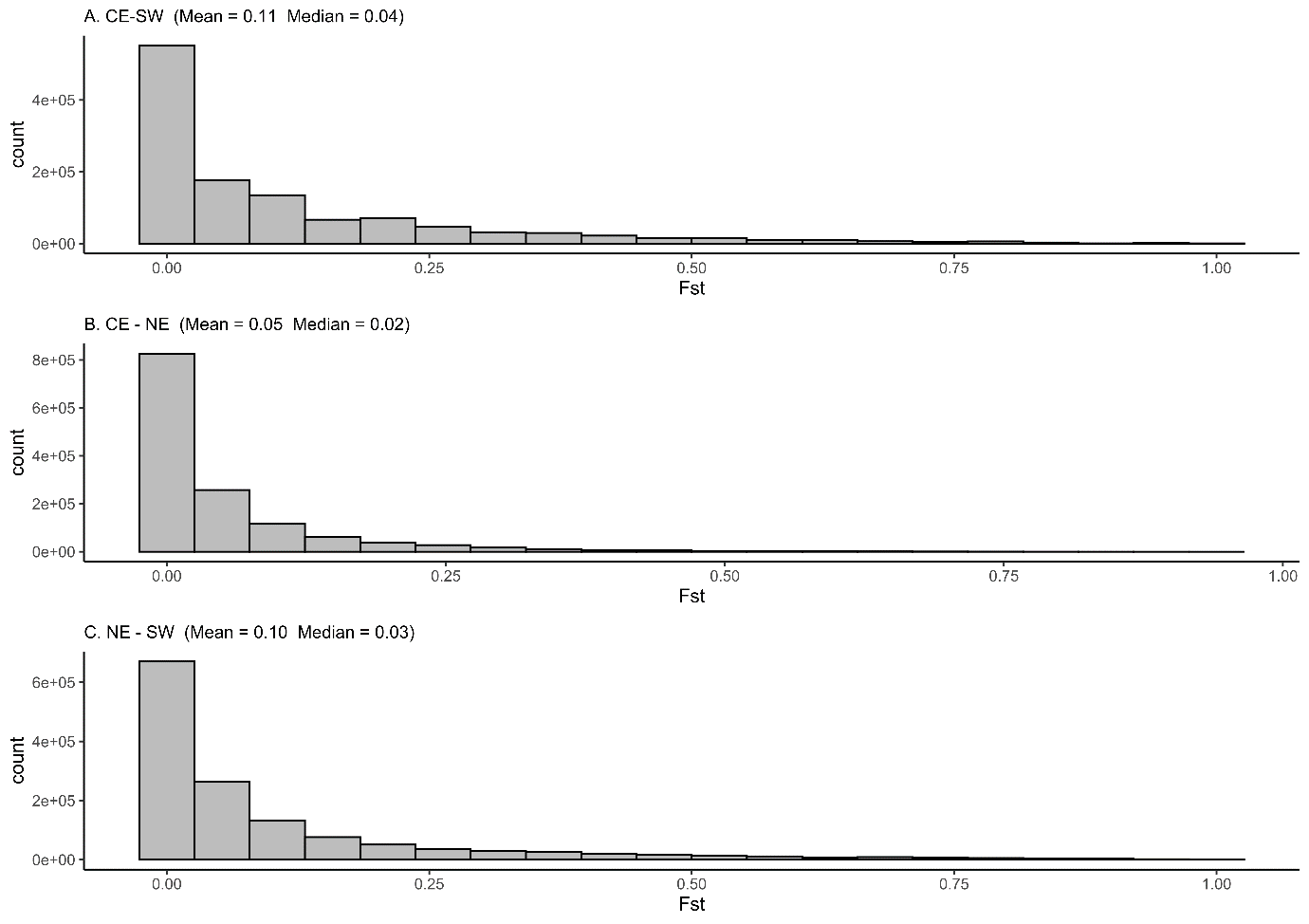

**Figure S14.** Pairwise nucleotide diversity π bar plots per population, computed in windows of 5,000 bp across the 14 *B. platyphylla* chromosomes. Populations according to *fastSTRUCTURE* at K = 3, excluding admixed individuals (q < 0.9).

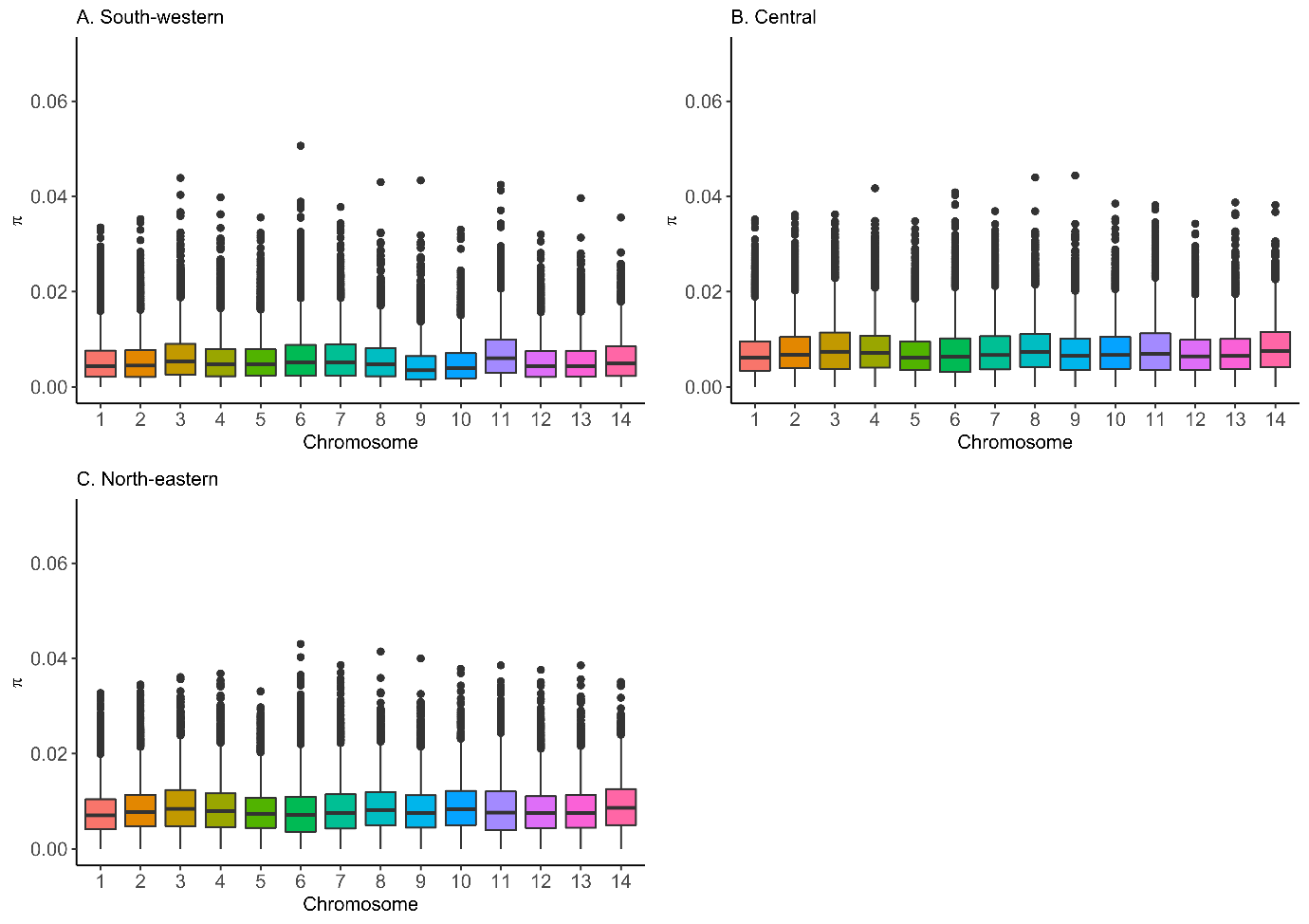

**Figure S15.** Pairwise nucleotide diversity π bar plots per population, computed in windows of 5,000 bp across the entire *B. platyphylla* genome. Populations according to *fastSTRUCTURE* at K = 3, excluding admixed individuals (q < 0.9).

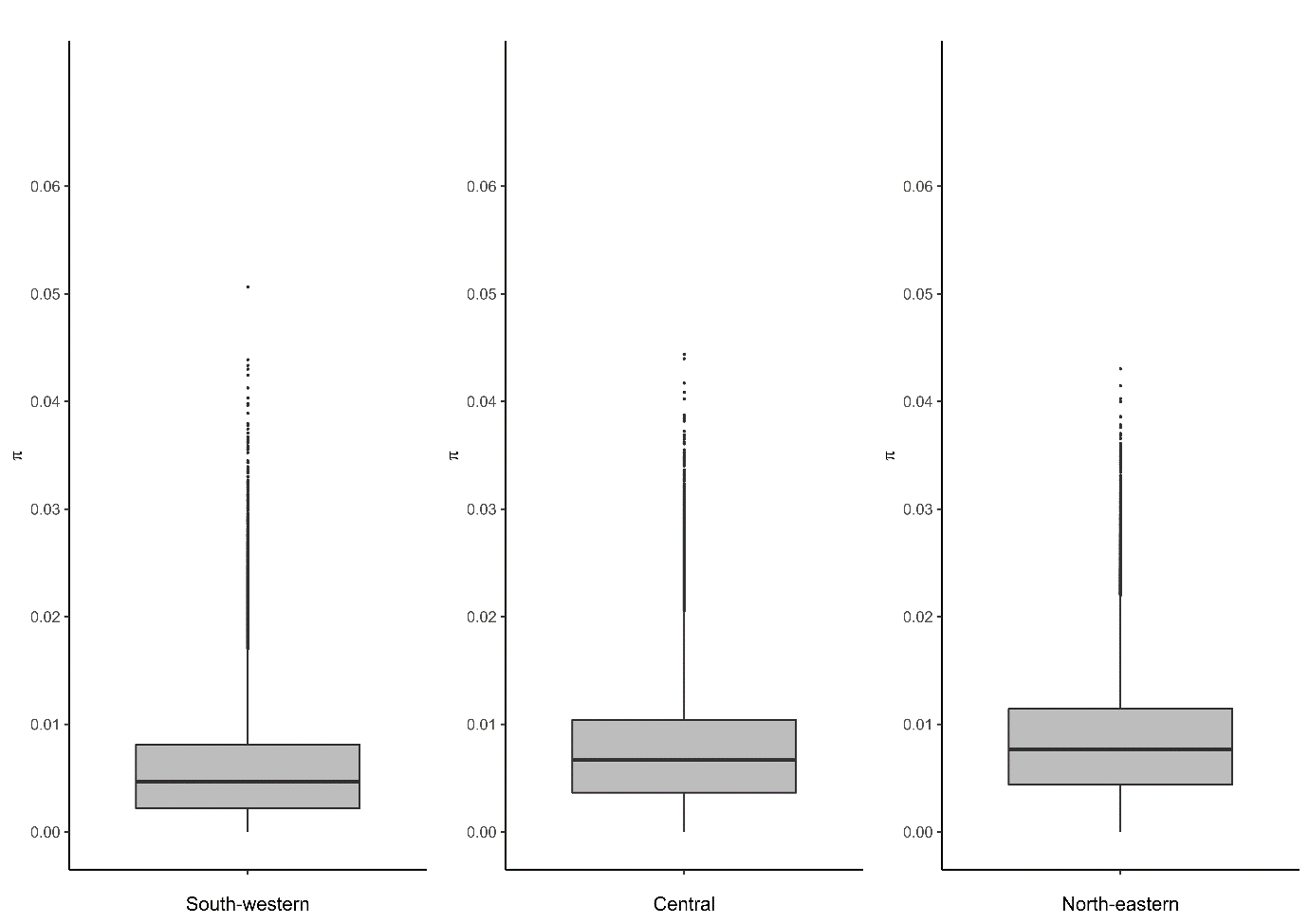

**Figure S16.** Jack-knife test of variable importance, using training gain for the Maxent model including 11 variables.

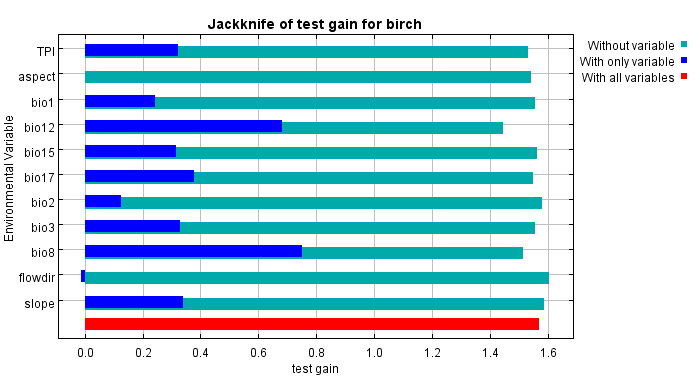

**Figure S17.** Jack-knife test of variable importance, using test gain instead of training gain for the Maxent model including 11 variables.

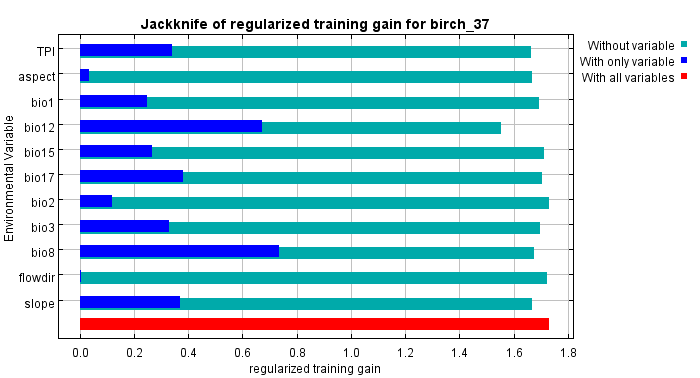

**Figure S18.** Jack-knife test of variable importance, using AUC on test data for the Maxent model including 11 variables.

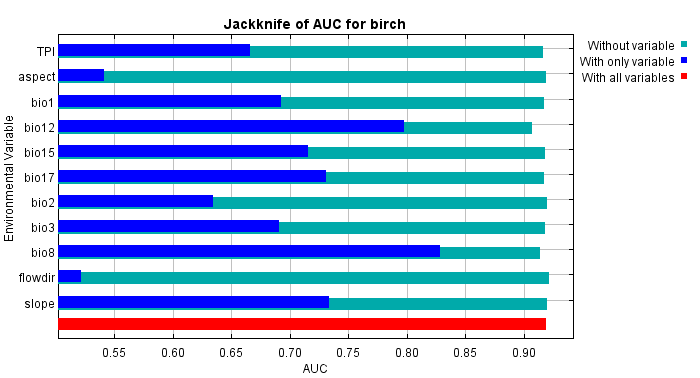

**Figure S19.** Omission rate and predicted area as a function of the cumulative threshold for the final Maxent model including nine variables, averaged over 50 replicate runs.

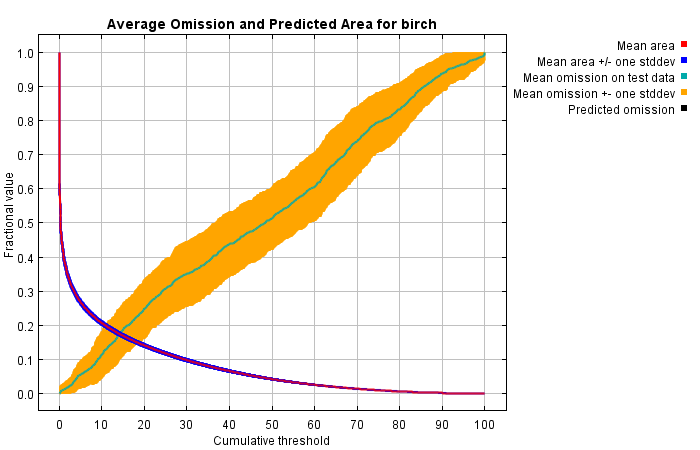

**Figure S20.** Receiver Operator Characteristic (ROC) curve for the final Maxent model including nine variables, averaged over 50 replicate runs. The average test AUC for the replicate runs is 0.913, and the standard deviation is 0.017.

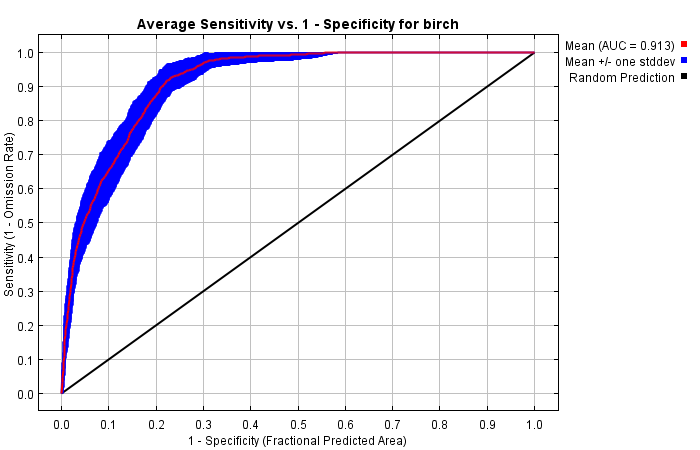

**Figure S21.** LFMM2 results. A) P-values Manhattan plot of the *LFMM2* analysis with K = 3. Green points are SNP with corresponding q-value (FDR) < 0.01. B) Histogram of p-values across environmental variables, suggesting that the rate of false positive is well controlled. The histogram of significance values is expected to be flat with a peak near 0. Total number of SNPs tested = 1,387,994.

A)

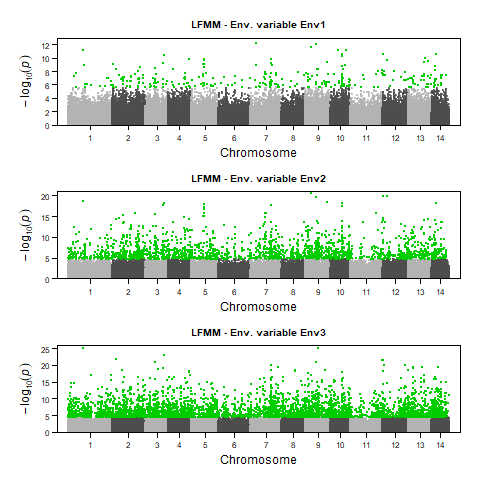

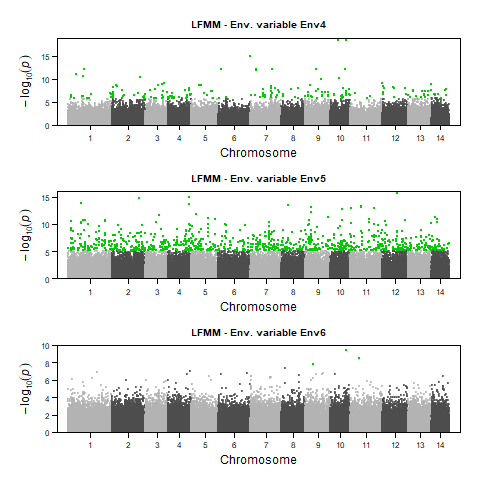

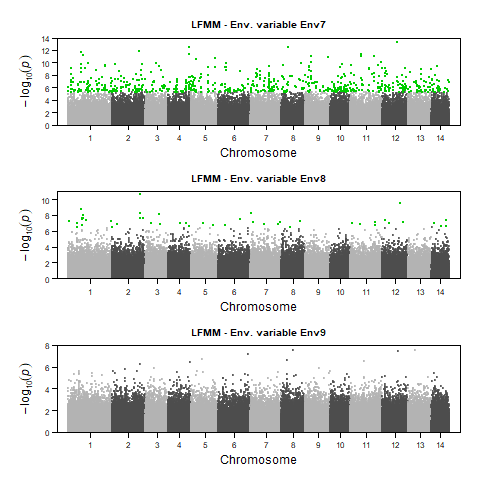

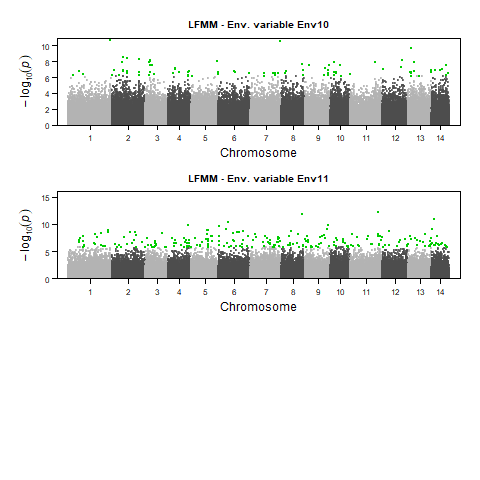

B)

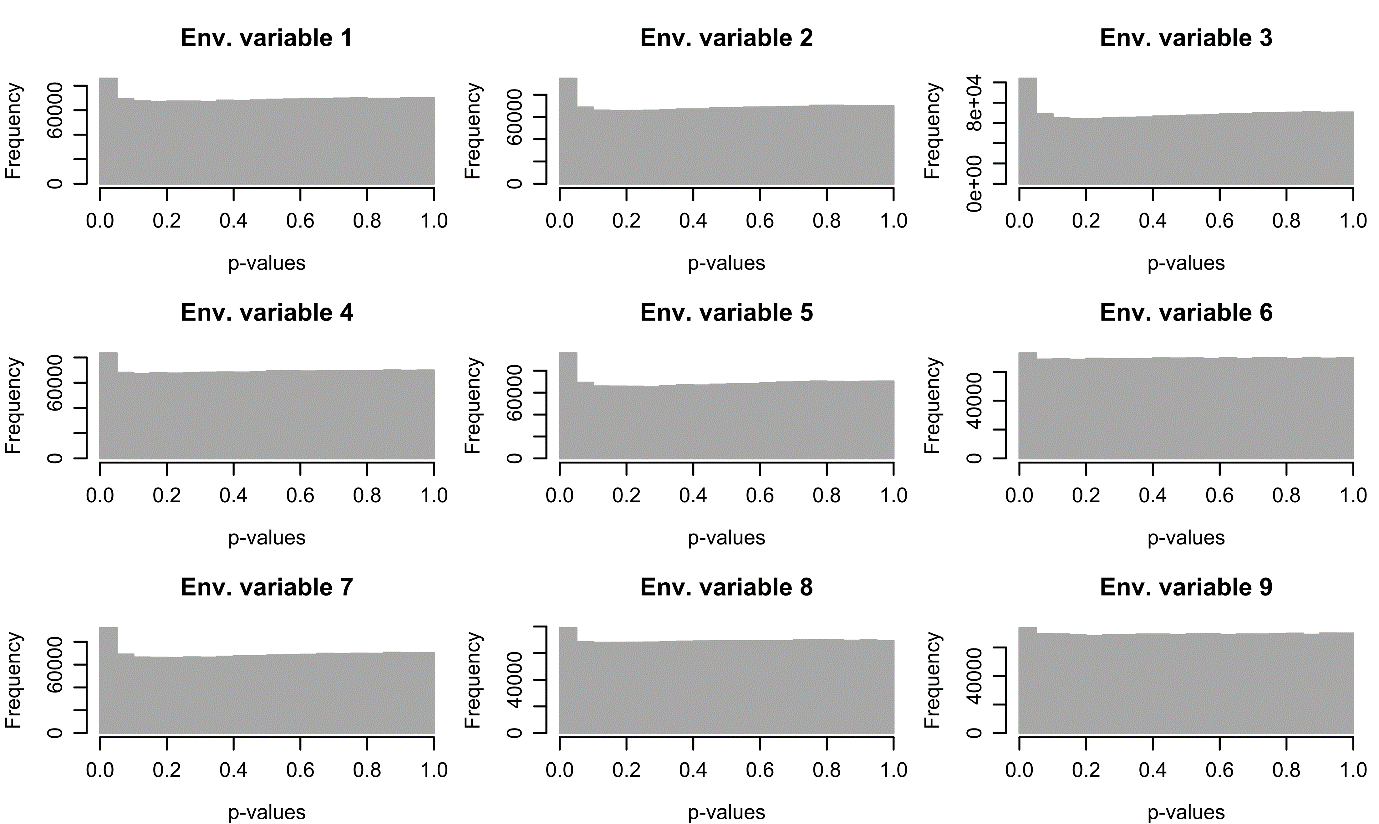

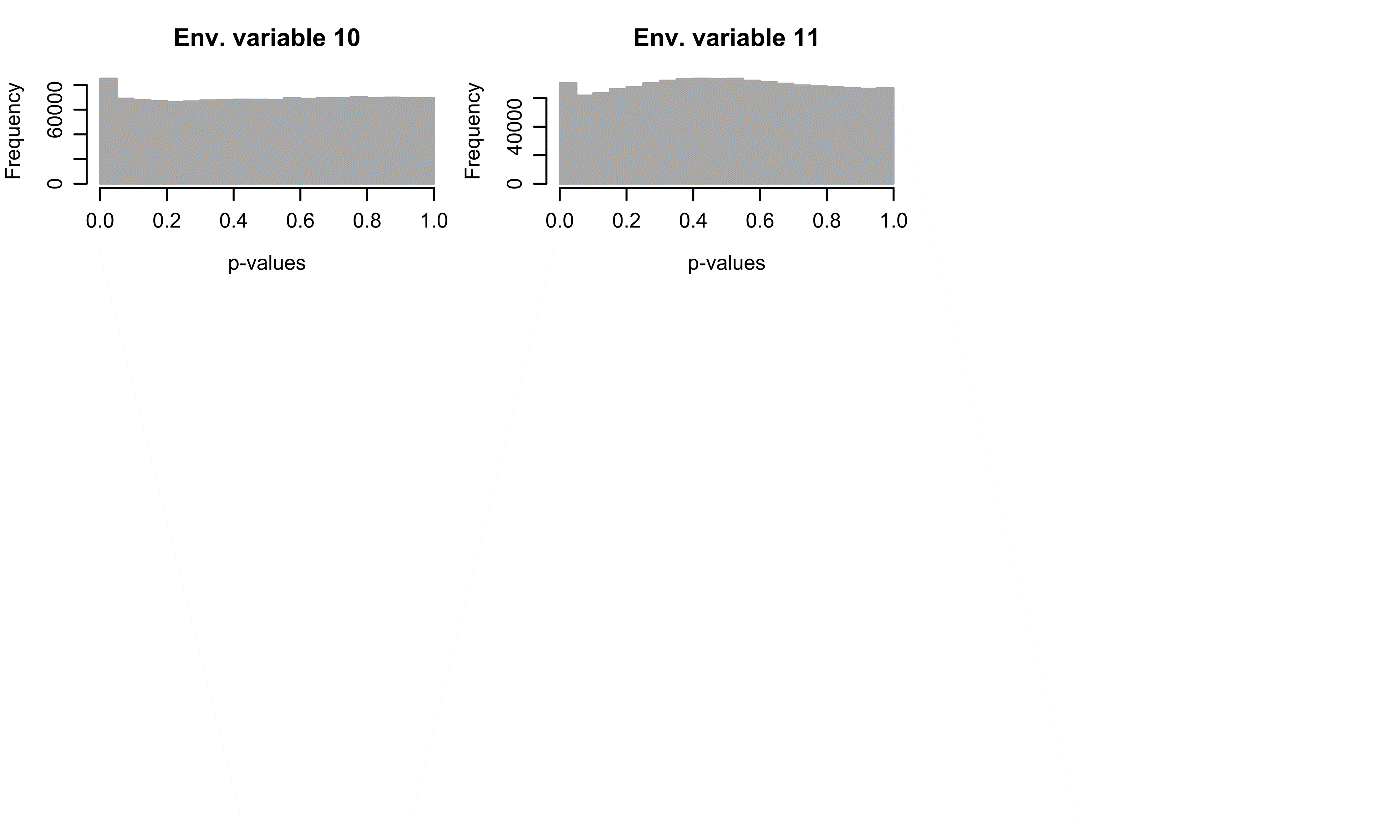

Figure S22. EAA results. **A)** Distribution of the 7,609 putatively adaptive SNPs identified with LFMM2 across the *B. pendula* reference genome. **B)** Distribution of the 11,304 SNP-environment associations detected (q < 0.01 in LFMM2) across environmental variables.

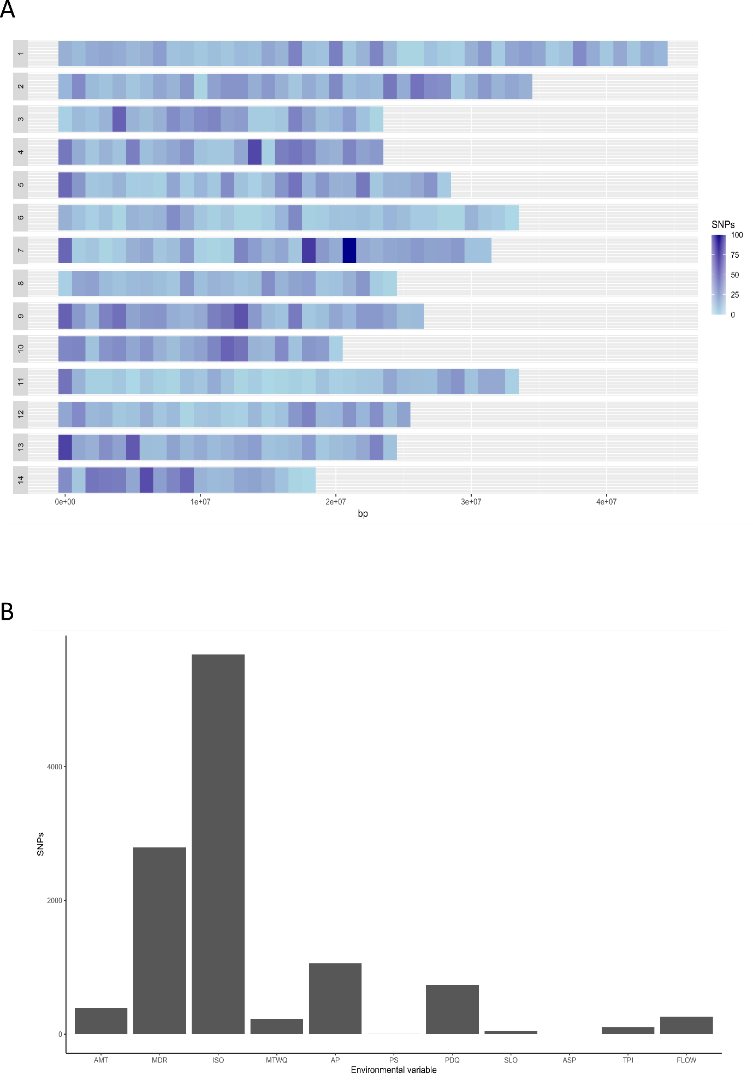

**Figure S23.** PCA of 71 *B. platyphylla* individuals based on the 7,609 putatively adaptive SNPs identified in this study. Colours represent fastSTRUCTURE populations assignments at K = 3. A) PC1 against PC2. B) PC1 against PC3. C) PC2 against PC3. D) Eigenvalues of the computed principal components.
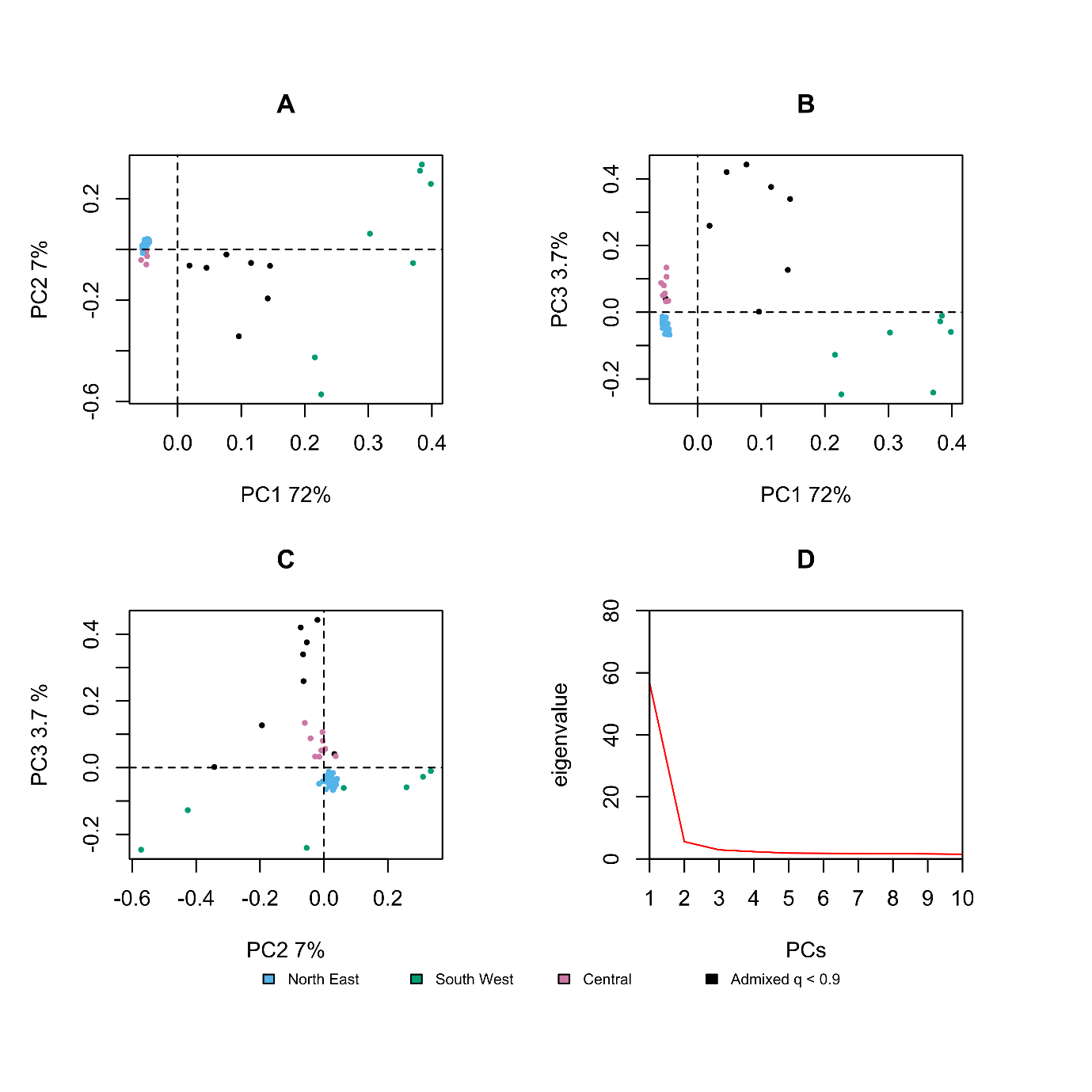

**Figure S24.** PCA of 71 *B. platyphylla* individuals based on 7,500 putatively “neutral” SNPs. Colours represent fastSTRUCTURE populations assignments at K = 3. A) PC1 against PC2. B) PC1 against PC3. C) PC2 against PC3. D) Eigenvalues of the computed principal components.
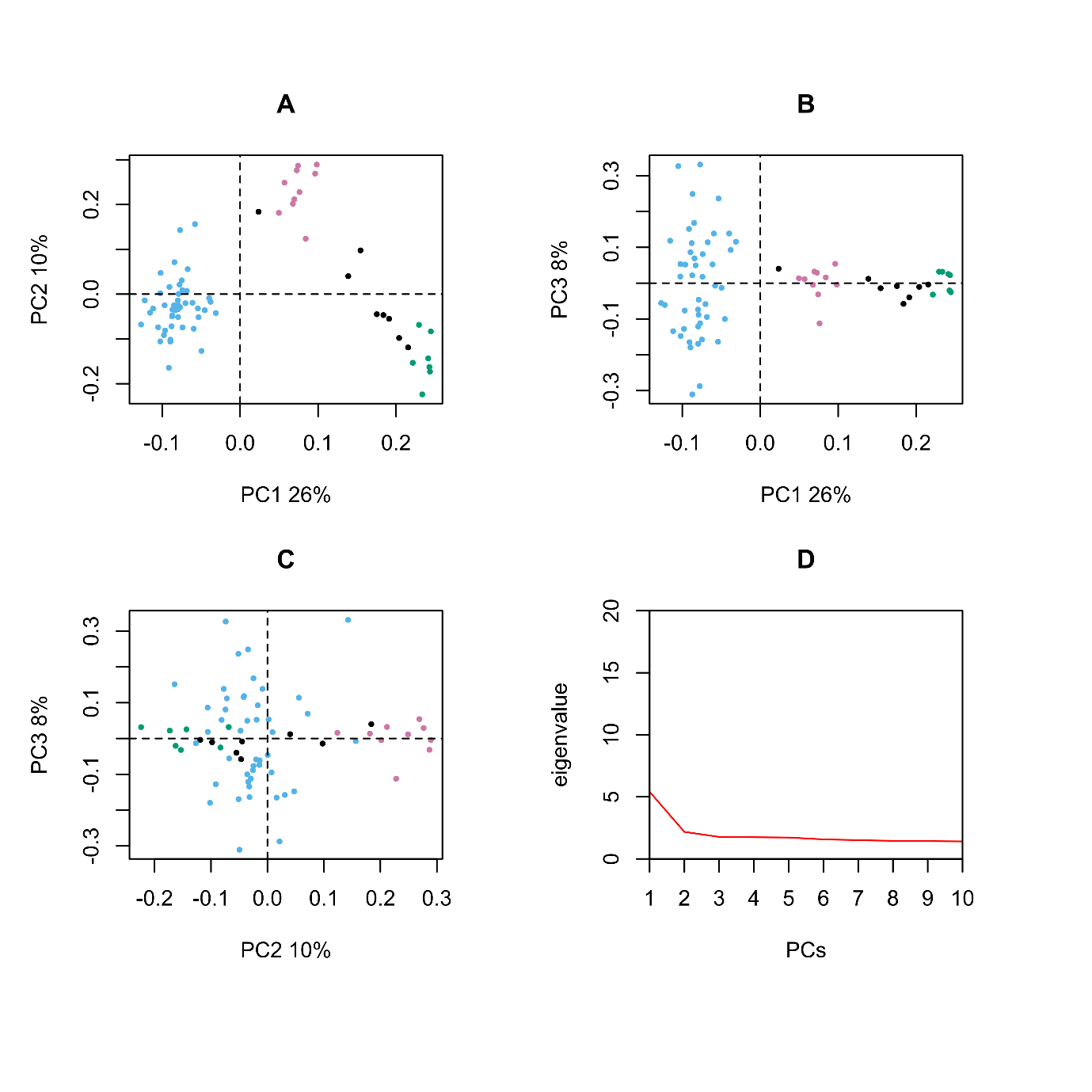

**Figure S25.** Fst (by SNP site) distribution between *B. platyphylla* populations, based on 7,609 adaptive SNPs and excluding admixed individuals. Populations’ assignment based on *fastSTRUCTURE* at K = 3, excluding admixed individuals (q < 0.9).

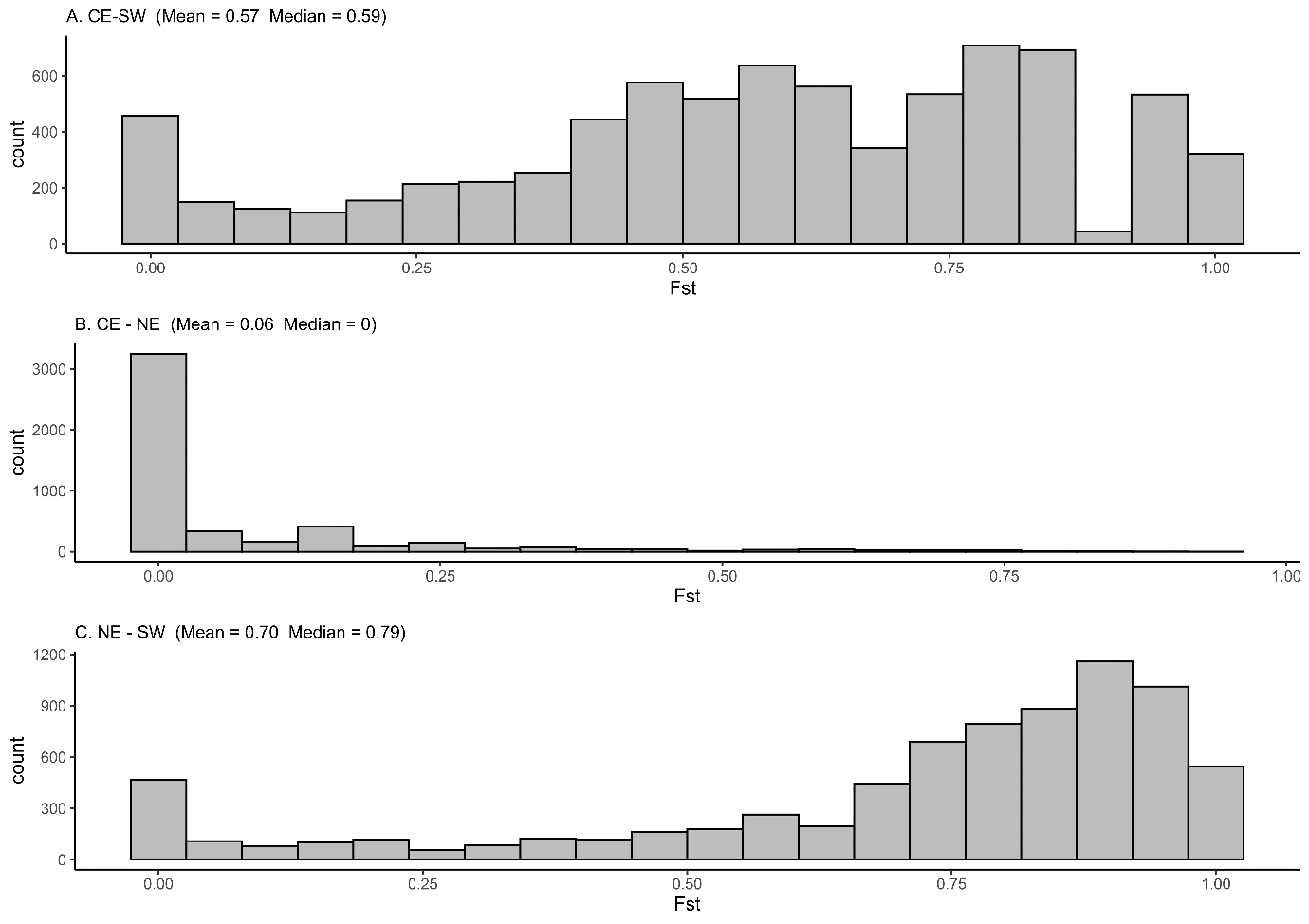

**Figure S26.** Fst (by SNP site) distribution between *B. platyphylla* populations, based on 7,500 neutral SNPs and excluding admixed individuals. Populations’ assignment based on *fastSTRUCTURE* at K = 3, excluding admixed individuals (q < 0.9).

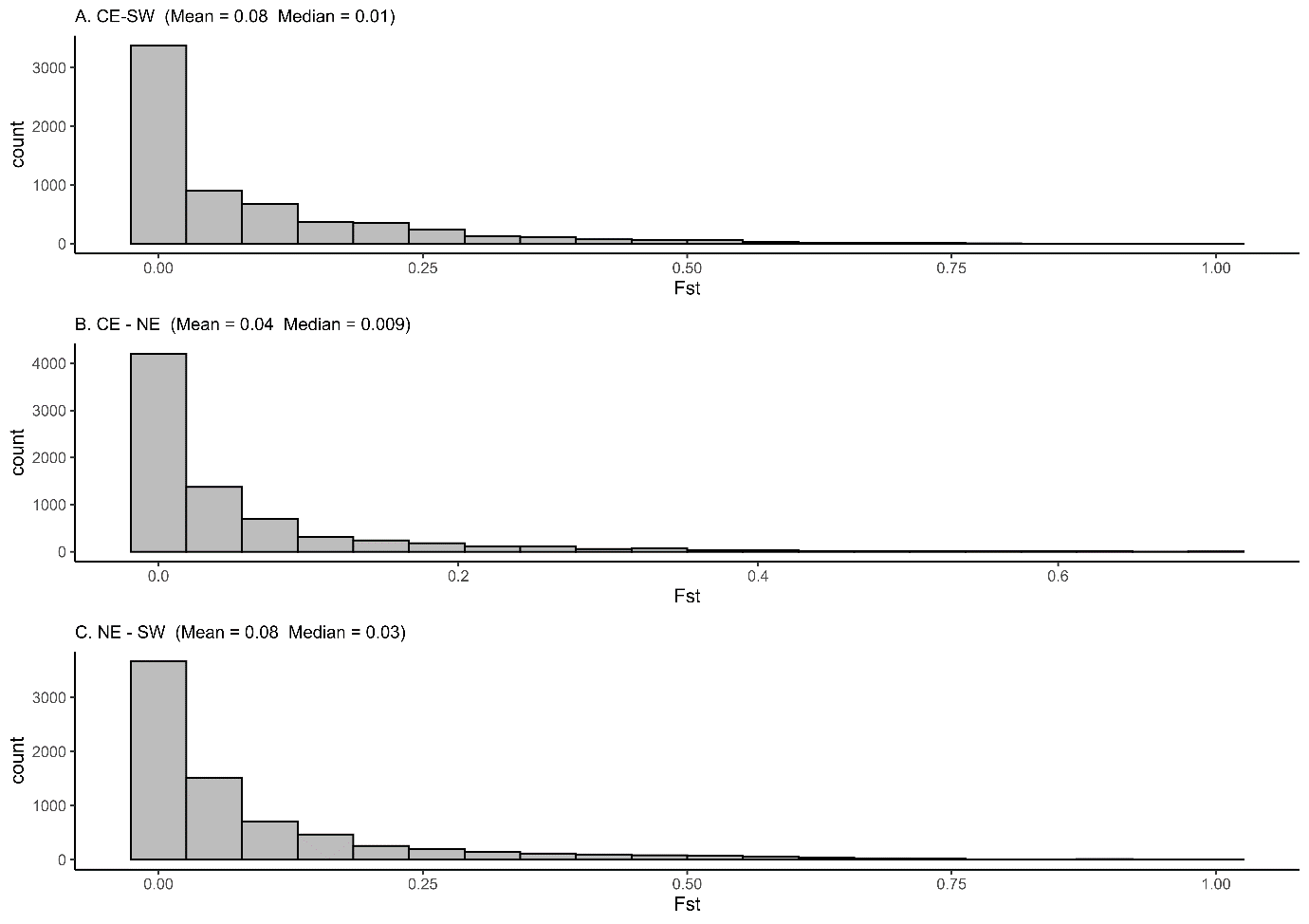

**Figure S27.** Omics Box functional enrichment analysis results (FDR < 5%) of the 1,633 genes spanned by the 7,609 adaptive SNPs identified with *LFMM2*. The plot includes all three ontology levels. The GOs retrieved for each gene region have been reduced to the most specific term in the gene ontology hierarchy.

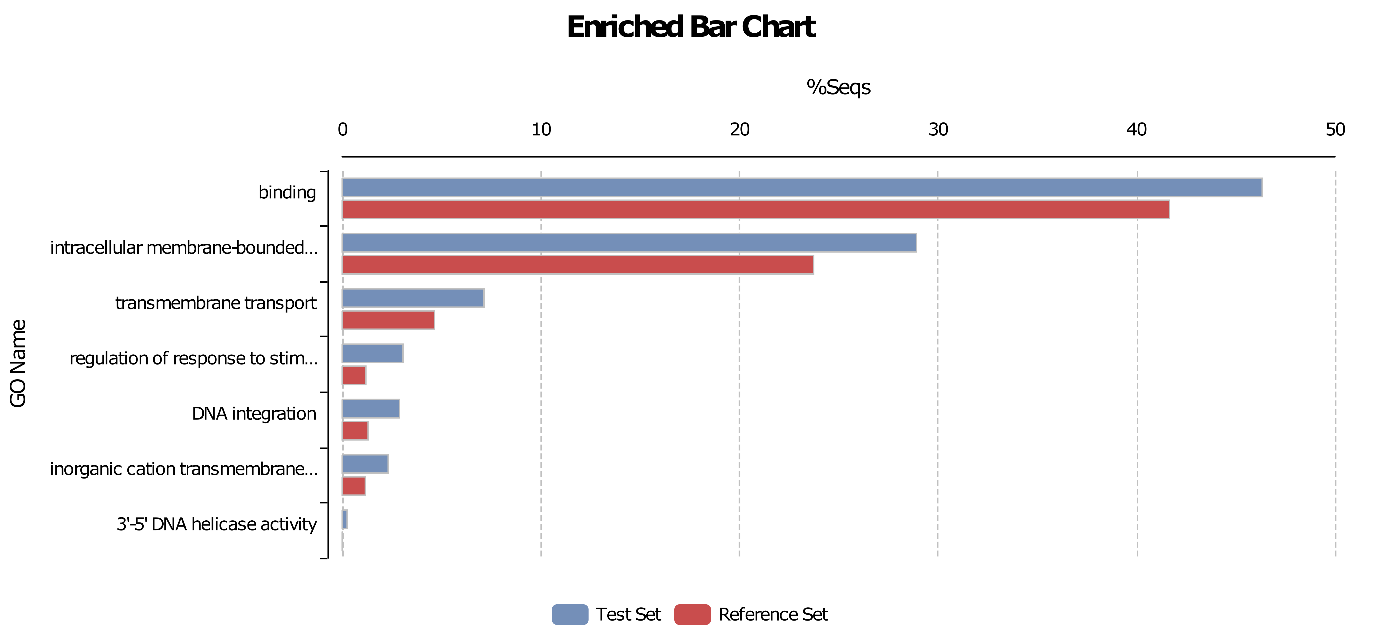

**Table S1.** Detailed information on sampling and resequencing.

| *Population code* | *Latitude* | *Longitude* | *Number of clean reads* | *Number of mapped reads* | *Percentage of mapped reads (%)* |
| --- | --- | --- | --- | --- | --- |
| AHT15 | 48.3777 | 85.7447 | 95108278 | 91773480 | 96.5 |
| AHT20 | 48.3795 | 85.7479 | 79444471 | 76620148 | 96.4 |
| ALE027 | 51.2818 | 121.4191 | 96472026 | 93032815 | 96.4 |
| ALS046 | 50.9 | 121.4212 | 78941821 | 76179630 | 96.5 |
| ALY036 | 51.5469 | 121.7336 | 75425352 | 72546335 | 96.2 |
| BEJ3 | 47.7249 | 86.9159 | 92422883 | 89176947 | 96.5 |
| BJC007 | 53.4901 | 122.3499 | 70706776 | 68192740 | 96.4 |
| BMX54 | 32.8011 | 100.8116 | 87728865 | 84442989 | 96.3 |
| BX54 | 30.8295 | 102.7403 | 82641706 | 79579298 | 96.3 |
| CGZ003 | 51.9907 | 124.617 | 86423518 | 82807447 | 95.8 |
| DCX5 | 28.6167 | 100.1922 | 74270014 | 71367883 | 96.1 |
| DFX11 | 31.1147 | 100.9644 | 80531220 | 74391328 | 92.4 |
| DQS13 | 41.0516 | 111.8089 | 83821719 | 80586874 | 96.1 |
| DTX94 | 37.186 | 101.5427 | 88645240 | 85119253 | 96.0 |
| EDG18 | 41.7171 | 126.4611 | 68634376 | 65886090 | 96.0 |
| EDG27 | 41.717 | 126.4618 | 84619385 | 77580250 | 91.7 |
| FHSJ1 | 44.2193 | 127.9786 | 93948831 | 90247913 | 96.1 |
| FYX7 | 47.2135 | 89.843 | 73588524 | 70957171 | 96.4 |
| HBH12 | 48.0726 | 86.342 | 86639208 | 83393478 | 96.3 |
| HBH2 | 48.0727 | 86.3415 | 93353556 | 89974372 | 96.4 |
| HGS18 | 47.1663 | 130.2832 | 86411628 | 83065783 | 96.1 |
| HLS53 | 42.4833 | 129.0528 | 83804019 | 81096539 | 96.8 |
| HNX7 | 46.3334 | 130.886 | 89061012 | 86065739 | 96.6 |
| HR6 | 41.2565 | 125.1582 | 134,129,672 | 129136963 | 96.3 |
| HTY15 | 41.8068 | 126.3612 | 74296931 | 71749626 | 96.6 |
| HX1 | 34.0296 | 105.9652 | 84099348 | 80988071 | 96.3 |
| JD24 | 29.2119 | 101.4489 | 79669091 | 75939795 | 95.3 |
| JS39 | 27.4391 | 99.8034 | 75823319 | 72834926 | 96.1 |
| JXS19 | 45.3755 | 130.9656 | 74515373 | 71741428 | 96.3 |
| JY23 | 48.6551 | 130.4574 | 92096901 | 88942754 | 96.6 |
| JYX28 | 42.292 | 126.7209 | 74378176 | 71889202 | 96.7 |
| KCL2 | 41.0696 | 125.0931 | 120,208,514 | 115,947,440 | 96.5 |
| KNS12 | 48.6767 | 87.014 | 81694447 | 77800819 | 95.2 |
| KNS4 | 48.6787 | 87.0135 | 78621478 | 75626395 | 96.2 |
| LBX31 | 47.7191 | 130.8627 | 73488854 | 70960134 | 96.6 |
| LJP1 | 27.1222 | 100.2572 | 74489677 | 69887415 | 93.8 |
| LLZ018 | 41.8415 | 126.5507 | 68977737 | 66455786 | 96.3 |
| LM049 | 52.8803 | 123.1261 | 95296931 | 91875251 | 96.4 |
| LMS033 | 48.3454 | 122.2915 | 69164718 | 66810351 | 96.6 |
| LS33 | 36.8358 | 111.9603 | 79043181 | 76301165 | 96.5 |
| LXJ13 | 31.6495 | 102.821 | 101980534 | 97912512 | 96.0 |
| LYC11 | 34.4317 | 110.4852 | 72037777 | 69435165 | 96.4 |
| MDJ15 | 44.5341 | 130.1711 | 80833056 | 77707495 | 96.1 |
| MDJ2 | 44.5343 | 130.1708 | 82740819 | 79510087 | 96.1 |
| MG089 | 52.4248 | 122.5113 | 80846449 | 77864358 | 96.3 |
| MH024 | 53.3786 | 122.2587 | 68798644 | 66348179 | 96.4 |
| MNS1 | 43.8274 | 86.0682 | 80978331 | 78011246 | 96.3 |
| MQX50 | 34.6597 | 100.6272 | 83453311 | 80557376 | 96.5 |
| MSZ1 | 47.9875 | 130.759 | 85739346 | 82621706 | 96.4 |
| PQG49 | 37.89 | 111.4306 | 78183522 | 75533247 | 96.6 |
| QHX14 | 46.6841 | 90.3548 | 95738066 | 92386580 | 96.5 |
| QSLC51 | 48.5909 | 129.8668 | 81216632 | 78177595 | 96.3 |
| RTX16 | 32.5793 | 101.0815 | 73673220 | 71079464 | 96.5 |
| SDX29 | 31.9529 | 100.9266 | 70954809 | 68184929 | 96.1 |
| SLS16 | 30.9248 | 102.3128 | 67,482,194 | 64922238 | 96.2 |
| SWP3 | 35.4288 | 111.9715 | 74598260 | 71796850 | 96.2 |
| TZZ103 | 36.9395 | 102.6141 | 80145267 | 77185210 | 96.3 |
| WCS21 | 44.6561 | 127.3484 | 85343984 | 82003059 | 96.1 |
| WCS6 | 44.6563 | 127.3468 | 97872192 | 94574167 | 96.6 |
| WEG19 | 43.6511 | 129.516 | 82097483 | 79232118 | 96.5 |
| WEG26 | 43.6511 | 129.516 | 71811997 | 69041972 | 96.1 |
| WLG024 | 52.6535 | 124.4441 | 67993333 | 65198125 | 95.9 |
| WT42 | 38.831 | 113.8389 | 83683234 | 80511518 | 96.2 |
| WTG28 | 41.9269 | 126.0903 | 92797096 | 89223554 | 96.1 |
| WY29 | 43.0257 | 129.5483 | 72726663 | 70308073 | 96.7 |
| WYQ5 | 48.0796 | 129.2231 | 78521085 | 75804535 | 96.5 |
| XEXL17 | 45.2187 | 82.0478 | 73114971 | 70656562 | 96.6 |
| XEXL2 | 45.2199 | 82.0454 | 85742705 | 82188619 | 95.9 |
| XFL14 | 42.5305 | 128.7583 | 98912997 | 94705944 | 95.7 |
| XFL2 | 42.5303 | 128.7587 | 89848329 | 86157227 | 95.9 |
| XGLLP21 | 27.8293 | 99.7441 | 70443682 | 67688473 | 96.1 |
| XGLLP7 | 27.8293 | 99.7441 | 78361504 | 74936652 | 95.6 |
| XHC004 | 49.9104 | 124.5655 | 102022698 | 98582755 | 96.6 |
| XHS10 | 42.8874 | 127.0478 | 80830112 | 77985804 | 96.5 |
| XMX5 | 37.2623 | 101.9801 | 92847104 | 89270911 | 96.1 |
| XXG025 | 47.8172 | 122.6181 | 88881295 | 85900807 | 96.6 |
| XYQ060 | 50.7067 | 124.3103 | 77914626 | 75171347 | 96.5 |
| YCS25 | 47.6333 | 128.5499 | 71287688 | 65225061 | 91.5 |
| YJA54 | 30.0472 | 101.2854 | 80072173 | 76742373 | 95.8 |
| YKS014 | 49.1213 | 120.9051 | 104383304 | 99384140 | 95.2 |
| YLK048 | 48.8535 | 121.6293 | 86034800 | 83023217 | 96.5 |
| ZJ027 | 52.9555 | 122.5723 | 66,016,635 | 63643574 | 96.4 |
| ZSL004 | 49.2359 | 124.6643 | 75063914 | 72555543 | 96.7 |

**Table S2.** The relative likelihood of the 28 demographic models of three populations.

| **Models** | **Max(log10 likelihood)** | **AIC** | **No. of parameters** | **deltaAIC** | **Model normalized relative likelihood** |
| --- | --- | --- | --- | --- | --- |
| **Model 01** | -399620268.40 | 1840319359.74 | 7 | 6812077.36 | 0.00 |
| **Model 02** | -398348024.99 | 1834460478.35 | 15 | 953195.97 | 0.00 |
| **Model 03** | -398373761.40 | 1834578994.86 | 13 | 1071712.48 | 0.00 |
| **Model 04** | -399617963.68 | 1840308752.10 | 10 | 6801469.72 | 0.00 |
| **Model 05** | -398177381.08 | 1833674640.07 | 18 | 167357.69 | 0.00 |
| **Model 06** | -398218519.50 | 1833864085.50 | 16 | 356803.12 | 0.00 |
| **Model 07** | -399607108.92 | 1840258766.12 | 11 | 6751483.74 | 0.00 |
| **Model 08** | -398220720.40 | 1833874227.01 | 19 | 366944.63 | 0.00 |
| **Model 09** | -398209251.32 | 1833821405.98 | 17 | 314123.60 | 0.00 |
| **Model 10** | -399467637.62 | 1839616485.05 | 15 | 6109202.67 | 0.00 |
| **Model 11** | -398187369.00 | 1833720646.14 | 23 | 213363.76 | 0.00 |
| **Model 12** | -398143298.68 | 1833517694.85 | 23 | 10412.47 | 0.00 |
| **Model 13** | -398238082.88 | 1833954188.21 | 21 | 446905.83 | 0.00 |
| **Model 14** | -398141038.51 | 1833507282.38 | 21 | 0.00 | 1.00 |
| **Model 15** | -399556629.10 | 1840026293.93 | 9 | 6519011.55 | 0.00 |
| **Model 16** | -398344242.05 | 1834443061.26 | 17 | 935778.88 | 0.00 |
| **Model 17** | -398336547.46 | 1834407622.33 | 15 | 900339.95 | 0.00 |
| **Model 18** | -399221295.64 | 1838482032.29 | 12 | 4974749.91 | 0.00 |
| **Model 19** | -398205814.43 | 1833805584.49 | 20 | 298302.11 | 0.00 |
| **Model 20** | -398182689.76 | 1833699087.47 | 18 | 191805.09 | 0.00 |
| **Model 21** | -399539311.63 | 1839946552.05 | 13 | 6439269.67 | 0.00 |
| **Model 22** | -398219319.44 | 1833867779.38 | 21 | 360497.00 | 0.00 |
| **Model 23** | -398255329.24 | 1834033606.60 | 19 | 526324.22 | 0.00 |
| **Model 24** | -399343917.88 | 1839046738.56 | 17 | 5539456.18 | 0.00 |
| **Model 25** | -398211255.25 | 1833830650.41 | 25 | 323368.03 | 0.00 |
| **Model 26** | -398184701.65 | 1833708366.54 | 25 | 201084.16 | 0.00 |
| **Model 27** | -398246605.07 | 1833993438.32 | 23 | 486155.94 | 0.00 |
| **Model 28** | -398141051.48 | 1833507346.10 | 23 | 63.72 | 1.46E-14 |

**Table S3.** The relative likelihood of the 9 demographic models of two populations

| **Models** | **Max(log10 likelihood)** | **AIC** | **No. of parameters** | **deltaAIC** | **Model normalized relative likelihood** |
| --- | --- | --- | --- | --- | --- |
| **Model 2pop 01** | -152984945.57 | 704521718.26 | 4 | 2724567.60 | 0.00 |
| **Model 2pop 02** | -152468174.34 | 702141902.78 | 6 | 344752.10 | 0.00 |
| **Model 2pop 03** | -152960700.28 | 704410068.54 | 6 | 2612917.80 | 0.00 |
| **Model 2pop 04** | -152484646.83 | 702217765.42 | 8 | 420614.70 | 0.00 |
| **Model 2pop 05** | -152932273.54 | 704279160.59 | 7 | 2482009.90 | 0.00 |
| **Model 2pop 06** | -152445257.30 | 702036371.93 | 9 | 239221.20 | 0.00 |
| **Model 2pop 07** | -152763616.92 | 703502474.13 | 10 | 1705323.40 | 0.00 |
| **Model 2pop 08** | -152402557.36 | 701839737.40 | 12 | 42586.70 | 0.00 |
| **Model 2pop 09** | -152393309.78 | 701797150.72 | 12 | 0.00 | 1.00 |

Table S4. The 26 climatic variables downloaded from [www.worldclim.org](http://www.worldclim.org) for 1970-2000. 11 uncorrelated variables (correlation coefficient < 0.7) were retained for the ENM and GEA.

| Climatic Variable ID | Description | Retained |
| --- | --- | --- |
| AMT | Annual Mean Temperature | Yes |
| MDR | Mean Diurnal Range (Mean of monthly (max temp - min temp)) | Yes |
| ISO | Isothermality (BIO2/BIO7) (×100) | Yes |
| TS | Temperature Seasonality (standard deviation ×100) | No |
| MaxTWM | Max Temperature of Warmest Month | No |
| MINTCM | Min Temperature of Coldest Month | No |
| TAR | Temperature Annual Range (BIO5-BIO6) | No |
| MTWQ | Mean Temperature of Wettest Quarter | Yes |
| MTDQ | Mean Temperature of Driest Quarter | No |
| MTWARMQ | Mean Temperature of Warmest Quarter | No |
| MTCOLDQ | Mean Temperature of Coldest Quarter | No |
| AP | Annual Precipitation | Yes |
| PWM | Precipitation of Wettest Month | No |
| PDM | Precipitation of Driest Month | No |
| PS | Precipitation Seasonality (Coefficient of Variation) | Yes |
| PWQ | Precipitation of Wettest Quarter | No |
| PDQ | Precipitation of Driest Quarter | Yes |
| PWARMQ | Precipitation of Warmest Quarter | No |
| PCOLDQ | Precipitation of Coldest Quarter | No |
| ALT | Elevation | No |
| SLO | Slope | Yes |
| ASP | Aspect | Yes |
| TPI | Topographic Position Index (TPI) | Yes |
| TRI | Terrain Ruggedness Index (TRI) | No |
| TR | Terrain Roughness | No |
| WFD | Water flow direction | Yes |

Table S5. The nine variables retained in the final maxent model. Percent contribution: in each iteration of training, the increase in regularized gain is added to the contribution of the corresponding variable or subtracted from it if the change to the absolute value of lambda is negative. Permutation importance: the values of each variable (in turn) on training presence and background data are randomly permuted. The model is then re-evaluated on the permuted data, and the resulting drop in training AUC is shown in the table, normalized to percentages.

| Variable | Percent contribution | Permutation importance |
| --- | --- | --- |
| AP | 34.4 | 31.3 |
| MTWQ | 21.9 | 17.6 |
| PDQ | 12 | 12.1 |
| SLO | 10.4 | 2.6 |
| TPI | 7.4 | 0.7 |
| ISO | 4.5 | 10.9 |
| MDR | 3.3 | 1.5 |
| AMT | 3.2 | 21.2 |
| PS | 3 | 2.2 |

**Table S6.** RONA results for each of the 71 *B. platyphylla* individual for seven environmental variables, under the future climate profile ssp370 at 2080-2100. The second row shows the number of SNPs identified associated with *LFMM2* and therefore included in the RONA calculation for each environmental variable.

| Variable | AMT | MDR | ISO | MTWQ | AP | PS | PDQ |
| --- | --- | --- | --- | --- | --- | --- | --- |
| SNPs | 391 | 2794 | 5679 | 227 | 1059 | 3 | 735 |
| ALE027 | 0.0644 | 0.0273 | 0.0144 | 0.277 | 0.0138 | 0.009 | 0.0126 |
| ALS046 | 0.0564 | 0.0205 | 0.0176 | 0.2569 | 0.0174 | 0.0406 | 0.0126 |
| ALY036 | 0.0558 | 0.0251 | 0.0039 | 0.2109 | 0.0123 | 0.0232 | 0.0063 |
| BJC007 | 0.0713 | 0.0116 | 0.0186 | 0.1656 | 0.0054 | 0.0034 | 0.0126 |
| BMX54 | 0.098 | 0.0319 | 0.0054 | 0.0239 | 0.0718 | 0.0026 | 0 |
| BX54 | 0.0923 | 0.0686 | 0.1573 | 0.096 | 0.0381 | 0.0968 | 0.0063 |
| CGZ003 | 0.1212 | 0.0162 | 0.0186 | 0.0666 | 0.0165 | 0.0595 | 0.0189 |
| DCX5 | 0.1061 | 0.0309 | 0.0271 | 0.1854 | 0.0163 | 0.2788 | 0.044 |
| DFX11 | 0.111 | 0.0395 | 0.0613 | 0.2209 | 0.0141 | 0.0597 | 0.0063 |
| DQS13 | 0.1931 | 0.0039 | 0.0363 | 0.2004 | 0.0267 | 0.3215 | 0.0126 |
| DTX94 | 0.0714 | 0.0159 | 0.0054 | 0.1264 | 0.024 | 0.0974 | 0.0126 |
| EDG18 | 0.2518 | 0.0093 | 0.0563 | 0.2356 | 0.0695 | 0.1824 | 0 |
| EDG27 | 0.2523 | 0.0093 | 0.0563 | 0.2332 | 0.0694 | 0.1824 | 0 |
| FHSJ1 | 0.2281 | 0.033 | 0.0152 | 0.239 | 0.0424 | 0.1606 | 0.0312 |
| HGS18 | 0.2464 | 0.016 | 0.0238 | 0.3724 | 0.0288 | 0.0325 | 0.0126 |
| HLS53 | 0.2335 | 0.014 | 0.0391 | 0.089 | 0.0484 | 0.1292 | 0.0126 |
| HNX7 | 0.2231 | 0.0336 | 0.0118 | 0.3261 | 0.0315 | 0.0094 | 0 |
| HR6 | 0.2276 | 0.0201 | 0.0517 | 0.2754 | 0.0271 | 0.1306 | 0.0345 |
| HTY15 | 0.2581 | 0.0076 | 0.051 | 0.3718 | 0.0645 | 0.1782 | 0 |
| HX1 | 0.1485 | 0.0098 | 0.0107 | 0.2984 | 0.0451 | 0.0414 | 0 |
| JD24 | 0.0738 | 0.0079 | 0.0488 | 0.1624 | 0.0098 | 0.0504 | 0 |
| JS39 | 0.1041 | 0.0063 | 0.0617 | 0.2531 | 0.0011 | 0.3347 | 0.0702 |
| JXS19 | 0.2457 | 0.0358 | 6e-04 | 0.3371 | 0.0267 | 0.0013 | 0 |
| JY23 | 0.2448 | 0.038 | 0.0651 | 0.3956 | 0.0076 | 0.0094 | 0.0063 |
| JYX28 | 0.2725 | 0.0083 | 0.0491 | 0.3257 | 0.0477 | 0.1913 | 0.0174 |
| KCL2 | 0.2121 | 0.0095 | 0.0611 | 0.3379 | 0.0212 | 0.0759 | 0.0172 |
| LBX31 | 0.2485 | 0.0116 | 0.0404 | 0.3515 | 0.0152 | 0.0478 | 0.0126 |
| LJP1 | 0.0951 | 0.0191 | 0.0308 | 0.2418 | 0.0054 | 0.3116 | 0.1259 |
| LLZ018 | 0.2834 | 0.0083 | 0.0511 | 0.2297 | 0.0528 | 0.2101 | 0.0294 |
| LM049 | 0.0439 | 0.0121 | 0.0126 | 0.0612 | 0.0106 | 0.0626 | 0.0251 |
| LMS033 | 0.2346 | 0.0037 | 0.0182 | 0.2406 | 0.0174 | 0.1257 | 0.0189 |
| LS33 | 0.2297 | 0.0015 | 0.016 | 0.3736 | 0.0038 | 0.1086 | 0.044 |
| LXJ13 | 0.0145 | 0.0096 | 0.0206 | 0.0469 | 0.0234 | 0.0358 | 0.0062 |
| LYC11 | 0.2093 | 0.0187 | 0.0322 | 0.3113 | 0.0348 | 0.0729 | 0.0616 |
| MDJ15 | 0.2721 | 0.0078 | 0.0244 | 0.325 | 0.0392 | 0.0761 | 0.0186 |
| MDJ2 | 0.2726 | 0.0078 | 0.0244 | 0.3263 | 0.0392 | 0.0761 | 0.0186 |
| MG089 | 0.0518 | 0.0017 | 0.0311 | 0.1053 | 0.0098 | 0.0356 | 0.0063 |
| MH024 | 0.0558 | 0.0157 | 0.0184 | 0.081 | 0.0021 | 0.0151 | 0 |
| MQX50 | 0.1198 | 0.0418 | 0.0604 | 0.189 | 0.0424 | 0.1299 | 0.0126 |
| MSZ1 | 0.2427 | 0.0085 | 0.054 | 0.3173 | 0.0256 | 0.0234 | 0.0063 |
| PQG49 | 0.1895 | 0.0093 | 0.0152 | 0.3765 | 0.0234 | 0.1692 | 0.0312 |
| QSLC51 | 0.2733 | 0.0157 | 0.0344 | 0.2716 | 0.0016 | 0.027 | 0.0251 |
| RTX16 | 0.0767 | 0.0351 | 0.0113 | 0.0044 | 0.0604 | 0.0153 | 0 |
| SDX29 | 0.0796 | 0.0396 | 0.0427 | 0.0153 | 0.0353 | 0.0436 | 0.0063 |
| SLS16 | 0.1267 | 0.0172 | 0.0373 | 0.1384 | 0.0539 | 0.0471 | 0.0565 |
| SWP3 | 0.2477 | 0.0069 | 0.0491 | 0.3891 | 0.0022 | 0.0322 | 0.0669 |
| TZZ103 | 0.1335 | 0.0222 | 0.0407 | 0.2946 | 0.0048 | 0.1358 | 0.0314 |
| WCS21 | 0.2496 | 0.0275 | 0.0114 | 0.374 | 0.0098 | 0.1328 | 0.0186 |
| WCS6 | 0.2498 | 0.0274 | 0.0114 | 0.3741 | 0.0098 | 0.1328 | 0.0186 |
| WEG19 | 0.2203 | 0.0193 | 0.0429 | 0.3058 | 0.0555 | 0.128 | 0.0126 |
| WEG26 | 0.2203 | 0.0193 | 0.0429 | 0.3067 | 0.0555 | 0.128 | 0.0126 |
| WLG024 | 0.1163 | 0.0199 | 0.0185 | 0.0856 | 0.0098 | 0.0218 | 0.0126 |
| WT42 | 0.2875 | 0.0039 | 0.0444 | 0.3282 | 0.0256 | 0.3431 | 0.063 |
| WTG28 | 0.2398 | 0.0142 | 0.049 | 0.2558 | 0.0745 | 0.1958 | 0 |
| WY29 | 0.2403 | 0.0124 | 0.0584 | 0.3272 | 0.0473 | 0.1136 | 0 |
| WYQ5 | 0.2511 | 0.0231 | 0.0214 | 0.2645 | 0.0103 | 0.0334 | 0.0189 |
| XFL14 | 0.1901 | 0.0339 | 0.0073 | 0.0544 | 0.0658 | 0.1758 | 0.0315 |
| XFL2 | 0.201 | 0.0337 | 0.0073 | 0.0529 | 0.0598 | 0.1758 | 0.0315 |
| XGLLP21 | 0.1147 | 0.041 | 0.0057 | 0.2543 | 0.0033 | 0.3226 | 0.0599 |
| XGLLP7 | 0.1147 | 0.0406 | 0.0057 | 0.2547 | 0.0033 | 0.1667 | 0.0598 |
| XHC004 | 0.245 | 0.0117 | 0.0233 | 0.201 | 0.0326 | 0.1132 | 0 |
| XHS10 | 0.2566 | 0.033 | 0.0182 | 0.3528 | 0.0549 | 0.1781 | 0 |
| XMX5 | 0.1059 | 0.0212 | 0.0102 | 0.1855 | 0.0041 | 0.1126 | 0.0188 |
| XXG025 | 0.2757 | 0.0113 | 0.0274 | 0.364 | 0.0104 | 0.1005 | 0.0189 |
| XYQ060 | 0.1896 | 0.0104 | 0.027 | 0.1329 | 0.0268 | 0.0582 | 0.0063 |
| YCS25 | 0.2582 | 0.0218 | 0.0272 | 0.3147 | 0.0185 | 0.0191 | 0.0126 |
| YJA54 | 0.141 | 0.0012 | 0.0232 | 0.259 | 0.025 | 0.0471 | 0.0251 |
| YKS014 | 0.1758 | 0.0083 | 0.001 | 0.0549 | 0.0191 | 0.1696 | 0.0496 |
| YLK048 | 0.1673 | 0.0123 | 0.0088 | 0.1623 | 0.0327 | 0.1448 | 0.0249 |
| ZJ027 | 0.0455 | 0.0181 | 0.0273 | 0.068 | 0.0078 | 0.0331 | 0.0189 |
| ZSL004 | 0.2739 | 0.0125 | 0.0178 | 0.3271 | 0.0349 | 0.0717 | 0.0063 |
| Mean | 0.1773 | 0.0189 | 0.0305 | 0.2314 | 0.0281 | 0.1073 | 0.0207 |
| Min r^2^ | 1e-04 | 0 | 0 | 1e-04 | 0 | 0.136 | 0 |
| Max r^2^ | 0.3956 | 0.2522 | 0.8405 | 0.5437 | 0.2643 | 0.1809 | 0.1398 |
| Average r^2^ | 0.1959 | 0.0157 | 0.459 | 0.1854 | 0.0977 | 0.1627 | 0.0214 |

**Table S7.** Weighted-mean RONA and maximum RONA of the 71 *B. platyphylla* individuals, relative to the future climate profile ssp370 in 2080-2100.

| ID | mean RONA | max RONA |
| --- | --- | --- |
| ALE027 | 0.086 | 0.277 |
| ALS046 | 0.0829 | 0.2569 |
| ALY036 | 0.0664 | 0.2109 |
| BJC007 | 0.0525 | 0.1656 |
| BMX54 | 0.0418 | 0.098 |
| BX54 | 0.0609 | 0.1573 |
| CGZ003 | 0.0359 | 0.1212 |
| DCX5 | 0.0796 | 0.2788 |
| DFX11 | 0.0771 | 0.2209 |
| DQS13 | 0.0877 | 0.3215 |
| DTX94 | 0.0528 | 0.1264 |
| EDG18 | 0.1117 | 0.2518 |
| EDG27 | 0.111 | 0.2523 |
| FHSJ1 | 0.1028 | 0.239 |
| HGS18 | 0.1257 | 0.3724 |
| HLS53 | 0.0622 | 0.2335 |
| HNX7 | 0.111 | 0.3261 |
| HR6 | 0.1069 | 0.2754 |
| HTY15 | 0.1455 | 0.3718 |
| HX1 | 0.1065 | 0.2984 |
| JD24 | 0.055 | 0.1624 |
| JS39 | 0.0979 | 0.3347 |
| JXS19 | 0.1119 | 0.3371 |
| JY23 | 0.1243 | 0.3956 |
| JYX28 | 0.1297 | 0.3257 |
| KCL2 | 0.116 | 0.3379 |
| LBX31 | 0.1158 | 0.3515 |
| LJP1 | 0.1025 | 0.3116 |
| LLZ018 | 0.1093 | 0.2834 |
| LM049 | 0.0295 | 0.0626 |
| LMS033 | 0.0889 | 0.2406 |
| LS33 | 0.1212 | 0.3736 |
| LXJ13 | 0.0265 | 0.0469 |
| LYC11 | 0.1197 | 0.3113 |
| MDJ15 | 0.1206 | 0.325 |
| MDJ2 | 0.1209 | 0.3263 |
| MG089 | 0.0381 | 0.1053 |
| MH024 | 0.0268 | 0.081 |
| MQX50 | 0.0842 | 0.189 |
| MSZ1 | 0.1096 | 0.3173 |
| PQG49 | 0.1293 | 0.3765 |
| QSLC51 | 0.0907 | 0.2733 |
| RTX16 | 0.032 | 0.0767 |
| SDX29 | 0.0284 | 0.0796 |
| SLS16 | 0.077 | 0.1384 |
| SWP3 | 0.128 | 0.3891 |
| TZZ103 | 0.0983 | 0.2946 |
| WCS21 | 0.1226 | 0.374 |
| WCS6 | 0.1226 | 0.3741 |
| WEG19 | 0.1227 | 0.3058 |
| WEG26 | 0.123 | 0.3067 |
| WLG024 | 0.0358 | 0.1163 |
| WT42 | 0.1335 | 0.3431 |
| WTG28 | 0.1189 | 0.2558 |
| WY29 | 0.124 | 0.3272 |
| WYQ5 | 0.0905 | 0.2645 |
| XFL14 | 0.0621 | 0.1901 |
| XFL2 | 0.0596 | 0.201 |
| XGLLP21 | 0.096 | 0.3226 |
| XGLLP7 | 0.0903 | 0.2547 |
| XHC004 | 0.0825 | 0.245 |
| XHS10 | 0.1356 | 0.3528 |
| XMX5 | 0.0635 | 0.1855 |
| XXG025 | 0.1203 | 0.364 |
| XYQ060 | 0.0589 | 0.1896 |
| YCS25 | 0.1064 | 0.3147 |
| YJA54 | 0.0915 | 0.259 |
| YKS014 | 0.0432 | 0.1758 |
| YLK048 | 0.0732 | 0.1673 |
| ZJ027 | 0.0293 | 0.068 |
| ZSL004 | 0.1173 | 0.3271 |
| Mean | 0.0896 | 0.2533 |
| Max | 0.1455 | 0.3956 |

**Table S8.** RONA of each population for the seven environmental variables tested, under ssp370 at 2080-2100. Each population RONA was calculated by averaging the RONA of the individuals belonging to that population, according to *fastSTRUCTURE* at K = 3 and excluding admixed individuals.

|  | Central | South-western | North-eastern | All |
| --- | --- | --- | --- | --- |
| AMT | 0.17 | 0.10 | 0.20 | 0. 17 |
| MDR | 0.01 | 0.02 | 0.01 | 0. 01 |
| ISO | 0.02 | 0.02 | 0.02 | 0. 03 |
| MTWQ | 0.28 | 0.23 | 0.24 | 0. 23 |
| AP | 0.02 | 0.009 | 0.03 | 0. 02 |
| PS | 0.12 | 0.21 | 0.09 | 0. 10 |
| PDQ | 0.03 | 0.05 | 0.01 | 0. 02 |

### Final Maxent model response curves

There are two plots for each variable included in the final Maxent model. The first plots show how the predicted probability of presence changes as each environmental variable is varied, keeping all other environmental variables at their average sample value. The second plots show the predicted probability of presence in a Maxent model created using only the corresponding variable.

**Figure S28. AMT response curves.**

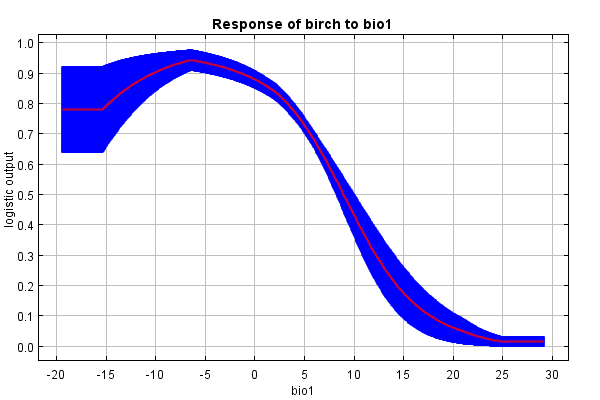

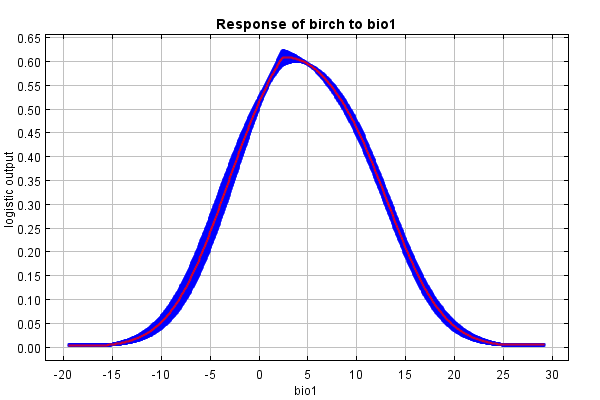

**Figure S29. MDR response curves.**

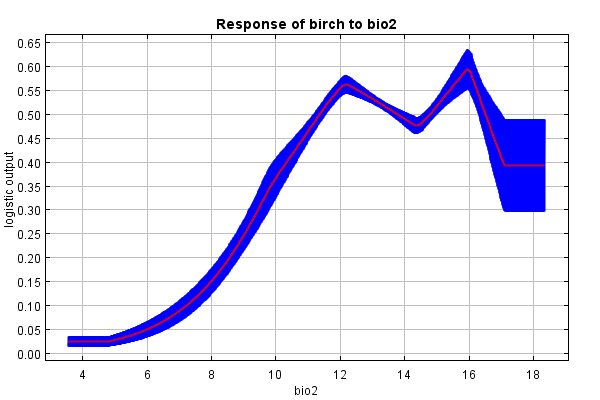

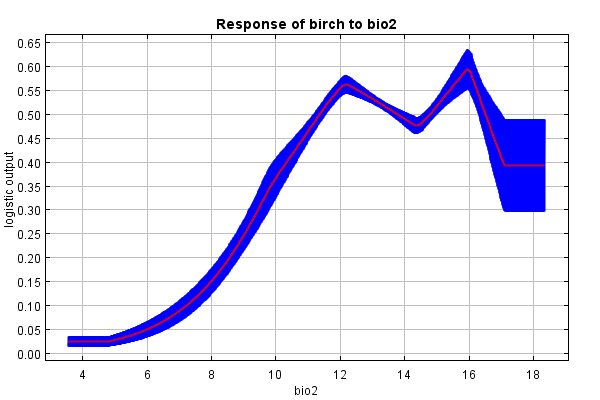

**Figure S30. ISO response curves.**

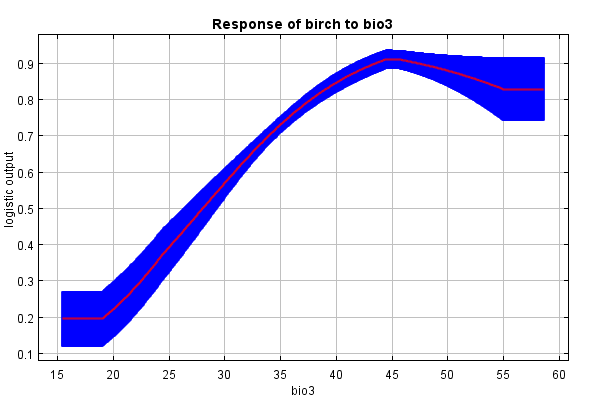

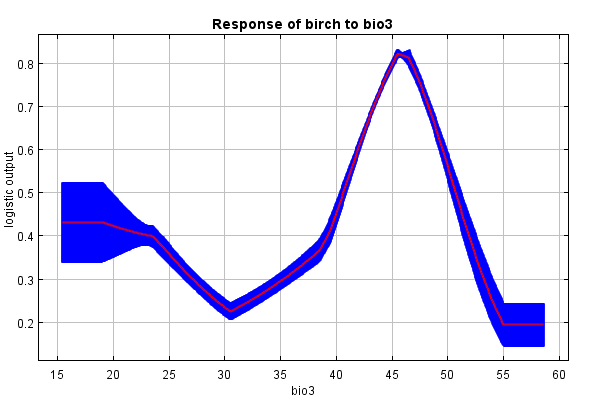

**Figure S31. MTWQ response curves.**

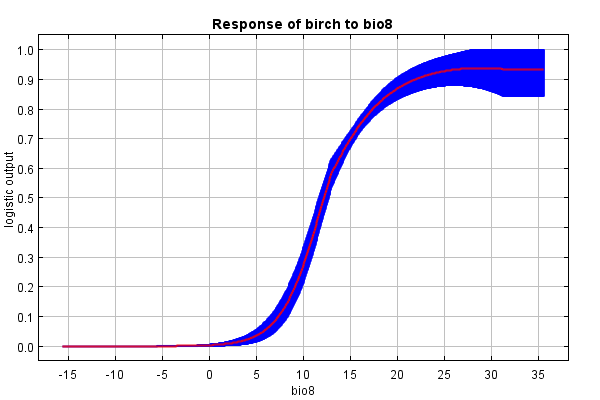

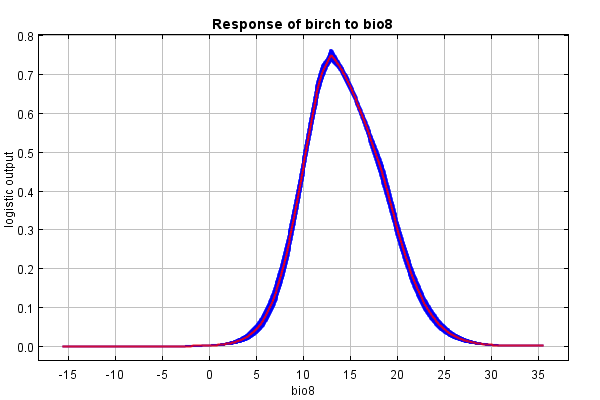

**Figure S32. AP response curves.**

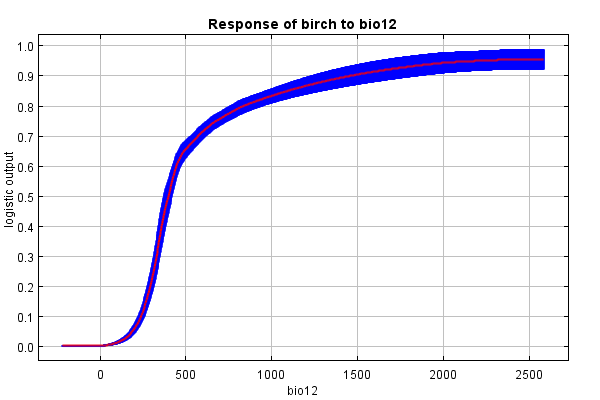

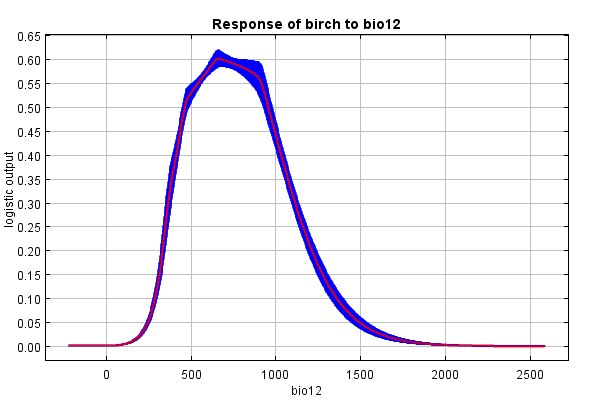

**Figure S33. PS response curves.**

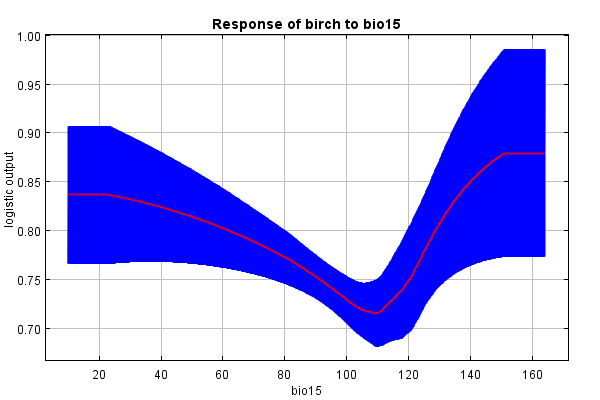

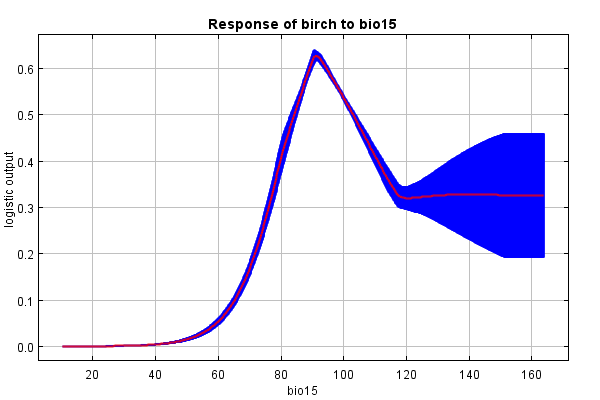

**Figure S34. PDQ response curves.**

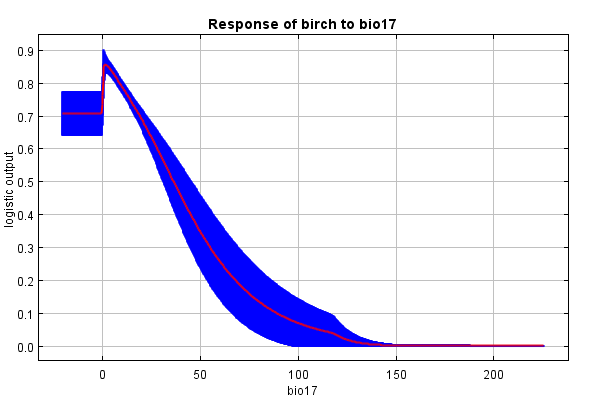

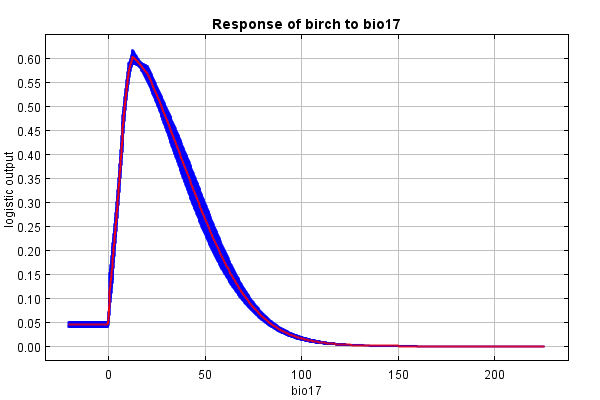

**Figure S35. SLOPE response curves.**

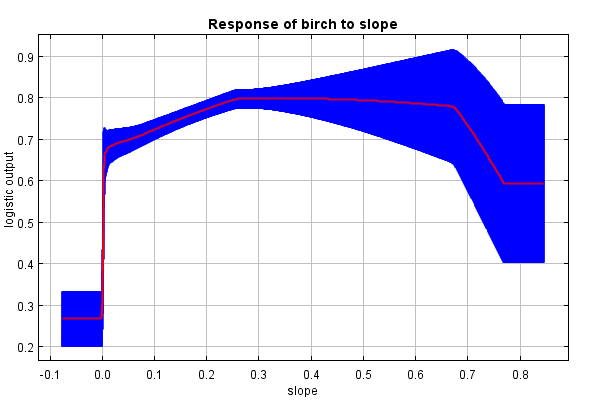

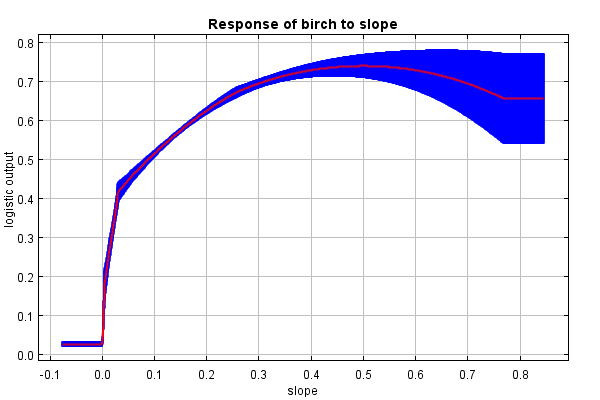

**Figure S36. TPI response curves.**

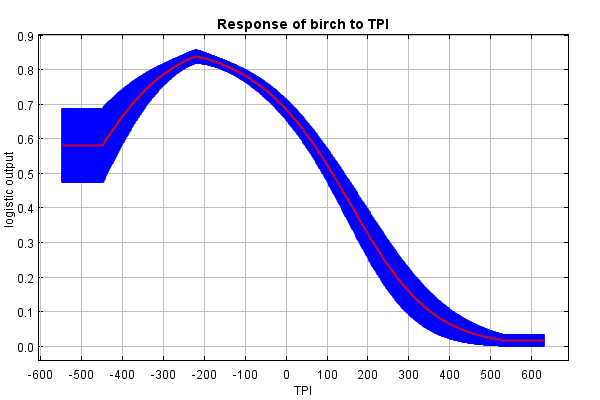

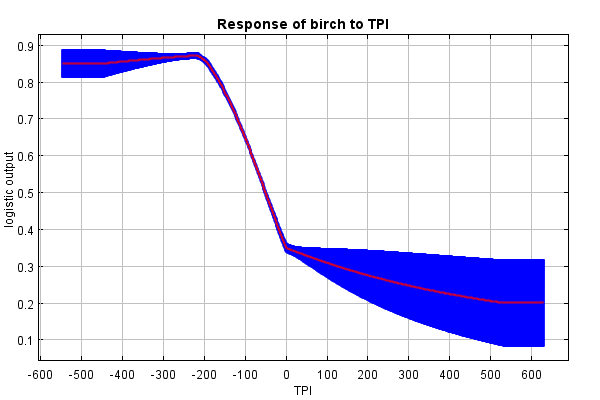
